# *Arabidopsis* TRM5 encodes a nuclear-localised bifunctional tRNA guanine and inosine-N1-methyltransferase that is important for growth

**DOI:** 10.1101/485516

**Authors:** Q. Guo, PQ. Ng, S. Shi, D. Fan, J. Li, H. Wang, T. Do, R. David, P. Mittal, R. Bock, M. Zhao, W. Zhou, I. R. Searle

## Abstract

Modified nucleosides in tRNAs are critical for protein translation. N^1^-methylguanosine-37 and N^1^-methylinosine-37 in tRNAs, both located at the 3’-adjacent to the anticodon, are formed by Trm5 and here we describe *Arabidopsis thaliana AtTrm5* (At3g56120) as a Trm5 ortholog. We show that *AtTrm5* complements the yeast *trm5* mutant, and *in vitro* methylates tRNA guanosine-37 to produce N^1^-methylguanosine (m^1^G). We also show *in vitro* that AtTRM5 methylates tRNA inosine-37 to produce N^1^-methylinosine (m^1^I) and in *Attrm5* mutant plants, we show a reduction of both N^1^-methylguanosine and N^1^-methylinosine. We also show that AtTRM5 is localized to the nucleus in plant cells. *Attrm5* mutant plants have overall slower growth as observed by slower leaf initiation rate, delayed flowering and reduced primary root length. In *Attrm5* mutants, mRNAs of flowering time genes are less abundant and correlated with delayed flowering. Finally, proteomics data show that photosynthetic protein abundance is affected in mutant plants. Our findings highlight the bifunctionality of AtTRM5 and the importance of the post-transcriptional tRNA modifications m^1^G and m^1^I at tRNA position 37 in general plant growth and development.

## Introduction

RNA has over 100 different post-transcriptional modifications that have been identified in organisms across all three domains of life (1–5). While several RNA modifications have been recently identified on mRNAs in plants and animals, tRNAs are still thought to be the most extensively modified cellular RNAs (6–9). These tRNA modifications are introduced at the post-transcriptional level by specific enzymes. These enzymes recognize polynucleotide substrates and modify individual nucleotide residues at highly specific sites. Some tRNA modifications have been shown to have a clear biological and molecular function (10, 11). Several tRNA modifications around the anticodon have been demonstrated to have crucial functions in translation, for example, by enhancing decoding (12), influencing the propensity to ribosomal frameshifting or facilitating wobbling (13–15). Modifications distal to the tRNA anticodon loop can also directly influence the tRNA recognition and/or translation process (16) or can have roles in tRNA folding and stability (1, 17). However, the precise functions of many tRNA modifications still remain unknown despite often being conserved across species. Often, loss of a tRNA modification does not negatively impair cell growth or cell viability under standard laboratory growth conditions (18). However, under environmental stress, such mutants display a discernible phenotype (10).

The tRNA anticodon loop position 37 is important to maintain translational fidelity and efficiency (11, 19) and almost all tRNAs are modified at this site. In bacteria, N^1^-methylation of guanosine at position 37, m^1^G37, is performed by TrmD-type enzymes (20). In Archaea and Eukaryotes, the m^1^G37 modification is enzymatically performed by functionally and evolutionarily unrelated Trm5-type proteins (21). In yeast *Saccharomyces cerevisiae*, m^1^G37 methylation of cytoplasmic and mitochondrial tRNAs is performed by Trm5p and complete loss of function mutants are lethal (22, 23). In humans, TRMT5 (tRNA methyltransferase 5), catalyzes the formation of m^1^G37 *in vivo*, and mitochondrial tRNA^PRO^ and tRNA^LEU^ have been found to also contain m^1^G37 (24, 25). Mutations in human *TRMT5* cause patients to have multiple respiratory-chain deficiencies and a reduction in mitochondrial tRNA m^1^G37 (11). In plants, the enzymes catalysing m^1^G37 methylation have not been identified to date.

N^1^-methylguanosine has been described in eukaryotic tRNAs at two positions; at position 37 catalysed by Trm5, and the other at position 9 catalysed by Trm10 (26). Trm5 in humans, yeast, and *Pyrococcus abyssi* has been described as having multifunctionality (23, 24, 27). In contrast to TrmD which requires a guanosine at position 37, human Trm5 can also recognise and methylate inosine at position 37 with some limited activity (24). Similarly,Trm5p have also been shown to catalyse inosine to m^1^I modification in yeast in a two-step reaction, where the first adenosine-to-inosine modification was mediated by Tad1p (10, 18, 22, 28). As m^1^G is an intermediate during the modification of guanosine to wybotusine (yW), tRNAs from *trm5* mutants were also devoid of yW (22). The yeast Trm5p protein has been shown to be localised to the cytoplasm and mitochondria (23) and it is thought that Trm5p protein present in the mitochondria is required to prevent unmodified tRNA affecting translational frameshifting (22). In the unicellular parasite *Trypanosome brucei*, Trm5 was located in both the nucleus and mitochondria and reducing Trm5 expression led to reduced mitochondria biogenesis and impaired growth (29). Interestingly, Trm5 and m^1^G37 were shown to be essential for mitochondrial protein synthesis but not cytosolic translation (29).

Little is known about tRNA modifying enzymes in plants. Here, we report the identification and functional analysis of *AtTrm5* (At3g56120) from the model plant *Arabidopsis thaliana*. We demonstrate that *Attrm5* mutants are slower growing, have reduced shoot and root biomass and display late flowering. Furthermore, we demonstrate that *in vitro TRM5* is required for m^1^G37 and m^1^I37 methylation at the position 3’ to the anticodon and *in vivo* tRNAs enriched from *Attrm5* plants have reduced m^1^G and m^1^I.

## Methods

### Plant material and root growth experiments

*Arabidopsis thaliana* (Columbia accession) wild type and mutant plants were grown in Phoenix Biosystems growth under metal halide lights as previously described (30). For plate experiments, seeds were first surface sterilized, plated on ½ MS medium supplemented with 1% sucrose and sealed as previously described (30, 31). All plants were grown under either long-day photoperiod conditions of 16 h light and 8 h darkness or short-day photoperiods of 10 h light and 14 h darkness.

Characterization of the mutant alleles, *trm5-1* (SALK_022617) and *trm5-2* (SALK_032376) are as described previously (32). The *tad1-2* mutant was used as described previously (10). Nucleotide sequence data for the following genes are available from The *Arabidopsis* Information Resource (TAIR) database under the following accession numbers: TRM5 (At3g56120), At4g27340, At4g04670, and TAD1 (At1g01760).

Analysis of root phenotypes was carried out on 11-day-old seedlings grown on ½ MS agar plates. A flatbed scanner (Epson) was used to non-destructively acquire images of seedling roots grown on the agar surface. Once captured, the images were analyzed by software package RootNav (33, 34).

### Plasmid construction and generation of transgenic plants

For the 35SCaMV:TRM5 construct, the full-length genomic region of At3g56120 including the 5’UTR and 3’UTR was amplified from Col-0 genomic DNA template with primers provided in (Supplementary Table S1) and cloned into Gateway entry vector pENTRTM/SD/D-TOPO (Invitrogen). The insert was sequenced and then cloned into the binary destination vector pGWB5 by an LR recombination reaction, using the Gateway cloning system following the manufacturers protocol (Invitrogen), resulting in the 35S:TRM5 construct. For the TRM5_Pro_::TRM5 construct, the full-length genomic region of At3g56120 that included the promoter, 5’UTR and 3’UTR was amplified from Col-0 genomic DNA template with primers provided in (Supplementary Table S1) and cloned into Gateway entry vector PCR8 TOPO-TA (Invitrogen). The insert was sequenced and then cloned into the a modified (35S promoter removed) destination pMDC32 vector, using the Gateway cloning system (35) following the manufacturers protocol (Invitrogen), resulting in TRM5_Pro_::TRM5. The 35S:TRM5 construct was transformed into *A. thaliana* wild type Col-0 plants or *trm5-1* mutant plants by *Agrobacterium-mediated* floral dip method respectively (36). The TRM5_Pro_::TRM5 construct was transformed into *trm5-2* mutant plants by *Agrobacterium-mediated* floral dip method. Transgenic plants were selected on ½ MS media supplemented with 50 μg ml^−1^ kanamycin. TRM5 transcript abundance was assessed in at least five independent T_1_ plants using qRT-PCR and two lines showing the highest TRM5 transcript levels were carried through to homozygous T_3_ generation for phenotypic analysis.

### Sub-cellular localization of TRM5

For analysis of subcellular localization of TRM5, the construct described above was introduced into *A. tumefaciens* strain GV3101 and transiently expressed in 5-week-old *Nicotiana benthamiana* leaves. Fluorescence was analyzed using a confocal laser-scanning microscope (Zeiss microscope, LSM700) and excited with 488-nm line of an argon ion laser. GFP fluorescence was detected via a 505- to 530-nm band-pass filter. The cut leaves were immersed in 10 μM 4′, 6-diamidino-2-phenylindole (DAPI) at room temperature for 45 min, and then washed with PBS for 3 times (5 min each). The blue fluorescence of DAPI was imaged using 404-nm line for excitation and a 435- to 485nm band pass filter for emission.

### Shoot apical meristem sections

14, 18, or 22-day-old seedlings were fixed for 1 day in FAA containing 50% ethanol, 5% acetic acid and 3.7% formaldehyde. The samples were then dehydrated through an ethanol series of five one-hour steps (50%, 60%, 70%, 85%, 95% ethanol) ending in absolute ethanol (100%). The ethanol was gradually replaced with Histoclear containing safranine to stain the tissue, as following: incubated in a Histoclear series (75:25, 50:50, 25:75 Ethanol: Histoclear) for 30 min and followed by twice one-hour incubation in 100% Histoclear. The paraffin was polymerised by baking overnight at 60 °C, and the samples were embedded in paraffin. Sections were cut, attached to slides and dried on a slide warmer overnight at 42 °C to allow complete fixation. The shoot apical meristem was observed using light microscopy.

### Quantitative RT-PCR

For the transcription profiling of flowering-related genes and circadian clock-related genes, 17-day-old seedlings were sampled from Zeitgeber time (ZT) 1 and collected every 3h during the day and night cycles, respectively. Total RNA was extracted the leaf samples using Trizol reagent (Invitrogen). The relative expression levels of *AtTRM5* were determined using quantitative real-time PCR with gene-specific primers (Supplementary Table S1). Real-time PCR was performed using the StepOnePlus real-time PCR system (Applied Biosystems) using Absolute SYBR Green ROX mix (Applied Biosystems) for quantitation. Three biological replicates were carried out for each sample set. The relative expression was corrected using a reference gene *EF1alpha* (At5g60390) and calculated using the 2^−ΔΔCt^ method as described previously (10).

### mRNA-sequencing

Total RNA, 1 ug, was extracted from 20-day-old *Arabidopsis* leaf samples using Trizol reagent (Invitrogen) and purified using the RNAeasy Mini RNA kit (Qiagen). One hundred nanograms of RNA were used for RNA-seq library construction according the manufacturer’s recommendations (Illumina). First-strand cDNA was synthesized using SuperScript II Reverse Transcriptase (Invitrogen). After second strand cDNA synthesis and adaptor ligation, cDNA fragments were enriched, purified and then sequenced on the Illumina Hiseq X Ten. Three biological replicates were used for RNA-seq experiments.

### tRNA purification and tRNA-sequencing

Total RNA was isolated from wild type and *trm5* 10-day-old *Arabidopsis* seedlings using the Spectrum Plant total RNA kit (SIGMA-ALDRICH) and contaminating DNA removed using DNase I (SIGMA-ALDRICH). To enrich for tRNAs, 10μg of total RNA was separated on a 10% polyacrylamide gel, the region containing 65-85 nts was removed and RNA was purified as previously described (31). Purified tRNAs were used for library construction using NEB Ultradirectional RNA library kit. Given the short sequences of tRNAs, the fragmentation step of the library preparation was omitted, and samples were quickly processed for first-strand cDNA synthesis after the addition of the fragmentation buffer. The remaining steps of library construction were performed as per the manufacturer’s instructions. Illumina sequencing was performed on a MiSeq platform at The Australian Cancer Research Foundation (ACRF) Cancer Genomics Facility, Adelaide.

### Yeast complementation

AtTrm5 (At3g56120) and ScTrm5 (YHR070W) was PCR amplified from cDNA and cloned into pYE19 using a Gibson assembly reaction (NEB). Mutant AtTrm5 (R166D) was generated by synthesising gene blocks (IDT) with nucleotides that mutated the translated proteins at R166 and the gene block was cloned into pYE19 using a Gibson assembly reaction (NEB). Yeast Δtrm5 (*Mat a, hisD1, leu2D1, met15D0, trm5:KanMX*) was previously described (37). Recombinant plasmids were transformed into Δ*trm5* mutant strain and the resulting strain was analysed for growth phenotypes and m^1^G nucleoside levels.

### AtTrm5, AtTAD1, ScTrm5 protein expression and purification and tRNA methylation

Full length AtTrm5, mutant AtTrm5 (R166D) and AtTAD1 (At1g01760) cDNAs were cloned into pGEX resulting in GST-AtTrm5, GST-AtTRM5-mutant, GST-TAD1, respectively. Mutant AtTAD1 (E76S) was generated by synthesising a gene block (IDT) with mutated nucleotides and the gene block was cloned into pGEX using a Gibson assembly reaction (NEB) resulting in GST-TAD1 mutant. IPTG (0.5 μM) was used to induce expression of the proteins and the recombinant proteins were purified on a GST resin column (ThermoFisher Scientific). tRNA-Asp-GUC or tRNA-Ala-AGC were transcribed in vitro with T7 polymerase (Promega). Methylation reactions were performed in 100 mM Tris-HCl, 5mM MgCl_2_, 100 mM KCl, 2mM DTT, 50mM EDTA, 0.03 mg/mL BSA and 25 μM AdoMet. Substrate tRNA was provided in a final concentration of 1-5 μM, AtTrm5 or AtTrm5 mutant proteins from 6.0 to 12 μM and AtTAD1 or AtTAD1 mutant proteins at 5.0 μM.

### Bioinformatics analysis of mRNA-seq and tRNA-seq

#### Global Mapping

Reads were first adapter trimmed using Trim galore v0.4.2 (http://www.bioinformatics.babraham.ac.uk/projects/trimgalore/) with default parameter. The quality of the trimmed reads were checked using Fastqc (https://www.bioinformatics.babraham.ac.uk/projects/fastqc/) and ngsReports (38). Reads were then globally mapped to *Arabidopsis* reference genomes (TAIR10) (39) using STAR v2.5.3 (40). Mapped reads were counted using featureCounts (41). The raw read counts were normalized using sample size factor for sequence depth and differential expression analysis between wild type and *trm5-1* mutant samples was performed using DESeq2 v1.18.1 (42). The tRNA-enriched reads were mapped to tDNA reference acquired from GtRNAdb (generated from tRNA-Scan SE v2.0) of *Arabidopsis thaliana* derived from TAIR10 reference genome (39, 43–45) using segemehl v0.2.0 (45). The mapped reads were then processed for variant calling using GATK v3.7 (46). Haplotype Caller in GATK v3.7 was implied to call for variants, with the variant filtered using hard filtering as recommended.

#### Proportion estimation

The common SNPs identified in wild type and *trm5-1* samples were cross compared. SNPs at position 37 were extracted and analysed using vcftools (47). Changes in base pair modifications were indicated by base substitution due to the property of next generation sequencing as mentioned in (11, 48). The ratio of the expected A-to-T conversion in wild type samples and both the ratio of A-to-T (indicating no change in comparison to wild type) and A-to-G in *trm5-1* were analyzed as an indication of m^1^I depletion. Identification of other base changes was attempted to identify putative m^1^I modification.

### Sequence analysis of tRNA editing

tRNA purification and tRNA editing analysis were performed as described previously (10). Cytosolic tRNA-Ala (AGC) were amplified by reverse transcriptase (RT)-PCR with specific primers (tRNA-Ala-f and tRNA-Ala-r, Supplementary Table S1), and purified PCR products were directly Sanger sequenced.

### TMT-based proteome determination and data analysis

Total protein was extracted from 20-day-old *Arabidopsis* leaf samples and purified according to a method described by (49). Protein digestion was performed according to FASP procedure described previously (50), and 100 ųg peptide mixture of each sample was labelled using TMT reagent according to the manufacturer’s instructions (Thermo Fisher Scientific). LC-MS/MS analysis was performed on a Q Exactive mass spectrometer (Thermo Fisher Scientific) that was coupled to Easy nLC (Thermo Fisher Scientific). MS data was acquired using a data-dependent top20 method dynamically choosing the most abundant precursor ions from the survey scan (200-1800 m/z) for HCD fragmentation (Shanghai Applied Protein Technology Co., Ltd). Determination of the target value is based on predictive Automatic Gain Control (pAGC). SEQUEST HT search engine configured with Proteome Discoverer 1.4 workflow (Thermo Fischer Scientific) was used for mass spectrometer data analyses. The latest *Arabidopsis* protein databases (2018) was downloaded from http://www.arabidopsis.org/download/index-auto.jsp?dir=/downloadfiles/Proteins and configured with SEQUEST HT for searching the datasets. The following screening criteria was deemed as differentially expression: fold-changes >1.2 or <0.83 (up- or down-regulation) and *P* < 0.05 were considered significant.

### tRNA nucleoside analysis

tRNAs were purified as previously described (31). Twenty-five μg of tRNAs were digested with P1 nuclease (Sigma-Aldrich) and 1.75 units of calf intestine alkaline phosphatase in 20 mM Hepes–KOH (pH 7.0) at 37 °C for 3 h. API 4000 Q-Trap mass spectrometer coupled with LC-20A HPLC system was used for nucleoside separation and detection, with a diode array UV detector (190–400 nm) set for positive ion mode.

## RESULTS

### Identification of At3g56120 as a TRM5 homolog

In yeast (*Saccharomyces cerevisiae*), m^1^G37 nucleoside modification is catalysed by ScTrm5 (22). We searched for *Arabidopsis thaliana* homologs by using blastp and HMMER and identified a high confidence candidate, At3g56120, with 49% similarity to ScTrm5 (Supplementary Figure S1). Alignment of At3g56120 with yeast, human, Drosophilia, Pyrocococcus, and Methanococcus Trm5 homologs identified three conserved motifs and catalytically required amino acids (R166, D192, E206) present in At3g56120 (Supplementary Figure S1). We subsequently will refer to At3g56120 as AtTrm5. In the *Arabidopsis* genome, AtTrm5 has homology to At4g27340 and At4g04670 and both genes were recently been named as Trm5B and Trm5C respectively (Supplementary Figure S1 and (51)). We have also identified Trm5 homologs in algae, bryophytes and vascular plants (Figure 1A).

**Figure 1.**
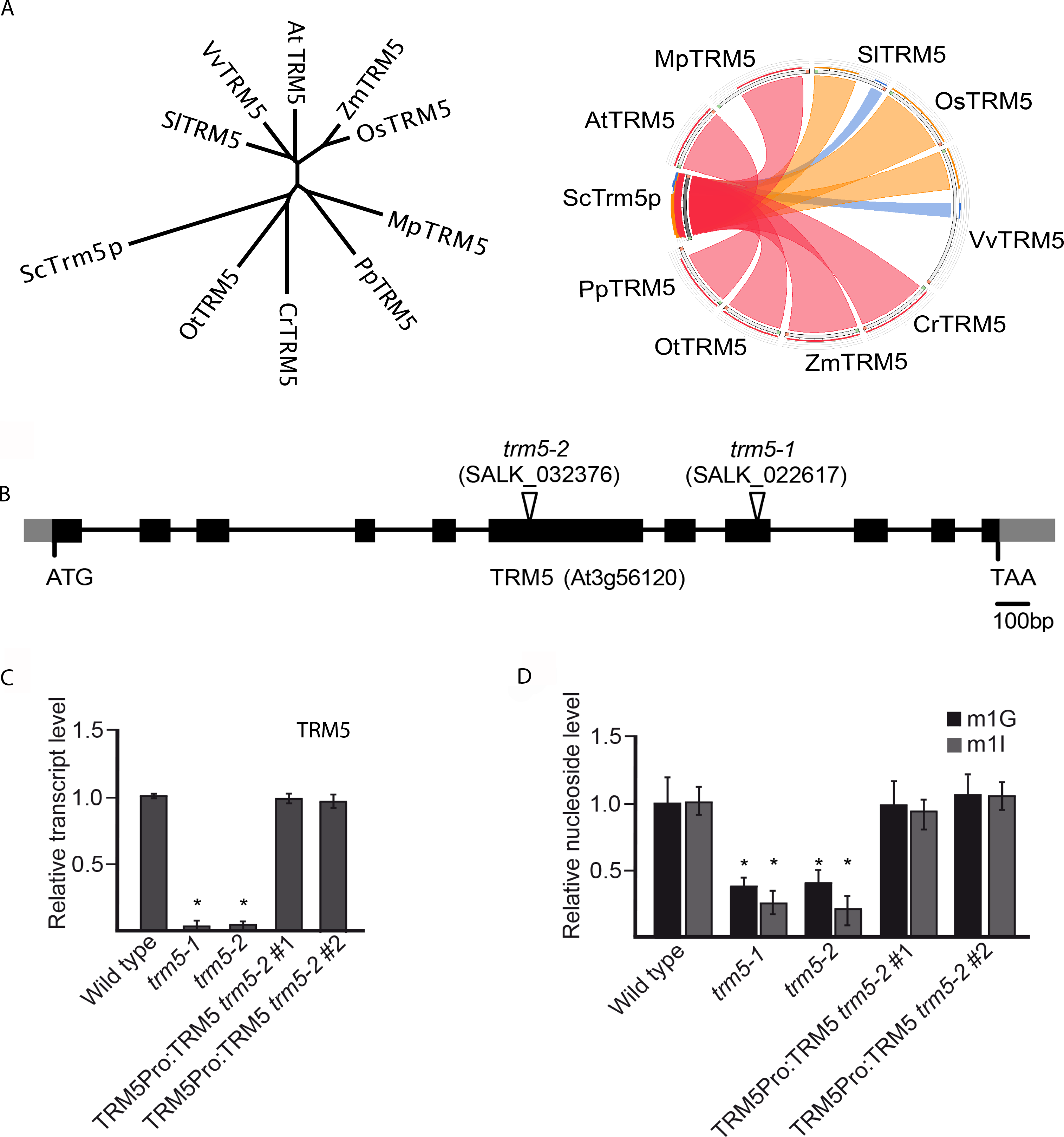
TRM5 is conserved in plants and has dual-functionality in modifying RNA bases. (**A**) Unrooted phylogenetic tree and sequence conservation Circos plot of putative TRM5 proteins from yeast (Sc), tomato (Sl), grape (Vv), *Arabidopsis* (At), maize (Zm), rice (Os), *Marchantia* (Mp), *Physcomitrella* (Pp), *Chlamydomonas* (Cr), and *Ostreococcus* (Ot). (**B**) Exon-intron structure of the putative TRM5 locus (At3g56120) showing the T-DNA insertion sites of the *trm5-1* and *trm5-2* alleles (as indicated by the open triangles). Black boxes and grey boxes represent coding regions and untranslated regions, respectively. (**C**) Relative transcript level detected by qRT-PCR in wild type, *trm5-1* or *trm5-2* seedlings. (**D**) Relative nucleoside level of modification m^1^G and m^1^I detected by immunoprecipitation in wild type, *trm5-1* or *trm5-2* seedlings.

To functionally characterize AtTrm5 we isolated two T-DNA insertions, SALK_022617 and SALK_032376, and identified homozygous mutant plants for each insertion (Figure 1B). SALK_022617 and SALK_032376 were named *trm5-1* and *trm5-2*, respectively. Next, we measured AtTrm5 mRNA abundance in both mutants and detected almost no transcripts in both mutants (Figure 1C). We generated a genomic construct of AtTrm5 that contained the endogenous promoter, coding region and UTRs, transformed the construct into *trm5-2* and demonstrated that the AtTrm5 mRNA levels were similar in two complemented lines when compared to wild type plants. Subsequently, the extracted tRNAs from wild type and the *trm5* mutants were purified, digested and modified nucleosides measured by mass spectrometry (Figure 1D). In both *trm5* mutant alleles, nucleoside m^1^G levels were reduced to about 30% of the wild type and m^1^G levels were restored to wild type levels in both complemented lines (Figure 1D). Nucleoside m^1^G is present at tRNA positions 9 and 37 (22), therefore the residual m^1^G levels in both *trm5* mutants may be the result of tRNA m^1^G at position 9.

In yeast, Trm5 has also been reported to also catalyse m^1^I on tRNAs (10, 22, 51). We therefore measured m^1^I levels in purified tRNAs from both *Arabidopsis trm5* mutants and wild type control plants. In both *trm5-1* and *trm5-2* mutant alleles, nucleoside m^1^I levels were reduced to about 10% of wild type levels and were restored to wild type levels in plants of both complemented lines (Figure 1D). In summary, in *Arabidopsis thaliana* we identified At3g56120 as a Trm5 homolog in *Arabidopsis thaliana*, identified two AtTrm5 mutant alleles, *trm5-1* and *trm5-2*, and both mutants showed a significant reduction in m^1^G and m^1^I.

### AtTRM5 m^1^G methyltransferase activity

To test AtTrm5 m^1^G methyltransferase activity *in vivo*, a yeast, a *Δtrm5* mutant strain in yeast (*Saccharomyces cerevisiae*, Sc) that is defective for the tRNA m^1^G37 modification was used for genetic complementation. The mutant not only has defective tRNA m^1^G37 but also a slow growth phenotype when compared to wild type or a congenic strain (Figure 2A). Full-length *ScTrm5* and *AtTrm5* were cloned into yeast expression vectors. From the *AtTrm5* expression vector, a catalytically inactive mutant *Attrm5* R166D was generated by site-directed mutagenesis. After the three vectors had been transformed into the yeast *Δtrm5* mutant, cell growth and m^1^G nucleoside levels were observed (Figure 2). Not only were the slow growth and nucleoside levels rescued when expressing *ScTrm5* but they were also rescued when expressing *AtTrm5* (Figure 2A and B). However, the catalytically inactive *Attrm5* R166D did not rescue either the slow growth or m^1^G nucleoside levels (Figure 2A and B).

**Figure 2.**
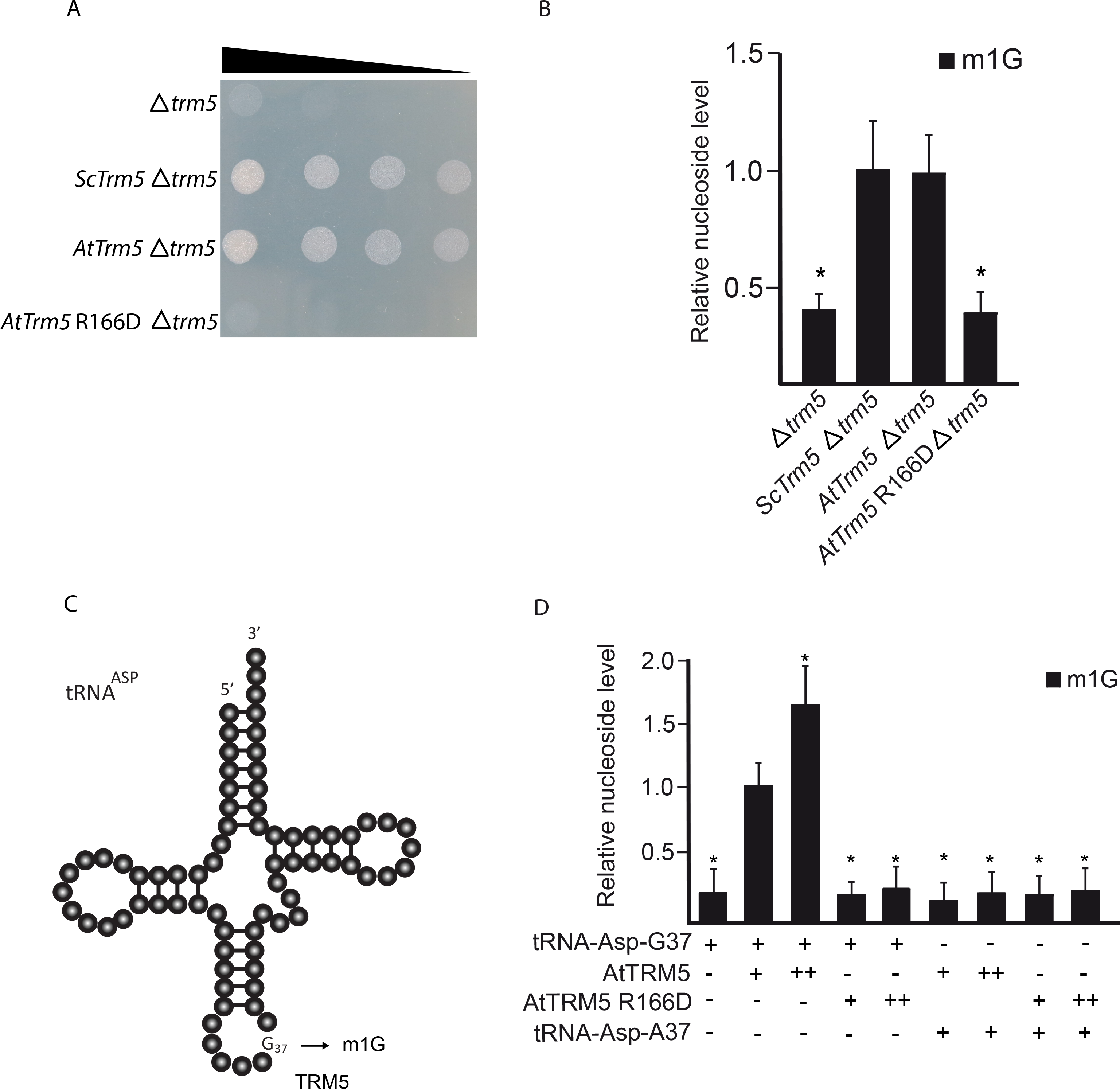
TRM5 modifies tRNA_37_ guanosine (G) to m^1^G in yeast. (**A**) Serial-dilution growth assay. Complementation experiment of TRM5 from yeast (Sc), *Arabidopsis thaliana* (At) or a catalytically inactive mutant AtTrm5 R166D in the yeast *trm5* mutant. (**B**) Relative nucleoside level of modification m^1^G quantified by immunoprecipitation of each complemented yeast strains with either ScTRM5 or AtTRM5. (**C**) Proposed model of TRM5-mediated m^1^G modification of yeast tRNA-Asp. (**D**) Relative nucleoside level of modification m^1^G with varying conditions of tRNA-Asp-G37, AtTRM5, AtTRM5 catalytic mutant, and tRNA-Asp-A37. + indicates presence, ++ indicates two-fold increase, - indicates absence.

To test the m^1^G methyltransferase activity of AtTRM5 *in vitro*, we incubated purified recombinant proteins with tRNA substrates and measured the m^1^G levels. We expressed *AtTRM5* as a GST fusion protein and purified the recombinant GST-AtTRM5 protein. We also generated a catalytically inactive GST-AtTRM5 recombinant protein by using site directed mutagenesis, expressed and purified the GST-AtTRM5 recombinant fusion protein. In yeast, ScTrm5 methylates tRNA-His-GUG, tRNA-Leu-UAA, tRNA-Asp-GUC and other tRNA isoacceptors (10, 22, 51). Yeast tRNA-Asp-GUC RNA transcripts were generated *in vitro* by using T7 polymerase, the tRNA transcripts incubated with the recombinant fusion proteins in the presence of the methyl donor S-adenosyl-methionine (AdoMet) and m^1^G nucleoside levels were measured (Figure 2D). m^1^G was detected only when AtTRM5 was provided (Figure 2D) and in a dosage dependent manner. No m^1^G was detected when the catalytically inactive mutant AtTRM5 was provided (Figure 2D). To test the specificity of the methyltransferase activity on tRNA-Asp guanine at position 37, the guanine nucleotide was mutated to an adenine nucleotide (tRNA-Asp-A37) and the m^1^G methyltransferase activity was measured. No m^1^G was detected after incubation with the fusion proteins (Figure 2D). The overall results of the yeast complementation experiments suggest that guanosine methylation occurred at position 37 of tRNA.

### AtTRM5 tRNA m^1^I methyltransferase activity

Previously in plants, TAD1 was demonstrated to oxidatively deaminate adenosine at position 37 of tRNA-Ala-(UGC) to inosine, and subsequently methylated by an unknown enzyme to N1-methylinosine (m1I; Figure 3A). Human TRM5 has been reported to methylate tRNA I37 but with limited activity (24). Given our observation that *Attrm5* mutant plants had reduced m^1^I (Figure 1D), we asked the question whether AtTRM5 has methyltransferase activity on tRNA I37. We developed a two-step approach, whereby purified AtTAD1 was first incubated with the substrate tRNA-Ala-A37 to produce tRNA-Ala-I37 and then the inosine methyltransferase activity of AtTRM5 was measured by incubating AtTRM5 with the tRNA-Ala-I37 substrate. Previously, in yeast ScTAD1 was demonstrated to deaminate tRNA-Ala-A37 to tRNA-Ala-I37 *in vitro* (22) and *Arabidopsis thaliana tad1* mutants were reported to have reduced tRNA-Ala-I37 (10). We expressed *AtTAD1* as a GST fusion protein and purified the recombinant GST-AtTAD1 protein. We also generated a catalytically inactive GST-AtTAD1 mutant recombinant protein by using site-directed mutagenesis, and expressed and purified the GST-AtTAD1 mutant fusion protein. In a two-step assay, yeast tRNA-Ala-UGC RNA transcripts were generated by *in vitro* transcription using T7 RNA polymerase, the tRNA transcripts were then incubated with the recombinant fusion proteins in the presence of Mg^2+^ and methyl donor S-adenosyl-methionine (AdoMet), and the m^1^I nucleoside levels measured (Figure 3B). m^1^I was detected only when GST-AtTAD1 and GST-AtTRM5 were provided, and its production occurred in a dosage-dependent manner (Figure 3B). No m^1^I was detected when the catalytically inactive mutants AtTAD1 E76S or AtTRM5 R166D were provided (Figure 3B). To test the specificity of the methyltransferase activity on tRNA-Ala alanine at position 37, the alanine nucleotide was mutated to a cytosine nucleotide (tRNA-Ala-C37) and the m^1^I methyltransferase activity was measured. No m^1^I was detected after incubation with the fusion proteins (Figure 3B). Collectively, these findings interestingly suggest that inosine methylation also occurrs at position 37, in addition to guanosine methylation.

**Figure 3.**
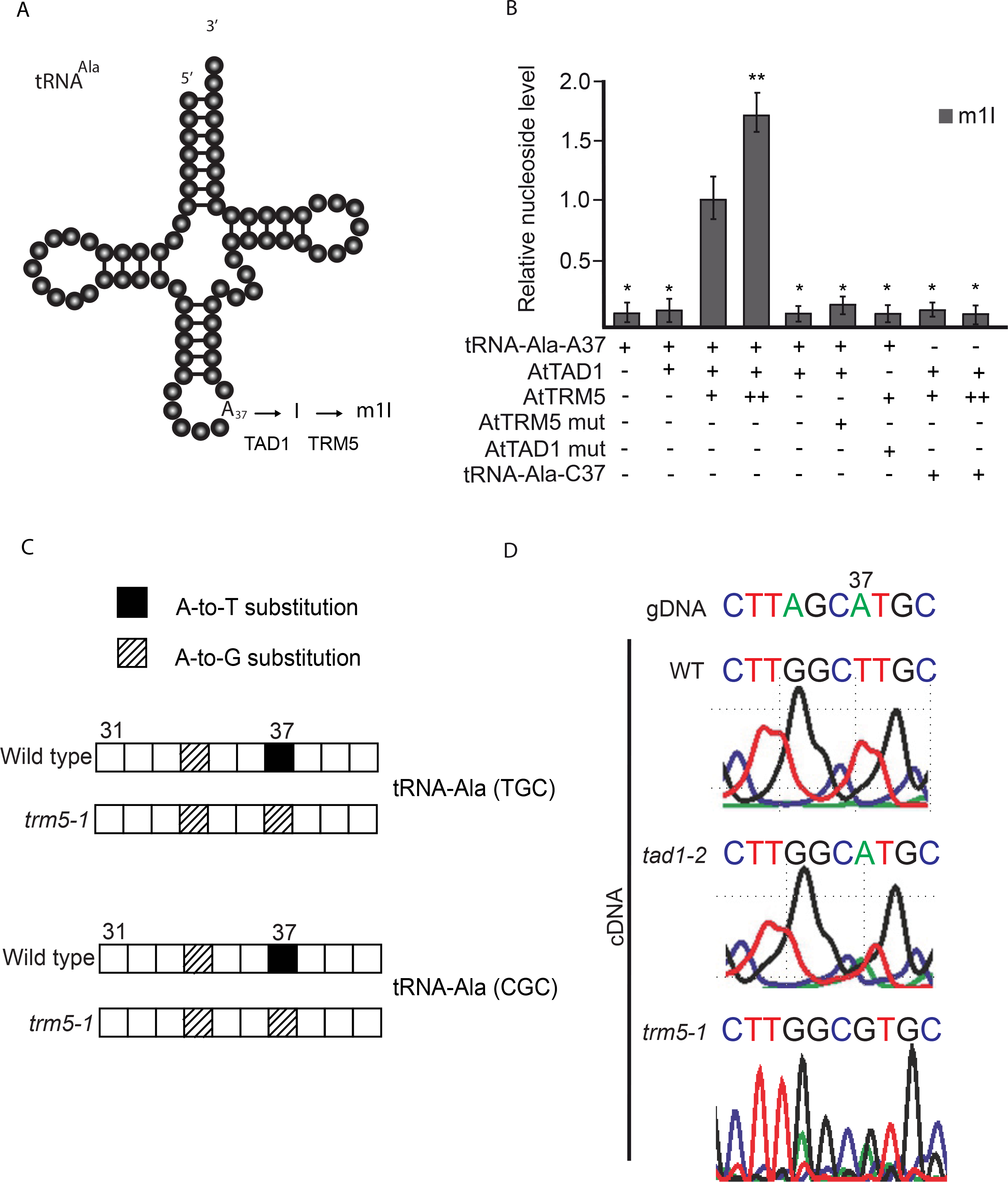
TRM5-mediated m^1^I modifications in two-step reaction. (**A**) Proposed two-step modification model of TRM5-mediated m^1^G modification of tRNA-Ala. (**B**) Relative nucleoside level of modification m11 with varying conditions of tRNA-Asp-A37, AtTAD1, AtTRM5 mutant, AtTAD1 mutant, and tRNA-Ala-C37. + indicates presence, ++ indicates two-fold increase, - indicates absence. (**C**) tRNAs were enriched, deep sequenced, aligned using segemehl to tRNA references and the modifications present were inferred from observed base substitutions between wild type and *trm5-1*. Position 37 in the gDNA is an adenine. In wild-type cDNA, a thymine was detected and in *trm5-1* cDNA, a guanine was observed. Results for tRNA-Ala^(TGC)^ and tRNA-Ala^(CGC)^ are shown. The average modification rate from 3 biological replicates is shown. (**D**) tRNA-Ala^(AGC)^ was PCR amplified from wild type, *tad1-2* and *trm5-1* and Sanger sequenced. Position 37 in the gDNA is an adenine. In wild-type cDNA, a thymine was detected, in *tad1-2* cDNA an adenine was observed and in *trm5-1* a guanine was detected.

To test the AtTRM5 inosine methyltransferase activity *in vivo*, we measured tRNA position modifications by cDNA sequencing from either mutant or wild type plants (Figure 3C and D). In the sequencing assay, modification events at position 37 of tRNAs can be directly detected by sequencing of amplified cDNA obtained by reverse transcription and comparison to the DNA reference sequence as inosine is read as 1 guanine (G) and m^1^I is read as thymine (T) by the reverse transcriptase (11, 48). As expected, we detected substitutions of A in the reference to T at position 37 for tRNA-Ala-TGC and tRNA-Ala-CGC by using Illumina sequencing in wild-type plants which 11 is consistent with the presence of m^1^I (Figure 3C). m^1^I modifications have previously been described in tRNA-Ala at position 37 in eukaryotes (10, 28, 52). In the *trm5-1* mutant, no T’s were detected at position 37 (Figure 3C) which is consistent with AtTRM5 acting as a tRNA m^1^I methyltransferase at position 37. Based on our *in vitro* assay results, AtTAD1 first deaminates tRNA-Ala-A37 to tRNA-Ala-I37, and then AtTRM5 methylates tRNA-Ala-I37 to tRNA-Ala-m^1^I37. We confirmed this pathway in *tad1* and *trm5* mutant plants by Sanger sequencing of tRNA-Ala-AGC (Figure 3D). As expected, at position 37, A was substituted to T in the wild type, whereas in *trm5* mutants a G and in *tad1* mutants an A were observed (Figure 3D). These sequencing results are consistent with AtTAD1 first deaminating A37 to I37 as previously reported by (10), and AtTRM5 then methylating I37 to m^1^I. We also attempted to detect the putative loss of m^1^G in the tRNA-sequencing data, as it has been reported that m^1^G is prone to be called as a T in sequencing (48). However, this was not observed in our datasets. Together, our *in vitro* and *in vivo* data provide support for AtTRM5 possessing tRNA m^1^I methyltransferase activity.

### AtTRM5 is localized to the nucleus

In yeast, ScTRM5 is localized to both the nucleus and mitochondria (23, 53). Localisation to mitochondria is thought to be important as yeast strains with only nuclear-localized ScTRM5 exhibited a significantly lower rate of oxygen consumption (23). In order to determine to which subcellular compartment(s) AtTRM5 is localized in *Nicotiana benthamiana*, we fused TRM5 to the Green Fluorescent Protein (GFP) reporter, transiently infiltrated the construct into leaves and performed laser-scanning confocal microscopy to detect GFP fluorescence. To unambiguously identify the nucleus, we stained the cells with DAPI. When we imaged the cells (n=100), we observed distinct DAPI fluorescence in a single large circular structure per cell, as expected for the nucleus (Figure 4). Next, we imaged the same cells for GFP fluorescence (Figure 4C) and overlayed the DAPI and GFP fluorescence. The two fluorescence signals showed perfect overlap (Figure 4D). We then searched for nuclear localisation signals (NLS) using the LOCALIZER (http://localizer.csiro.au/), LocSigDB (http://genome.unmc.edu/LocSigDB/), and cNLS mapper (http://nls-mapper.iab.keio.ac.jp/cgi-bin/NLS_Mapper_form.cgi) programs (54–56). While LOCALIZER and LocSigDB did not predict any canonical or bipartite NLS, cNLS mapper predicted with high confidence a 29 amino acid importin α-dependent NLS, QKGCFVYANDLNPDSVRYLKINAKFNKVD, that starts at amino acid 236. Previous finding in yeast suggested that no common canonical or bipartite NLS was detected in ScTrm5 (22), which explains the outcome from LOCALIZER and LocSigDB. In summary, we conclude that, unlike in yeast, AtTRM5 in *Arabidopsis* is only localized to the nucleus and may be imported from the cytoplasm into the nucleus by the importin α-dependent pathway.

**Figure 4.**
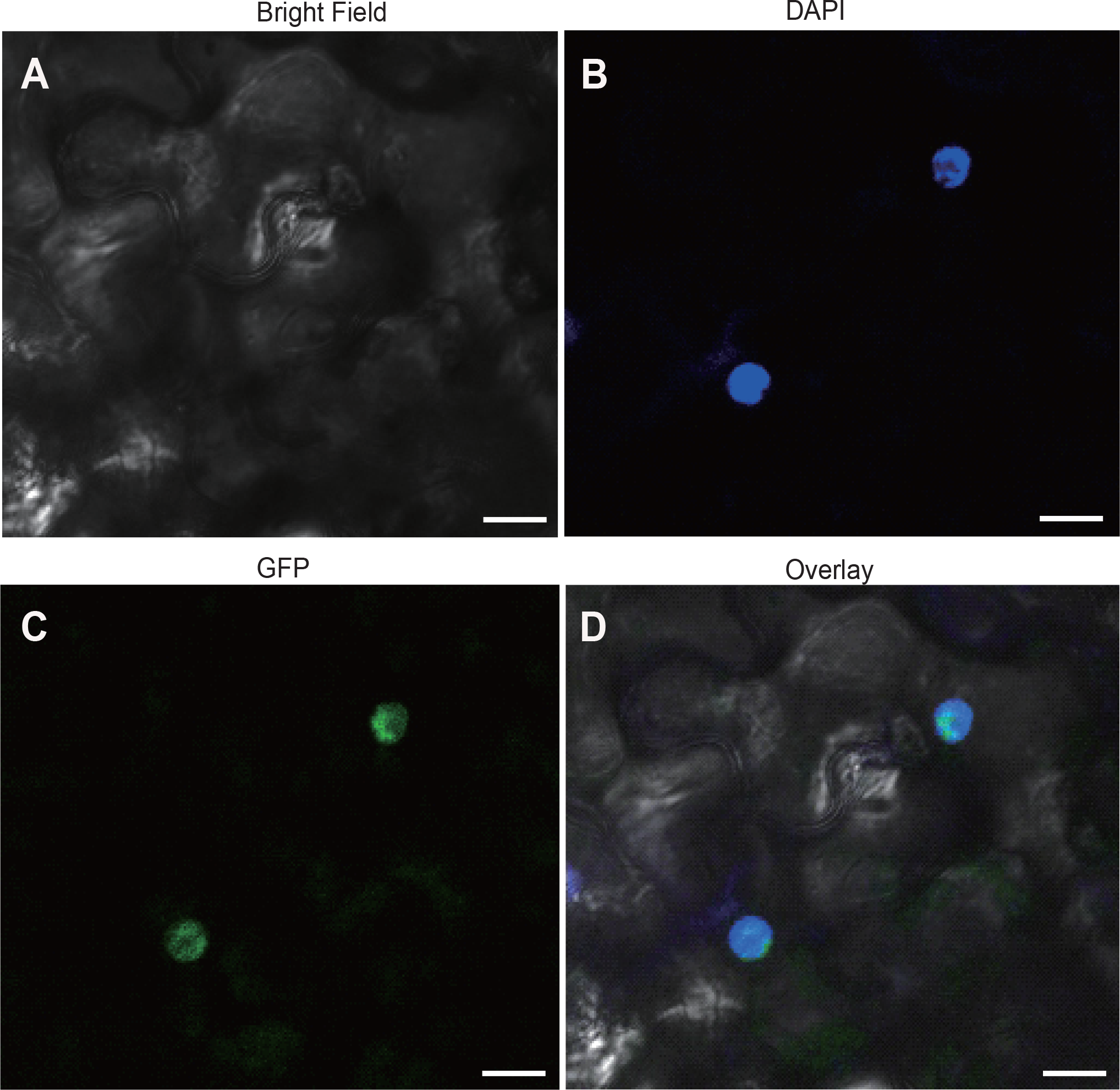
Subcellular localisation of TRM5 translational reporter protein in *Nicotiana benthamiana* leaves. TRM5 was fused to Green Fluorescent Protein (GFP) to yield a TRM5-GFP translational fusion recombinant protein. The construct was transiently expressed in *N. benthamiana* leaves and subcellular localisation was determined by confocal laser-scanning microscopy. (**A**) Bright field, (**B**) DAPI stained, (**C**) GFP fluorescence, (**D**) overlay of DAPI and GFP imagines. Scale bars = 20 μm.

### Proteins involved in photosynthesis are affected in *trm5* mutant plants

Next, we performed a proteomic analysis to identify proteins that differentially accumulate in *trm5-1* plants when compared to the wild type by using the Tandem Mass Tag (TMT) method (Figure 5). A total of 61571 peptide-spectrum match (PSM) were recorded, corresponding to 29011 peptides and 23055 unique peptides, respectively. 5242 protein groups were identified by blastp searches against the TAIR_pep database. Proteins with fold changes ≥1.20 or ≤0.83 and a significance level of *P* ≤0.05 were considered to be differentially expressed. In this way, a total of 263 proteins were identified (Supplementary Table S2). 102 proteins were upregulated, and 151 proteins were downregulated in *trm5* (Figure 5A). GO annotation of these differentially accumulating proteins revealed enrichment of the GO terms thylakoid, chloroplast, and photosystem I (Figure 5B). KEGG annotation revealed enrichment of proteins involved in photosynthesis, and photosynthetic proteins (Figure 5C). Taken together, the GO and KEGG analysis demonstrated that most differentially accumulating proteins are involved in processes related to photosynthesis.

**Figure 5.**
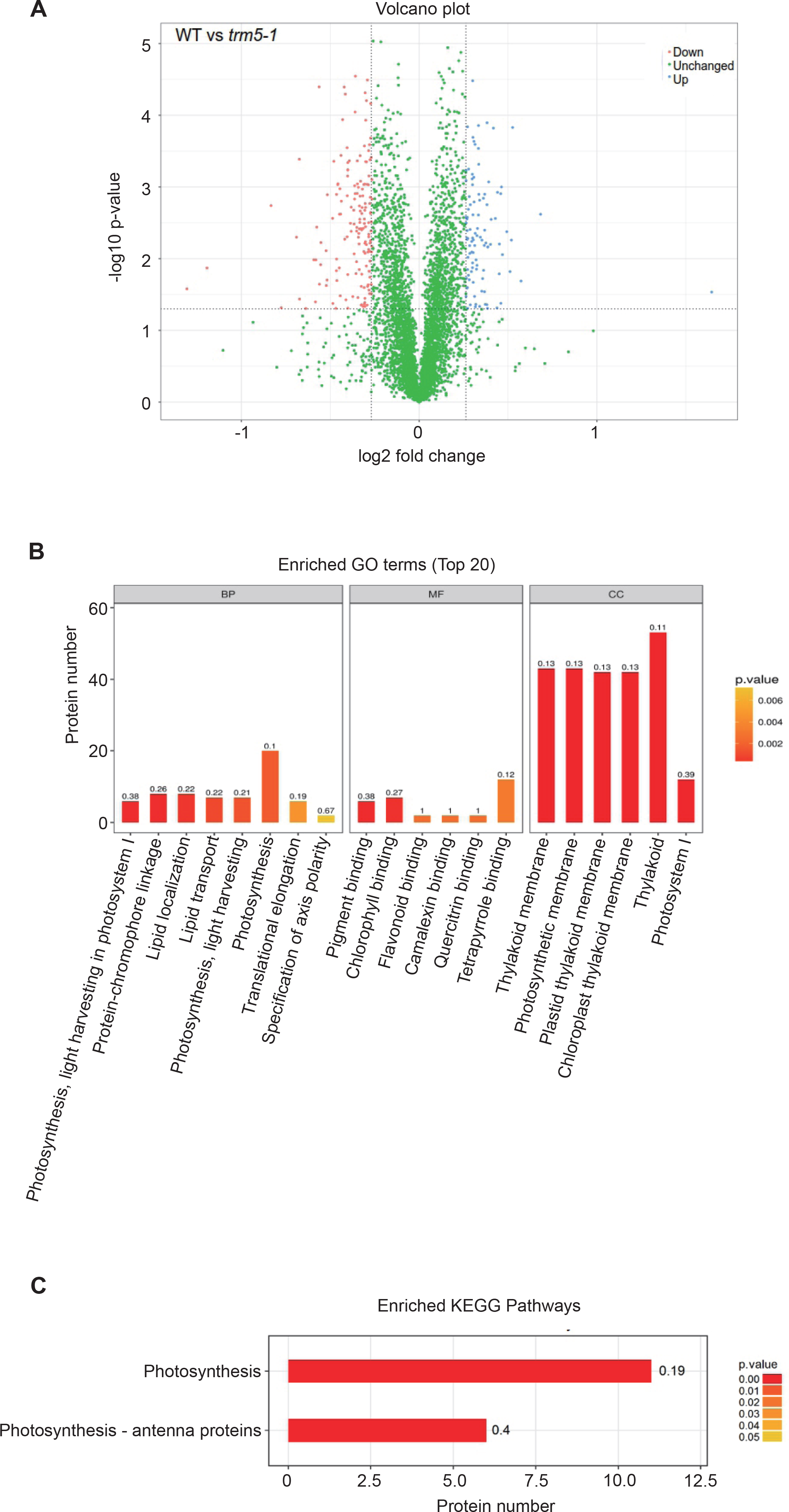
Proteomic analysis of wild type and *trm5-1*. (**A**) Volcano plot of differentially abundant proteins between wild type and *trm5-1*. In *trm5-1*, proteins that were increased in abundance are represented as blue dots, proteins decreased in abundance as red dots, with threshold fold change > 1.2 or < 0.83 (increased or decreased) and P-value < 0.05. There were 102 proteins increased and 161 proteins decreased in *trm5-1*. (**B**) GO and (**C**) KEGG terms enrichment analysis. Each differentially abundant protein was first annotated in the GO or KEGG databases, enrichment analysis was performed based on annotated differentially expressed proteins in wild type and *trm5-1*. The top 20 enriched GO terms from Biological Processes (BP), Molecular function (MF), and Cellular Component (CC) are reported.

Defects in tRNA m^1^G methylation can be expected to affect mRNA translation, particularly of proteins that have high numbers of affected codons. Therefore, we were interested in identifying genes that showed reduced expression at the protein level in our proteomics analysis of *trm5* plants, but no detectable reduction in mRNA abundance. To identify such mRNAs, we performed RNA-seq on wild type and *trm5* plants. We identified 1186 transcripts that were reduced in abundance in *trm5* and 580 transcripts that were increased in abundance by at least 2 fold and hierarchically clustered these transcripts (Supplementary Figure S4 and Supplementary Table S4). Comparison of the RNA-seq and proteomics datasets identified 133 proteins with reduced abundance, but with no detectable reduction in mRNA abundance (Supplementary Figure S4). We further inspected the data by selecting four candidate proteins with the highest fold change reported in the proteomics data (Table 1). From the selected candidate proteins, we discovered that three of the differentially expressed proteins reported in the proteomics data were not differentially expressed in the RNA-seq data, indicating that there is no change in transcripts levels leading to the fluctuation of their corresponding proteins. Only one protein candidate (AT2G45180.1) showed decreased fold change of mRNA, which correlates with the decrease of its corresponding protein reported in the proteomics data.

**Table 1:** Comparison between RNA-seq data and proteomics data of selected protein candidates. Four protein candidates with the highest fold change (*p-value* < 0.05) reported in the proteomics data were selected for further comparison.

| Gene ID | Protein | log2FC |  | Status |  |
| --- | --- | --- | --- | --- | --- |
|  |  | RNA-seq | Proteomics | RNA-seq | Proteomics |
| AT1G03540.1 | Pentatricopeptide repeat (PPR-like) superfamily protein | N/A | 3.12815 | non-differentially expressed | upregulated |
| AT2G45180.1 | Bifunctional inhibitor/lipid-transfer protein/seed storage 2S albumin superfamily protein | 2.012 | 0.56136 | downregulated | downregulated |
| AT2G39480.1 | P-glycoprotein 6 | N/A | 0.43734 | non-differentially expressed | downregulated |
| AT5G50160.1 | ferric reduction oxidase 8 | N/A | 0.40435 | non-differentially expressed | downregulated |

Finally, we performed a codon usage frequency analysis for this class of genes and detected no codon bias towards triplets read by both m^1^I and m^1^G modified tRNAs (Supplementary Figure S4).

### AtTRM5 is involved in leaf and root development and flowering time regulation

Before undertaking growth measurements, we grew wild-type Columbia, *trm5-1*, two complemented lines, and two overexpression lines together under long-day conditions, harvested and dried the seeds to minimise any maternal or environmental effects. To observe the early growth stages of seedlings, we grew the six lines (wild type, *trm5-1*, two complemented lines and two overexpression lines) on plates for 10 days (Figure 6A). The *trm5-1* seedlings were noticeably smaller than the wild type. In contrast, no clear differences were evident between wild type, the complemented and overexpression lines. To rule out the possibility that the reduced growth in *trm5-1* seedlings was due to slower germination, we measured the germination of *trm5-1* and wild type and no difference was observed (Supplementary Figure S2). Reduced growth of *trm5-1* roots was also evident on plate-grown plants (Figure 6B). Interestingly, *trm5-1* primary, lateral and total (primary + lateral) root lengths were reduced in *trm5-1* when compared to wild-type plants (Supplementary Figure S3). We also measured the lateral root number and found that *trm5-1* plants had reduced numbers when compared to the wild type (Supplementary Figure S3). In contrast, no differences in the root growth were evident upon comparison of the wild type and the complemented lines. In *TRM5* overexpression lines, primary and lateral root lengths were slightly longer than in the wild type (Figure 6B and Supplementary Figure S3).

**Figure 6.**
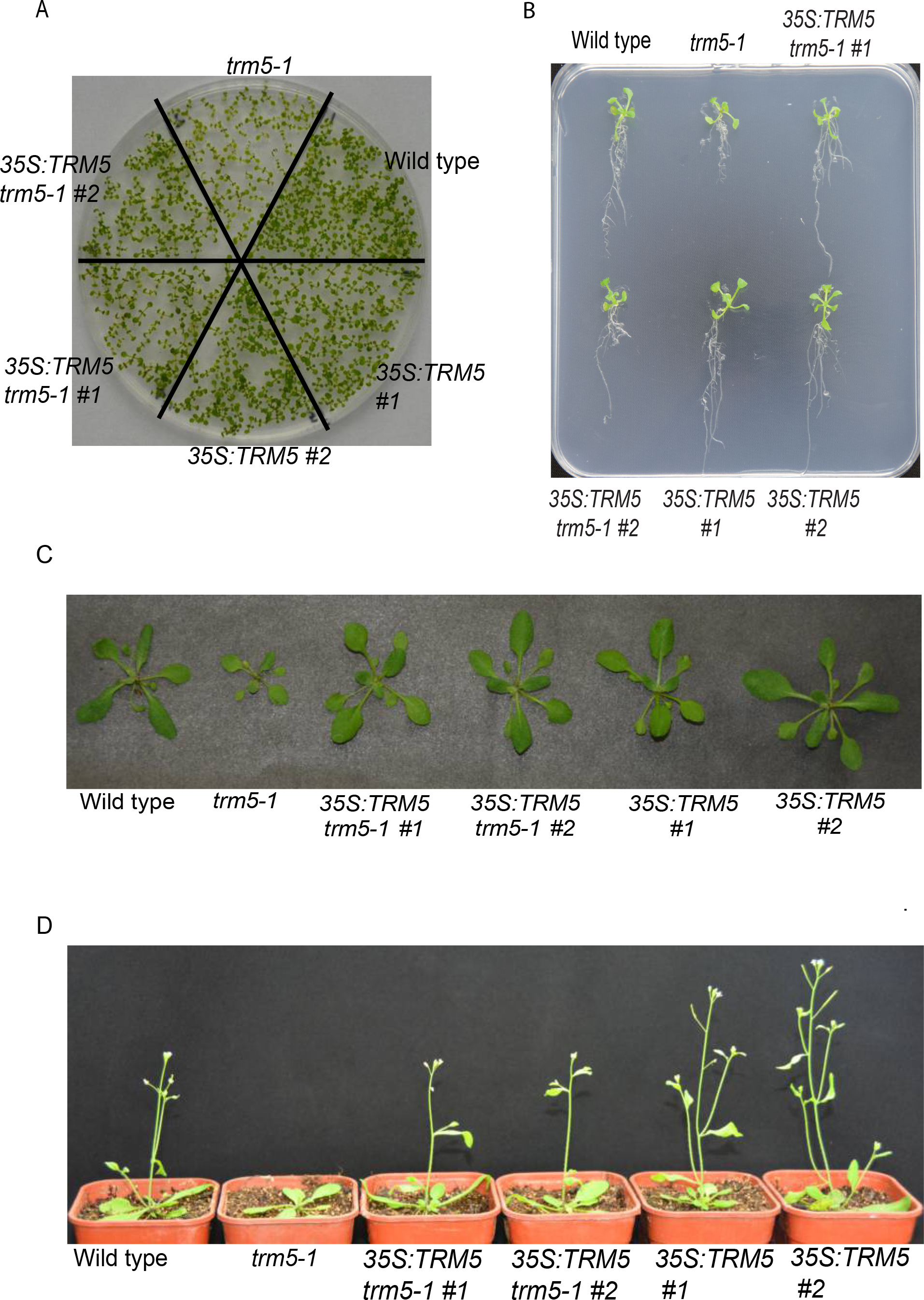
Phenotype analysis of *trm5*, complemented lines (35S:TRM5 *trm5-1*) and TRM5 overexpression lines (35S:TRM5). (**A**) Seedlings were sown on ½ MS media plates and grown for 7 days and photographed. (**B**) Seedlings of wild type, *trm5*, complemented lines (35S:TRM5 *trm5-1)*, TRM5 overexpression lines (35S:TRM5) were vertically grown on ½ MS medium for 10 days and then photographed. (**C**) Plants were grown on soil under long day photoperiods and photographed 15 days after germination. (**D**) Wild type, *trm5-1*, two complementing (35S:TRM5 *trm5-1*) and two overexpressing lines (35S:TRM5) were grown under long days and representative plants photographed at flowering.

At inflorescence emergence in wild-type plants grown under long days, we observed reduced rosette leaf numbers, smaller leaves and reduced fresh weight in *trm5-1* plants (Figure 6C and D; Supplementary Figure S2). Sectioning of the shoot apical meristems of wild type and *trm5-1* plants at wild-type floral transition, confirmed that *trm5* plants were later flowering (Supplementary Figure S2). We measured the flowering time of wild type, *trm5-1*, complemented, and overexpression lines under both short and long days and observed that mutants produced more rosette leaves and flowered later than the wild type (Figure 7A; Supplementary Figure S2). Plants overexpressing TRM5 flowered slightly earlier than wild type under both long and short-day conditions (Figure 6D; Supplementary Figure S2).

**Figure 7.**
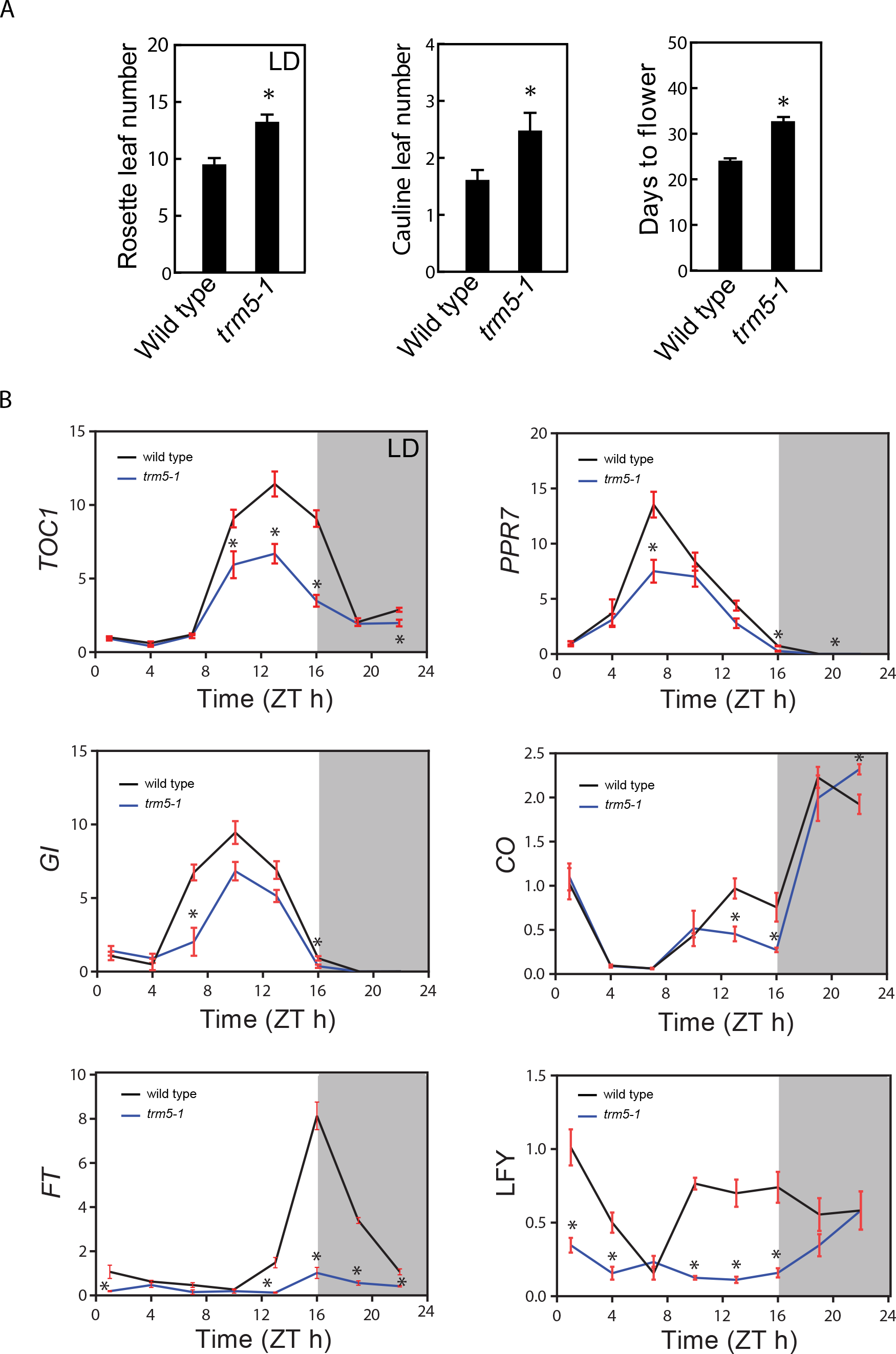
More leaves, slower flowering and impacted photosynthetic genes in trm5 mutant of *Arabidopsis thaliana*. (**A**) Rosette leaf number (long-day conditions), cauline leaf number and days to flower (short day conditions) of wild type and *trm5* mutant. (**B**) The mRNA abundance of circadian clock-related genes over a 24-hour period. 17-day-old seedlings of wild type, *trm5-1*, complemented lines (35S:TRM5 *trm5-1*), TRM5 overexpression lines (35S:TRM5) were grown on ½ MS medium for 10 days and then harvested every 3 hours. The expression levels of *TOC1, PRR7*, GI, CO, *FT*, and LFY were measured and normalized relative to *EF-1-α*. Data presented are means. Error bars are ± SE, n=3 biological replicates. An asterisk indicates a statistical difference (*P*<0.05) as determined by Student’s t-test. 0 hours is lights on and 16 hours is light off. Shaded area indicates night, whereas Zeitgeber time is abbreviated as ZT.

### AtTRM5 is involved in circadian clock and flowering time gene expression

To explore the molecular basis of delayed flowering in *trm5-1*, we measured the mRNA abundance of circadian clock and flowering time related genes by quantitative RT-PCR over a 24-hour period (Figure 7). In *trm5-1* plants, lower abundance of the clock genes *TIMING OF CAB EXPRESSION 1* (*TOC1*), and *Pseudo Response Regulators 7 (PPR7*), and the flowering time regulator genes *GIGANTEA (GI), CONSTANS (CO)* and *FLOWERING LOCUS T (FT)* were observed at ZT10 to ZT22. The reduced abundance of the flowering time regulators *GI*, *CO*, and *FT* in *trm5-1* mutants correlates with delayed flowering. As expected, the downstream floral meristem identity gene *LEAFY (LFY)* had lower abundance at almost all tested time points in the *trm5* mutant when compared to wild type (Figure 7B). Together these results support a role for TRM5 in plant growth, development, and flowering time regulation.

## Discussion

The discovery of m^1^G and m^1^I at position 37 in tRNAs of a wide range of eukaryotic and prokaryotic organisms underscores its importance as a key regulator of tRNA function and, presumably, translation (11, 20, 23). Previous studies in bacteria have shown that m^1^G37 is required for translational fidelity (20, 57, 58) and that mutations in the enzymes catalyzing m^1^G37 severely impact growth or cause lethal phenotypes (11, 23, 29). In eukaryotes, m^1^G37 modification requires the methyltransferase TRM5. Here we report that in plants, *trm5* mutants have a 60% reduction in m^1^G and m^1^I levels, display severely reduced growth and delayed transition to flowering. This is somewhat similar to human patients that are heterozygous for one mutant and one functional allele of *Trm5*. They show childhood failure to thrive and exercise intolerance symptoms (11).

With the 60% reduction of m^1^G in *trm5* plants, it is reasonable to anticipate a significant impact on protein translation due to the reduced stacking effect of m^1^G37 on base-pairing between position 36 and the first nucleoside in translated mRNA codons. We observed a reduction in abundance of proteins involved in ribosome biogenesis and photosynthesis, and this is likely to account for the observed reduced growth of the mutant plants. In the future, it would be interesting to test if translational errors, such as ribosome stalling or frame shifting, occur in *trm5* plants which would be similar to what has been previously reported in bacteria with reduced tRNA m^1^G (22).

We localized *Arabidopsis* TRM5 to the nucleus which is in contrast to the cytoplasmic and mitochondrial localisation in other eukaryotes: *Trypanosoma brucei, Homo sapiens* (HeLa cells) and *Saccharomyces cerevisiae* (11, 23, 29), although in one study *S. cerevisiae* Trm5 has also been localized exclusively to the nucleus (53). Interestingly, in yeast, Trm5 acts in the nucleus to form m^1^G on retrograde imported tRNA^Phe^ after initial export from the nucleus and subsequent splicing at the mitochondrial outer membrane. As retrograde tRNA import is conserved from yeast to vertebrates (59, 60), it is tempting to speculate that TRM5-mediated m^1^G formation occurs in retrograde imported tRNAs in *Arabidopsis*.

Direct comparison of RNA-seq and proteomics data has been challenging as variation in protein abundance in proteomics datasets can be confounded by multiple factors. Several factors can cause fluctuation in protein levels with no change of mRNA abundance: mRNA transcript abundance, translation rate, and translation resources availability (as reviewed in (as reviewed in (61)). This may explain the results of the initial comparison between our RNA-seq and proteomics data, and the seeming discrepancies in the derived sets of differentially expressed genes (Supplementary Figure S4). In our RNA-seq data, microtubule-related genes appeared to be significantly downregulated (AT1G21810, AT1G52410, AT2G28620, AT2G44190, AT3G23670, AT3G60840, AT3G63670, AT5G67270, AT4G14150, AT5G27000). However, we could not detect any differential expression of these proteins in our proteomics dataset. This is not unexpected, as it has been reported that protein and transcript abundances only rarely correlate in data analysis (61). However, it has been shown that transcript abundance can still be used to infer protein abundance (62). Among the selected four candidate proteins (Table 1), we could observe only one event of protein downregulation due to decreased transcript level: the lipid transfer protein (AT2G45180) which is involved in lipid transport in chloroplasts. Lipid transport is crucial in the formation of photosynthetic membranes in plants (63). As several lipids are important functional components of thylakoidal protein complexes involved in photosynthesis, the impacted lipid transport and lipid synthesis revealed by the GO term analysis may contribute to the observed dysregulation of photosynthesis-related genes. The remaining three candidate genes displayed no changes in transcript levels, but one candidate gene showed upregulation and the other two downregulation at the protein level. The exact function of pentatricopeptide repeat (PPR-like) superfamily protein (AT1G03540.1) is unknown, but the protein was suggested to play a significant role in post-transcriptional modification of tRNAs (64). The P-glycoprotein 6 (AT2G39480) belongs to the large P-glycoprotein family and is known to play a role in mediating auxin transport, and disturbed auxin transport is known to affect plant growth (65). Our proteomics data suggested that the auxin transport is affected (AT5G35735, AT3G07390, AT4G12980, AT5G35735) in the *trm5* mutant. Ferric reduction oxidase 8 (AT5G50160/AtFRO8) has been shown to participate in iron reduction and was implicated in leaf vein transport (66). We proposed that disturbed photosynthesis in *trm5* mutant plants is a secondary effect of dysregulated transport processes.

The role of post-transcriptional RNA modifications in tRNA and mRNA metabolism and their impact on plant growth and development in plants are only beginning to be elucidated. Here, we have described the TRM5-mediated m^1^G methylation in tRNAs and identified crucial links between this modification, photosynthesis, and plant growth. It appears likely that the many other tRNA modifications in plant tRNAs also play important roles in translation and/or translational regulation which remain to be discovered.

## Supporting information

## Data Access

The data sets supporting the results of this article are available in NCBI’s GEO database repository and are accessible through GEO Series accession number GSE114898. Proteomics raw data has been deposited on iProX, with under accession number IPX0001222000.

## Acknowledgements

We thank the staff at The Australian Cancer Research Foundation (ACRF) Cancer Genomics Facility, Adelaide. This research was partially supported by ARC grant FT130100525, partially funded an Australia-China Science and Research Fund grant ACSRF48187 awarded to I.S. and an APA awarded to PQ.N and J.L. This research was also partially supported by the National Natural Science Foundation (31570234) awarded to W.Z., W.Z. was supported by the Innovation Program of Chinese Academy of Agricultural Sciences and the Elite Youth Program of the Chinese Academy of Agricultural Science.

## Author’s contributions

Experiments were designed by I.S. and W.Z. Experiments were performed by R.D., PQ.N., I.S., Q.G., S.S., D.F., H.W., M.Z., T.D. and W.Z. Data analysis was performed by I.S., PQ.N., J.L., W.Z., and R.B. The manuscript was prepared and edited by PQ.N., Q.G., W.Z., I.S., and R.B. All authors read and approved the final manuscript.

## Competing interests

The authors declare that they have no competing interests.

