## Supplementary material for "*Arabidopsis* TRM5 encodes a nuclear-localised bifunctional tRNA guanine and inosine-N1-methyltransferase that is important for growth"

**Supplementary Figure S1. Bioinformatics characterization of TRM5 and related proteins.** (**A**) Multiple sequence alignment of TRM5 proteins from *Arabidopsis thaliana* (At), yeast (ScTrm5), humans (HsTrm5), *Drosophila melanogaster* (DmTrm5), *Pyrococcus horikoshii* (PhTYW2), and *Methanococcus jannaschii* (MjTYW2). Black shaded boxes are identical across all species. Light shaded boxes are similar and nearly conserved residues. Asterisk indicates catalytic important amino acids. (**B**) Multiple sequence alignment of yeast Trm5p, *Arabidopsis* TRM5 (At3g56120) and the two closest related proteins from *Arabidopsis*. Black shaded boxes are conserved in at least 2 sequences

**Supplementary Figure S2. Characterisation of growth and development in wild type, trm5 mutant, complemented and overexpression lines.** (**A**) Seeds (n=100) of wild type and *trm5-1* were sown on ½ MS plates, stratified at 4 $^{o}$C and then grown at 21 $^{o}$C under long day conditions for 32 hours. Germination was measured at 8, 16, 24 and 32 hours after shifting to 21 $^{o}$C. (**B**) Sections of the shoot apical meristems of wild type and *trm5-1* plants grown under long days for 14, 18 and 22 days. (**C**) The average fresh plant weight of long day grown plants. (**D**) The average rosette leaf number at flowering; (**E**) The average days to flowering under long days. (**F**) The rosette leaf number under short days. Data presented are means. Error bars are ± SE (n=16). NF= did not flower. An asterisk indicates a statistical difference (*P*<0.05) as determined by Student's t-test.

**Supplementary Figure S3. Root phenotypic analysis of seedlings.** Seedlings of wild type, *trm5*, complemented lines (35S:TRM5 *trm5-1*), TRM5 overexpression lines (35S:TRM5) were vertically grown on ½ MS medium for 10 days and then measured. (**A**) Total root length, (**C**B Primary root, (PR) length, (**C**) average lateral root (LR) length and (**D**) LR number were measured 10 days after germination. Data presented are means. Error bars are ± SE (n=10 plants).

.

**Supplementary Figure S4.** RNA-seq and proteomic analysis of wild type and *trm5-1*. (**A**) RNA was purified from 10-day-old seedlings of wild type (wt) and *trm5-1* (n=3). RNA-seq analysis was performed and differentially abundant transcripts were hierarchically clustered. (**B**) An upset plot showing positive overlapping mRNAs and proteins identified by RNA-seq and proteomics analysis. (**C**) Codon bias analysis of codons in the up and down regulated proteins identified by proteomics analysis.

**Supplementary Table S1.** Oligonucleotide primers used in this study.

**Supplementary Table S2.** Differentially abundant proteins identified by LC-MS/MS analysis of wild type and *trm5-1*.

**Supplementary Table S3.** Differentially expressed genes identified by RNA-seq analysis of wild type and *trm5-1*.

*

A

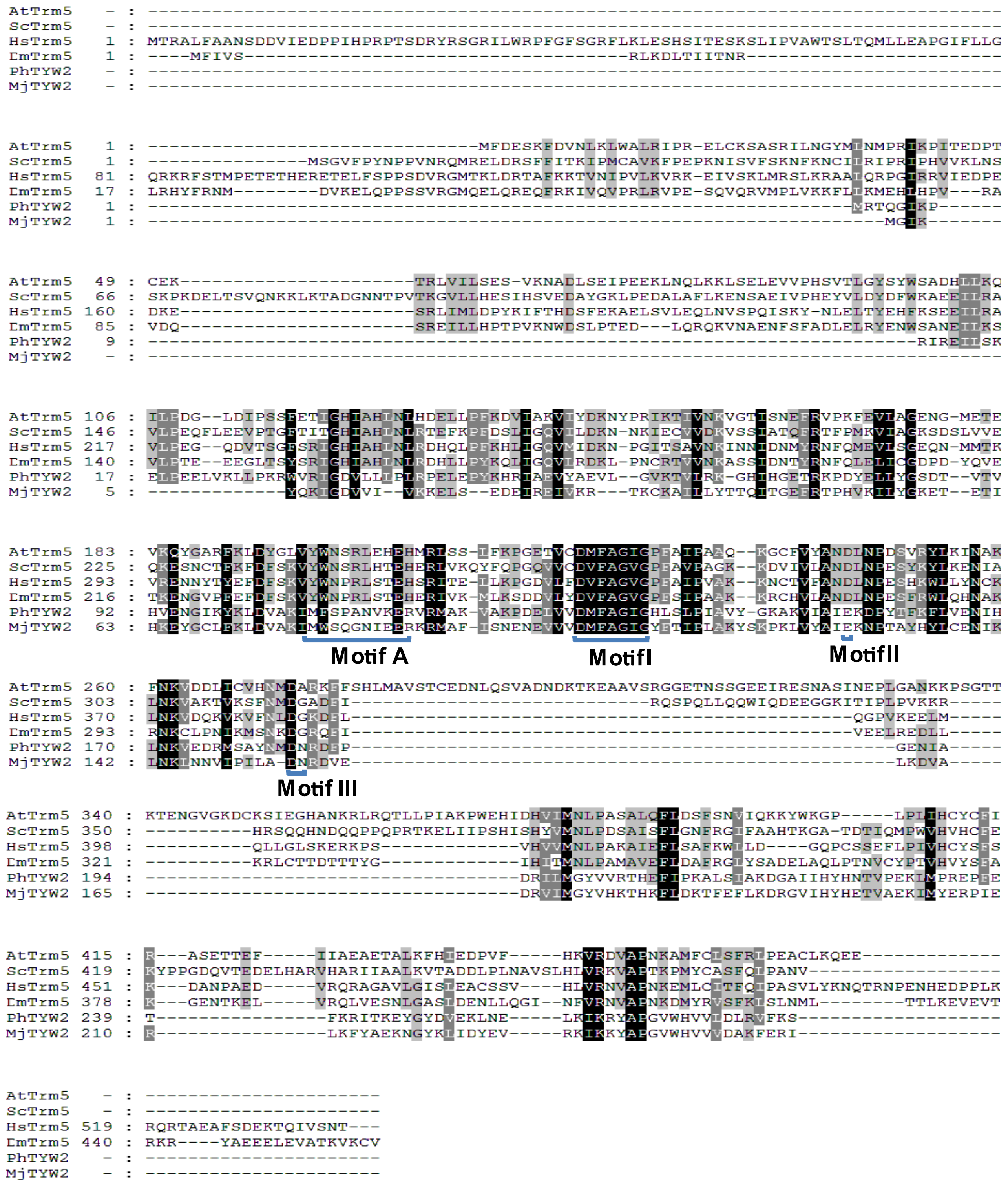

AtTrm5
ScTrm5
HsTrm5
DmTrm5
PhTYW2
MjTYW2
Motif A
MotifI
MotifII
Motif III


B

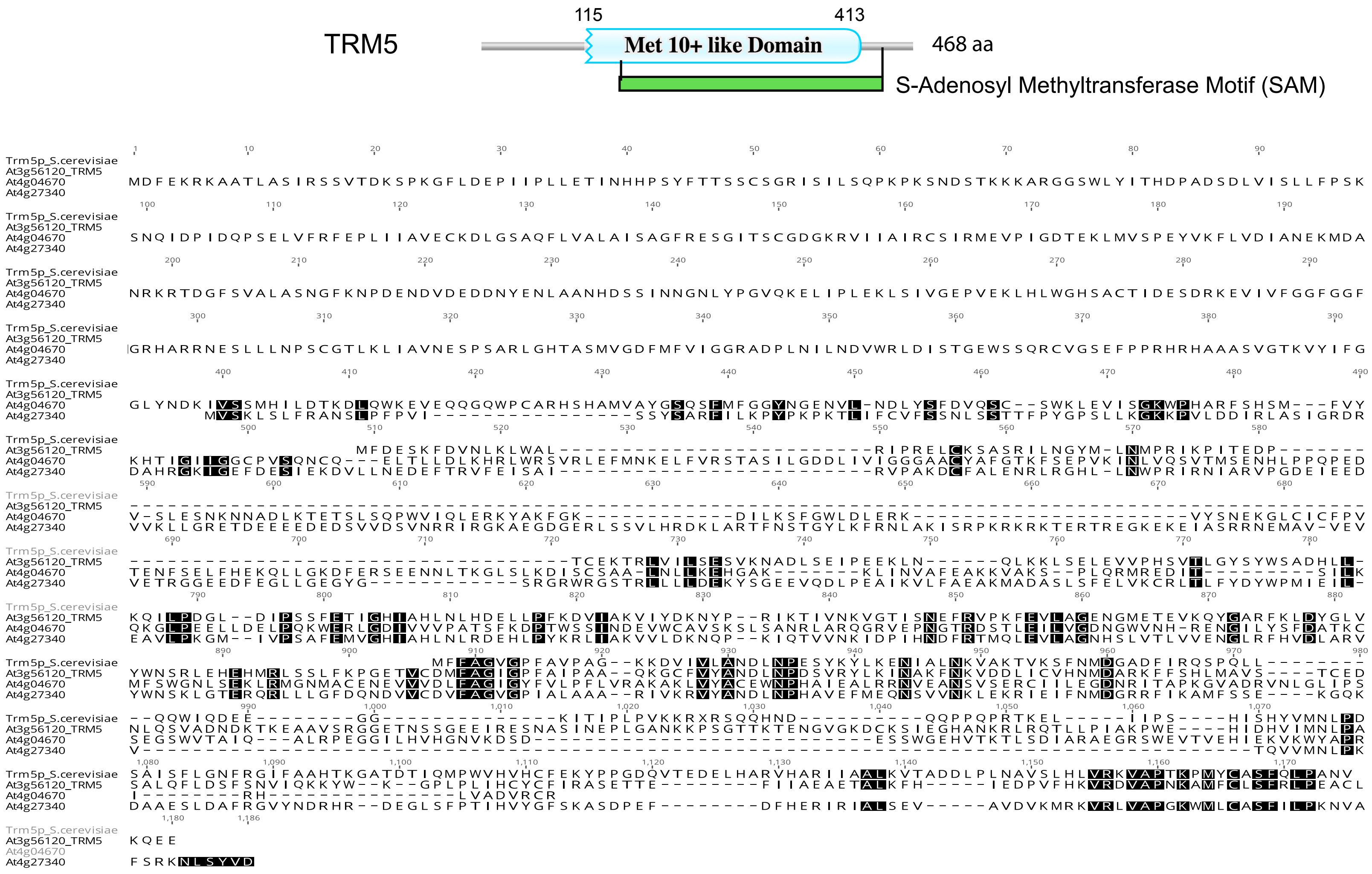

TRM5
115
413
Met 10+ like Domain
468 aa
S-Adenosyl Methyltransferase Motif (SAM)
Trm5p_S.cerevisiae
At3g56120_TRM5
At4g04670
At4g27340

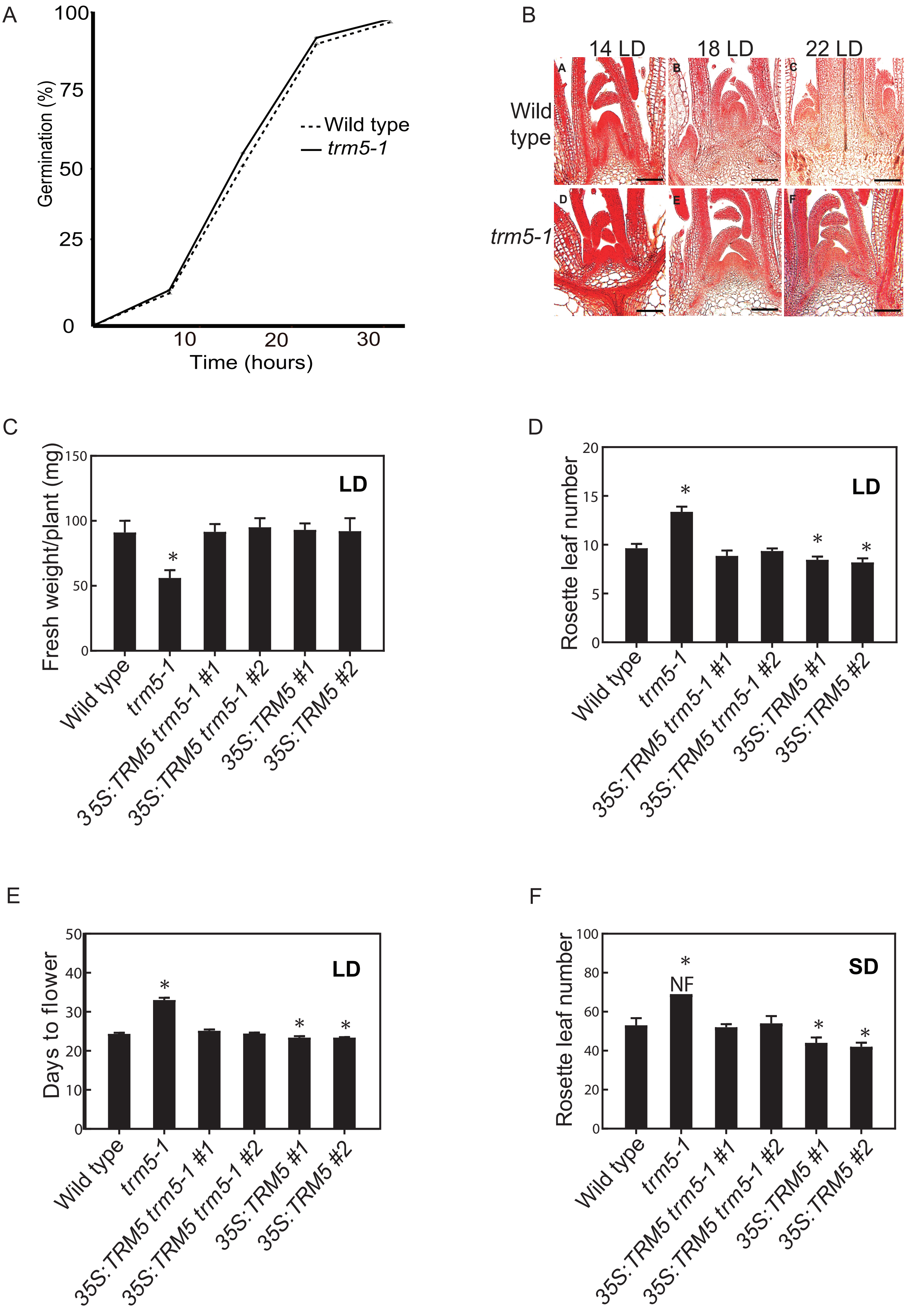
A
Germination (%)
100
75
50
25
0
10
20
30
Time (hours)
Wild type
trm5-1
B
14 LD
18 LD
22 LD
Wild type
trm5-1
C
Fresh weight/plant (mg)
LD
Wild type
trm5-1
35S:TRM5 trm5-1 #1
35S:TRM5 trm5-1 #2
35S:TRM5 #1
35S:TRM5 #2
D
Rosette leaf number
LD
E
Days to flower
LD
F
Rosette leaf number
SD
NF

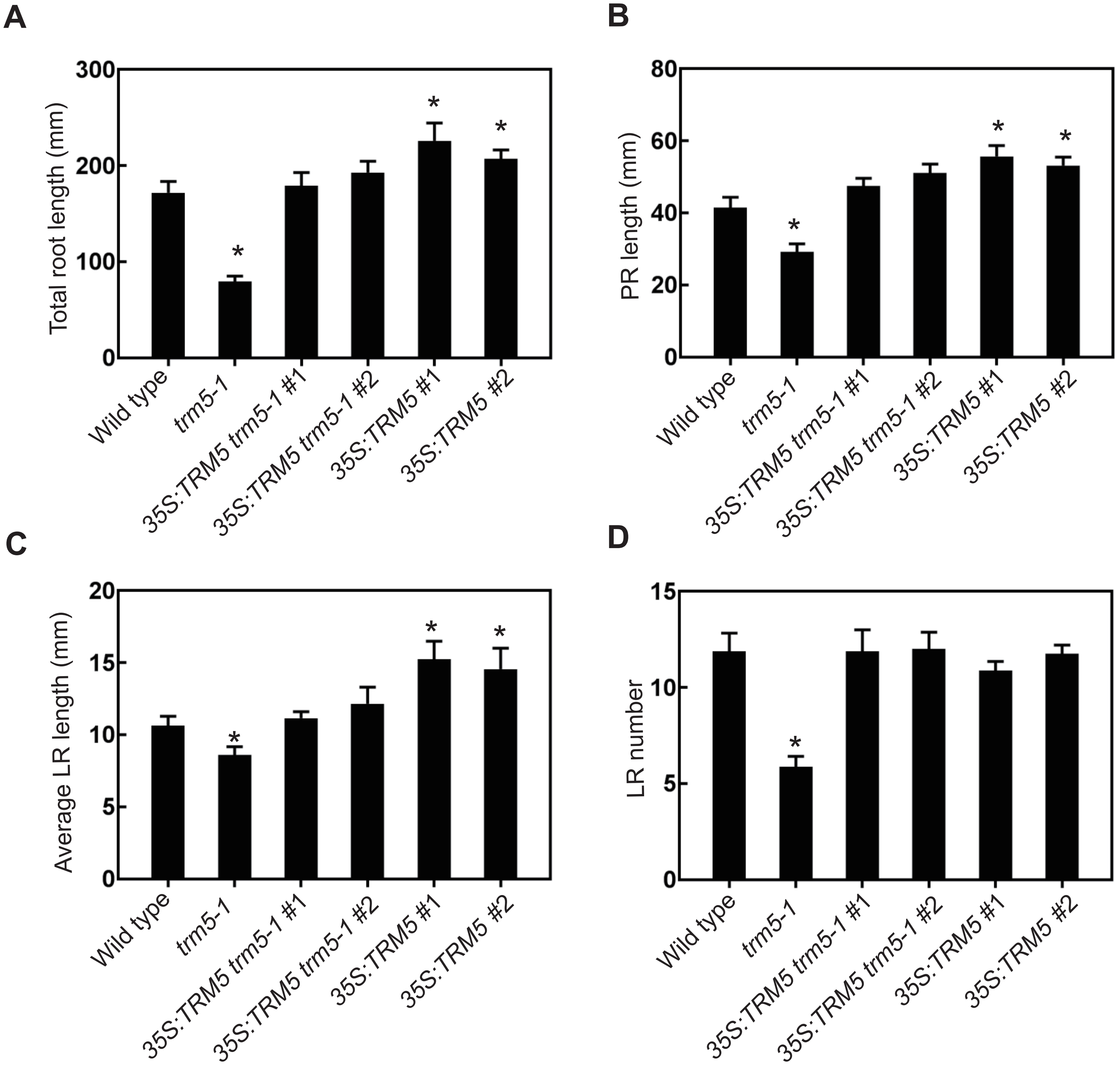

A
B
C
D
Total root length (mm)
PR length (mm)
Average LR length (mm)
LR number
0
100
200
300
20
40
60
80
5
10
15
20
*
Wild type
trm5-1
35S:TRM5 trm5-1 #1
35S:TRM5 trm5-1 #2
35S:TRM5 #1
35S:TRM5 #2

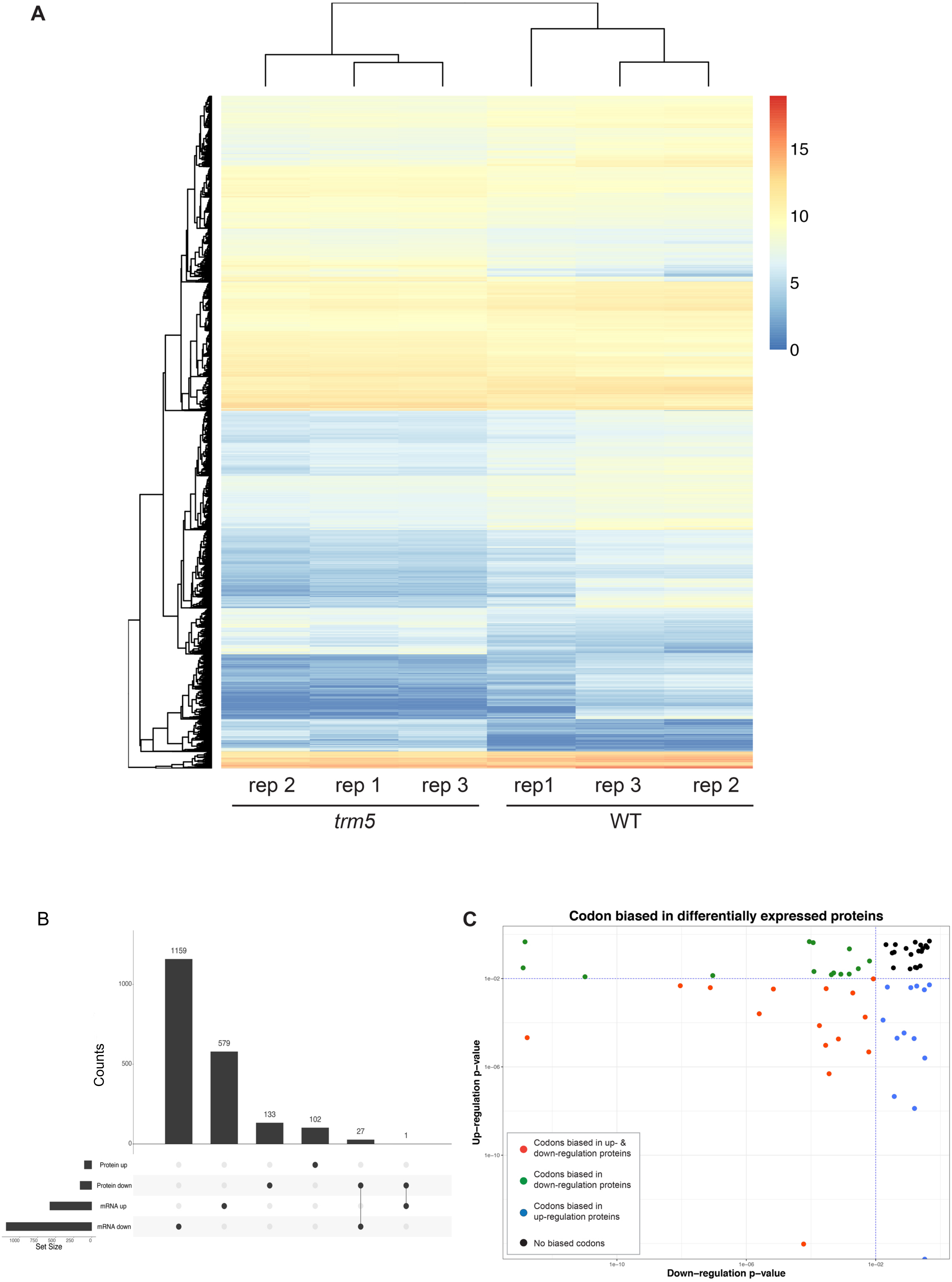
A
15
10
5
0
rep 2
rep 1
rep 3
rep1
rep 3
rep 2
trm5
WT
B
Counts
1159
579
133
102
27
1
1000
500
0
Protein up
Protein down
mRNA up
mRNA down
1000
750
500
250
0
Set Size
C
Codon biased in differentially expressed proteins
Up−regulation p−value
Down−regulation p−value
1e−02
1e−06
1e−10
1e−10
1e−06
1e−02
Codons biased in up- & down-regulation proteins
Codons biased in down-regulation proteins
Codons biased in up-regulation proteins
No biased codons

**Supplementary Table S1.** Oligonucleotide primers used in this study.

| **RT PCR Primers** | **Sequence** |
|---|---|
| TRM5-qRTPCR_F | 5'-ACACGCAAATAAGAGATTGAGAC-3' |
| TRM5-qRTPCR_R | 5'-TGGAACTTCAAAGCAGTTTCAGC-3' |
| FT- qRTPCR_F | 5'-GCTACAACTGGAACAACCTTTGGC-3' |
| FT- qRTPCR_R | 5'-TGAATTCCTGCAGTGGGACTTGG-3' |
| CO- qRTPCR_F | 5'-CTACAACGACAATGGTTCCATTAAC-3' |
| CO- qRTPCR_R | 5'-CAGGGTCAGGTTGTTGC-3' |
| GI- qRTPCR_F | 5'-GGGTAAATATGCTGCTGGAGA-3' |
| GI- qRTPCR_R | 5'-CAGTATGACACCAGCTCCATT-3' |
| SOC1- qRTPCR_F | 5'-AGCTGCAGAAAACGAGAAGCTCTCTG-3' |
| SOC1-qR | 5'-GGGCTACTCTCTTCATCACCTCTTCC-3' |
| LFY- qRTPCR_F | 5'-ATCGCTTGTCGTCATGGCTG-3' |
| LFY- qRTPCR_R | 5'-GCAACCGCATTGTTCCGCTC-3' |
| AP1- qRTPCR_F | 5'-CAGCAGCACCAAATCCAGC-3' |
| AP1- qRTPCR_R | 5'-GAGCCTAGCCACTATTTATATG-3' |
| CCA1- qRTPCR_F | 5'-CCTCAA ACTTCA GAGTCCAATGC-3' |
| CCA1- qRTPCR_R | 5'-GACCCTCGTCAGACACAGACTTC-3' |
| LHY- qRTPCR_F | 5'-GAAGTCTCCGAAGAGGGTCG-3' |
| LHY- qRTPCR_R | 5'-TATTCACATTCTCTGCCACTTGAG-3' |
| TOC1- qRTPCR_F | 5'-GCTATGAACAGAAGTAAAGATTCG-3' |
| TOC1- qRTPCR_R | 5'-GGATATCCCGTCATTCCATTCGGA-3' |
| PRR7- qRTPCR_F | 5'-CTGCACTCGTTATATCGTTACTG-3' |
| PRR7- qRTPCR_R | 5'-GGCATGATCACCTCTGTTAG-3' |

| | |
|---|---|
| EF-1-α- qRTPCR_F | 5'-TGAGCACGCTCTTCTTGCTTTCA-3' |
| EF-1-α- qRTPCR_R | 5'-GGTGGTGGCATCCATCTTGTTACA-3' |
| **Primer sequences for T-DNA insertion identification** | |
| TRM5-1_022617RP | 5'-TTGCGAGGATAATTTGCAATC-3' |
| TRM5-1_022617LP | 5'-CCTTGTTTGGAGCAACATCTC-3' |
| TRM5-2_ 032376RP | 5'- CATGATGAGCTCCTTCCATTCAAAG-3' |
| TRM5-2_ 032376LP | 5'- CGCATGTGCTCATGTTCCAATC-3' |
| LB-SALK-b1.3 | 5'-ATTTTGCCGATTTCGGAAC-3' |
| **Primer sequences for *TRM5*** | |
| TRM5-CDSf1 | 5'-CACCATGTTTGATGAAAGCAAGTTCGATGTC-3' |
| TRM5-CDSr1 | 5'-CTCTTCTTGCTTCAAGCATGCTTC-3' |
| TRM5Pro_F1 | 5'-AGTGGCTGATCTGAGGGACGGAAGAGG-3' |
| TRM53'_R1 | 5'-GAGAGCTGCAGAAGACGCCATTGGAAG-3' |
| **Primer sequences for $tRNA^{Ala}$ (AGC) amplification** | |
| tRNA-Ala-f | 5'-GGGGATGTAGCTCAGATGGTAG-3' |
| tRNA -Ala-f | 5'-TGGTGGAGATGCGGGGTATC-3' |

**Supplementary Table S2.** Differentially abundant proteins identified by LC-MS/MS analysis of

**Up-regulated in *trm5-1***

| Accession | Description |
|---|---|
| AT1G03540.1 | Pentatricopeptide repeat (PPR-like) superfamily protein |
| AT4G02520.1 | glutathione S-transferase PHI 2 glutathione S-transferase PHI 2 |
| AT3G06730.1 | Thioredoxin z Thioredoxin z Thioredoxin z Thioredoxin z Thioredoxin z |
| AT2G41100.1 | Calcium-binding EF hand family protein |
| AT5G08040.1 | mitochondrial import receptor subunit TOM5 homolog |
| AT4G26130.1 | unknown protein; FUNCTIONS IN: molecular_function unknown; INVOLVED IN |
| AT2G30860.2 | glutathione S-transferase PHI 9 glutathione S-transferase PHI 9 |
| AT5G16930.1 | AAA-type ATPase family protein AAA-type ATPase family protein |
| AT2G40360.1 | Transducin/WD40 repeat-like superfamily protein |
| AT3G20410.1 | calmodulin-domain protein kinase 9 calmodulin-domain protein kinase 9 |
| AT3G23620.1 | Ribosomal RNA processing Brix domain protein |
| AT2G20530.1 | prohibitin 6 prohibitin 6 prohibitin 6 prohibitin 6 prohibitin 6 prohibitin 6 |
| AT1G14880.1 | PLANT CADMIUM RESISTANCE 1 PLANT CADMIUM RESISTANCE 1 |
| AT2G21060.1 | glycine-rich protein 2B glycine-rich protein 2B glycine-rich protein 2B |
| AT5G42790.1 | proteasome alpha subunit F1 proteasome alpha subunit F1 |
| AT4G25100.1 | Fe superoxide dismutase 1 Fe superoxide dismutase 1 |
| AT4G32720.2 | La protein 1 La protein 1 La protein 1 La protein 1 La protein 1 La protein 1 |
| AT1G21250.1 | cell wall-associated kinase cell wall-associated kinase |
| AT3G23830.1 | glycine-rich RNA-binding protein 4 glycine-rich RNA-binding protein 4 |
| AT3G11450.1 | DnaJ domain ;Myb-like DNA-binding domain |
| AT3G62200.1 | Putative endonuclease or glycosyl hydrolase |
| AT4G27990.1 | YGGT family protein YGGT family protein YGGT family protein |
| AT1G25260.1 | Ribosomal protein L10 family protein Ribosomal protein L10 family protein |
| AT5G35735.1 | Auxin-responsive family protein Auxin-responsive family protein |
| AT1G79390.1 | unknown protein; Has 30201 Blast hits to 17322 proteins in 780 species: Arch |
| AT2G44310.1 | Calcium-binding EF-hand family protein |
| AT3G28900.1 | Ribosomal protein L34e superfamily protein |
| ATCG00560.1 | photosystem II reaction center protein L |
| AT1G02930.1 | glutathione S-transferase 6 glutathione S-transferase 6 |
| AT4G08310.1 | FUNCTIONS IN: molecular_function unknown; INVOLVED IN: biological_proce |
| AT4G12610.1 | transcription activators;DNA binding;RNA polymerase II transcription factors;ca |
| AT2G38530.1 | lipid transfer protein 2 lipid transfer protein 2 lipid transfer protein 2 |
| AT4G38740.1 | rotamase CYP 1 rotamase CYP 1 rotamase CYP 1 rotamase CYP 1 |
| AT3G04120.1 | glyceraldehyde-3-phosphate dehydrogenase C subunit 1 |
| AT3G06680.2 | Ribosomal L29e protein family Ribosomal L29e protein family |
| AT4G12480.1 | Bifunctional inhibitor/lipid-transfer protein/seed storage 2S albumin superfamily |
| AT3G05500.1 | Rubber elongation factor protein (REF) |
| AT2G40430.3 | FUNCTIONS IN: molecular_function unknown; INVOLVED IN: biological_proce |
| AT4G12790.1 | P-loop containing nucleoside triphosphate hydrolases superfamily protein |
| AT3G46430.1 | unknown protein; FUNCTIONS IN: molecular_function unknown; INVOLVED IN |
| AT2G05840.1 | 20S proteasome subunit PAA2 20S proteasome subunit PAA2 |
| AT1G54270.1 | eif4a-2 eif4a-2 eif4a-2 eif4a-2 eif4a-2 eif4a-2 eif4a-2 eif4a-2 eif4a-2 eif4a-2 |

| | |
|---|---|
| AT1G07920.1 | GTP binding Elongation factor Tu family protein |
| AT1G48920.1 | nucleolin like 1 nucleolin like 1 nucleolin like 1 nucleolin like 1 nucleolin like 1 |
| AT1G51720.1 | Amino acid dehydrogenase family protein |
| AT3G07050.1 | GTP-binding family protein GTP-binding family protein |
| AT4G28510.1 | prohibitin 1 prohibitin 1 prohibitin 1 prohibitin 1 prohibitin 1 prohibitin 1 |
| AT2G33180.1 | unknown protein; FUNCTIONS IN: molecular_function unknown; INVOLVED IN |
| AT5G36290.1 | Uncharacterized protein family (UPF0016) |
| AT1G80750.1 | Ribosomal protein L30/L7 family protein |
| AT1G05410.2 | Protein of unknown function (DUF1423) |
| AT4G21105.2 | cytochrome-c oxidases;electron carriers |
| AT1G51760.1 | peptidase M20/M25/M40 family protein |
| AT1G05010.1 | ethylene-forming enzyme ethylene-forming enzyme ethylene-forming enzyme |
| AT4G16250.1 | phytochrome D phytochrome D phytochrome D phytochrome D |
| AT1G21720.1 | proteasome beta subunit C1 proteasome beta subunit C1 |
| AT5G40770.1 | prohibitin 3 prohibitin 3 prohibitin 3 prohibitin 3 prohibitin 3 prohibitin 3 |
| AT3G44750.2 | histone deacetylase 3 histone deacetylase 3 histone deacetylase 3 |
| AT5G11670.1 | NADP-malic enzyme 2 NADP-malic enzyme 2 NADP-malic enzyme 2 |
| AT5G57290.2 | 60S acidic ribosomal protein family 60S acidic ribosomal protein family |
| AT3G52730.1 | ubiquinol-cytochrome C reductase UQCRX/QCR9-like family protein |
| AT5G64400.1 | CONTAINS InterPro DOMAIN/s: CHCH (InterPro:IPR010625); BEST Arabidops |
| AT1G69250.2 | Nuclear transport factor 2 (NTF2) family protein with RNA binding (RRM-RBD-R |
| AT3G57520.3 | seed imbibition 2 seed imbibition 2 seed imbibition 2 seed imbibition 2 |
| AT5G22650.2 | histone deacetylase 2B histone deacetylase 2B histone deacetylase 2B |
| AT1G03860.1 | prohibitin 2 prohibitin 2 prohibitin 2 prohibitin 2 prohibitin 2 prohibitin 2 |
| AT4G15770.1 | RNA binding RNA binding RNA binding RNA binding RNA binding |
| AT3G13920.1 | eukaryotic translation initiation factor 4A1 |
| AT5G58420.1 | Ribosomal protein S4 (RPS4A) family protein |
| AT1G78370.1 | glutathione S-transferase TAU 20 glutathione S-transferase TAU 20 |
| AT1G14060.1 | GCK domain-containing protein GCK domain-containing protein |
| AT1G29030.1 | Apoptosis inhibitory protein 5 (API5) Apoptosis inhibitory protein 5 (API5) |
| AT1G07070.1 | Ribosomal protein L35Ae family protein |
| AT1G49340.2 | Phosphatidylinositol 3- and 4-kinase family protein |
| AT3G16400.1 | nitrile specifier protein 1 nitrile specifier protein 1 nitrile specifier protein 1 |
| AT4G00810.1 | 60S acidic ribosomal protein family 60S acidic ribosomal protein family |
| AT2G34357.1 | ARM repeat superfamily protein ARM repeat superfamily protein |
| AT1G01100.3 | 60S acidic ribosomal protein family 60S acidic ribosomal protein family |
| AT4G30800.1 | Nucleic acid-binding, OB-fold-like protein |
| AT1G75950.1 | S phase kinase-associated protein 1 S phase kinase-associated protein 1 |
| AT2G43780.1 | unknown protein; Has 30 Blast hits to 30 proteins in 11 species: Archae - 0; Ba |
| AT3G55620.1 | Translation initiation factor IF6 Translation initiation factor IF6 |
| AT1G66240.1 | homolog of anti-oxidant 1 homolog of anti-oxidant 1 |
| AT4G31500.1 | cytochrome P450, family 83, subfamily B, polypeptide 1 |
| AT1G21130.1 | O-methyltransferase family protein O-methyltransferase family protein |
| AT2G29530.1 | Tim10/DDP family zinc finger protein Tim10/DDP family zinc finger protein |
| AT5G16070.1 | TCP-1/cpn60 chaperonin family protein |
| AT4G13180.1 | NAD(P)-binding Rossmann-fold superfamily protein |

| | |
|---|---|
| AT1G07830.1 | ribosomal protein L29 family protein ribosomal protein L29 family protein |
| AT1G35670.1 | calcium-dependent protein kinase 2 calcium-dependent protein kinase 2 |
| AT3G22660.1 | rRNA processing protein-related rRNA processing protein-related |
| AT1G29250.1 | Alba DNA/RNA-binding protein Alba DNA/RNA-binding protein |
| AT3G16780.1 | Ribosomal protein L19e family protein Ribosomal protein L19e family protein |
| AT5G27120.1 | NOP56-like pre RNA processing ribonucleoprotein |
| AT1G26400.1 | FAD-binding Berberine family protein FAD-binding Berberine family protein |
| AT2G42680.1 | multiprotein bridging factor 1A multiprotein bridging factor 1A |
| AT3G11200.1 | alfin-like 2 alfin-like 2 alfin-like 2 alfin-like 2 alfin-like 2 alfin-like 2 alfin-like 2 |
| AT3G07810.1 | RNA-binding (RRM/RBD/RNP motifs) family protein |
| AT1G16000.1 | unknown protein; BEST Arabidopsis thaliana protein match is: unknown protei |
| AT3G03920.1 | H/ACA ribonucleoprotein complex, subunit Gar1/Naf1 protein |
| AT5G02450.1 | Ribosomal protein L36e family protein Ribosomal protein L36e family protein |
| AT5G46030.1 | unknown protein; CONTAINS InterPro DOMAIN/s: Protein of unknown functior |

**Down-regulated in *trm5-1***

| Accession | Description |
|---|---|
| AT5G65840.1 | Thioredoxin superfamily protein |
| AT3G52340.1 | sucrose-6F-phosphate phosphohydrolase 2 |
| AT1G50320.1 | thioredoxin X |
| AT3G22060.1 | Receptor-like protein kinase-related family protein |
| AT2G32540.1 | cellulose synthase-like B4 |
| AT1G69295.1 | plasmodesmata callose-binding protein 4 |
| AT1G62600.1 | Flavin-binding monooxygenase family protein |
| AT1G09750.1 | Eukaryotic aspartyl protease family protein |
| AT1G76730.1 | NagB/RpiA/CoA transferase-like superfamily protein |
| AT3G56650.1 | Mog1/PsbP/DUF1795-like photosystem II reaction center PsbP family protein |
| ATCG00350.1 | Photosystem I, PsaA/PsaB protein |
| AT5G54270.1 | light-harvesting chlorophyll B-binding protein 3 |
| AT5G42650.1 | allene oxide synthase |
| AT3G53650.1 | Histone superfamily protein |
| AT5G52920.1 | plastidic pyruvate kinase beta subunit 1 |
| AT2G30170.2 | Protein phosphatase 2C family protein |
| AT2G20890.1 | photosystem II reaction center PSB29 protein |
| AT5G13410.1 | FKBP-like peptidyl-prolyl cis-trans isomerase family protein |
| AT2G34430.1 | light-harvesting chlorophyll-protein complex II subunit B1 |
| AT1G26230.2 | TCP-1/cpn60 chaperonin family protein |
| AT3G10060.1 | FKBP-like peptidyl-prolyl cis-trans isomerase family protein |
| AT1G29700.1 | Metallo-hydrolase/oxidoreductase superfamily protein |
| AT4G37980.1 | elicitor-activated gene 3-1 |
| AT1G55210.1 | Disease resistance-responsive (dirigent-like protein) family protein |
| AT5G42070.1 | unknown protein; FUNCTIONS IN: molecular_function unknown; INVOLVED IN |
| AT2G47400.1 | CP12 domain-containing protein 1 |
| AT1G78680.2 | gamma-glutamyl hydrolase 2 |
| AT5G14430.1 | S-adenosyl-L-methionine-dependent methyltransferases superfamily protein |
| AT4G26530.1 | Aldolase superfamily protein |

| | |
|---|---|
| AT1G73600.1 | S-adenosyl-L-methionine-dependent methyltransferases superfamily protein |
| AT3G51820.1 | UbiA prenyltransferase family protein |
| AT2G23600.1 | acetone-cyanohydrin lyase |
| AT4G25080.4 | magnesium-protoporphyrin IX methyltransferase |
| AT3G61870.1 | unknown protein; FUNCTIONS IN: molecular_function unknown; INVOLVED IN |
| AT1G10200.1 | GATA type zinc finger transcription factor family protein |
| AT3G23450.1 | unknown protein; FUNCTIONS IN: molecular_function unknown; INVOLVED IN |
| AT1G15820.1 | light harvesting complex photosystem II subunit 6 |
| AT1G03475.1 | Coproporphyrinogen III oxidase |
| AT5G02160.1 | unknown protein; FUNCTIONS IN: molecular_function unknown; INVOLVED IN |
| AT1G31190.1 | myo-inositol monophosphatase like 1 |
| AT4G12980.1 | Auxin-responsive family protein |
| AT4G24350.2 | Phosphorylase superfamily protein |
| AT2G10940.1 | Bifunctional inhibitor/lipid-transfer protein/seed storage 2S albumin superfamily |
| AT4G28750.1 | Photosystem I reaction centre subunit IV / PsaE protein |
| AT1G80030.1 | Molecular chaperone Hsp40/DnaJ family protein |
| AT4G15800.1 | ralf-like 33 |
| AT2G20830.1 | transferases;folic acid binding |
| AT4G23400.1 | plasma membrane intrinsic protein 1;5 |
| AT1G54780.1 | thylakoid lumen 18.3 kDa protein |
| AT4G09010.1 | ascorbate peroxidase 4 |
| AT5G44130.1 | FASCICLIN-like arabinogalactan protein 13 precursor |
| AT3G27840.1 | ribosomal protein L12-B |
| AT4G36810.1 | geranylgeranyl pyrophosphate synthase 1 |
| AT5G62350.1 | Plant invertase/pectin methylesterase inhibitor superfamily protein |
| AT5G25460.1 | Protein of unknown function, DUF642 |
| AT1G26761.1 | Arabinanase/levansucrase/invertase |
| AT4G38970.1 | fructose-bisphosphate aldolase 2 |
| AT3G20820.1 | Leucine-rich repeat (LRR) family protein |
| AT4G35090.1 | catalase 2 |
| ATCG00140.1 | ATP synthase subunit C family protein |
| AT3G60130.3 | beta glucosidase 16 |
| AT3G06000.1 | RNI-like superfamily protein |
| AT1G78260.2 | RNA-binding (RRM/RBD/RNP motifs) family protein |
| AT5G14780.1 | formate dehydrogenase |
| AT3G04290.1 | Li-tolerant lipase 1 |
| AT1G31220.1 | Formyl transferase |
| AT5G21430.1 | Chaperone DnaJ-domain superfamily protein |
| AT3G09580.1 | FAD/NAD(P)-binding oxidoreductase family protein |
| AT1G52230.1 | photosystem I subunit H2 |
| AT3G47470.1 | light-harvesting chlorophyll-protein complex I subunit A4 |
| AT1G31330.1 | photosystem I subunit F |
| AT2G39080.1 | NAD(P)-binding Rossmann-fold superfamily protein |
| AT3G45140.1 | lipoxygenase 2 |
| ATCG00150.1 | ATPase, F0 complex, subunit A protein |
| AT5G57170.1 | Tautomerase/MIF superfamily protein |

| | |
|---|---|
| AT5G47880.1 | eukaryotic release factor 1-1 |
| AT1G78915.1 | Tetratricopeptide repeat (TPR)-like superfamily protein |
| AT4G05180.1 | photosystem II subunit Q-2 |
| AT1G65590.1 | beta-hexosaminidase 3 |
| AT3G14210.1 | epithiospecifier modifier 1 |
| AT1G29660.1 | GDSL-like Lipase/Acylhydrolase superfamily protein |
| AT3G51600.1 | lipid transfer protein 5 |
| AT5G27670.1 | histone H2A 7 |
| AT4G39710.2 | FK506-binding protein 16-2 |
| AT4G00165.1 | Bifunctional inhibitor/lipid-transfer protein/seed storage 2S albumin superfamily |
| AT4G14030.1 | selenium-binding protein 1 |
| AT1G03870.1 | FASCICLIN-like arabinoogalactan 9 |
| AT5G46290.2 | 3-ketoacyl-acyl carrier protein synthase I |
| AT3G15520.1 | Cyclophilin-like peptidyl-prolyl cis-trans isomerase family protein |
| AT1G54350.1 | ABC transporter family protein |
| AT1G60550.1 | enoyl-CoA hydratase/isomerase D |
| AT3G63160.1 | FUNCTIONS IN: molecular_function unknown; INVOLVED IN: biological_proce |
| AT5G54760.1 | Translation initiation factor SUI1 family protein |
| AT5G24165.1 | unknown protein; FUNCTIONS IN: molecular_function unknown; INVOLVED IN |
| AT3G47630.1 | FUNCTIONS IN: molecular_function unknown; INVOLVED IN: biological_proce |
| AT3G26570.2 | phosphate transporter 2;1 |
| AT3G28300.1 | Protein of unknown function (DUF677) |
| AT1G09340.1 | chloroplast RNA binding |
| AT5G36870.1 | glucan synthase-like 9 |
| AT1G61520.2 | photosystem I light harvesting complex gene 3 |
| AT3G54890.4 | photosystem I light harvesting complex gene 1 |
| AT3G07390.1 | auxin-responsive family protein |
| AT1G64680.1 | unknown protein; BEST Arabidopsis thaliana protein match is: unknown protei |
| AT4G19170.1 | nine-cis-epoxycarotenoid dioxygenase 4 |
| AT3G54140.1 | peptide transporter 1 |
| AT2G05310.1 | unknown protein; FUNCTIONS IN: molecular_function unknown; INVOLVED IN |
| AT2G39730.2 | rubisco activase |
| AT4G12730.1 | FASCICLIN-like arabinogalactan 2 |
| AT5G65685.2 | UDP-Glycosyltransferase superfamily protein |
| AT4G16500.1 | Cystatin/monellin superfamily protein |
| AT2G25800.1 | Protein of unknown function (DUF810) |
| AT3G15840.4 | post-illumination chlorophyll fluorescence increase |
| AT1G03130.1 | photosystem I subunit D-2 |
| AT2G43360.1 | Radical SAM superfamily protein |
| AT1G21770.1 | Acyl-CoA N-acyltransferases (NAT) superfamily protein |
| AT2G33330.1 | plasmodesmata-located protein 3 |
| AT1G68100.1 | ZIP metal ion transporter family |
| AT4G16015.1 | Cysteine/Histidine-rich C1 domain family protein |
| AT5G24490.1 | 30S ribosomal protein, putative |
| AT3G59690.1 | IQ-domain 13 |
| AT5G15530.1 | biotin carboxyl carrier protein 2 |

| | |
|---|---|
| AT2G38710.1 | AMMECR1 family |
| AT2G22990.2 | sinapoylglucose 1 |
| AT3G46630.1 | Protein of unknown function (DUF3223) |
| AT1G25230.1 | Calcineurin-like metallo-phosphoesterase superfamily protein |
| AT1G31690.1 | Copper amine oxidase family protein |
| AT3G28270.2 | Protein of unknown function (DUF677) |
| AT1G52190.1 | Major facilitator superfamily protein |
| AT1G51890.2 | Leucine-rich repeat protein kinase family protein |
| AT1G21500.1 | unknown protein; Has 29 Blast hits to 29 proteins in 12 species: Archae - 0; B[illegible] |
| AT3G28500.1 | 60S acidic ribosomal protein family |
| AT1G53840.1 | pectin methylesterase 1 |
| AT1G69740.1 | Aldolase superfamily protein |
| AT5G24420.1 | 6-phosphogluconolactonase 5 |
| AT5G11680.1 | FUNCTIONS IN: molecular_function unknown; INVOLVED IN: biological_proce |
| AT4G30950.1 | fatty acid desaturase 6 |
| AT2G27290.1 | Protein of unknown function (DUF1279) |
| AT2G02100.1 | low-molecular-weight cysteine-rich 69 |
| AT1G36730.1 | Translation initiation factor IF2/IF5 |
| AT4G12310.1 | cytochrome P450, family 706, subfamily A, polypeptide 5 |
| AT1G22630.1 | unknown protein; LOCATED IN: chloroplast; EXPRESSED IN: 21 plant structu |
| AT5G15970.1 | stress-responsive protein (KIN2) / stress-induced protein (KIN2) / cold-responsi |
| AT3G52090.1 | DNA-directed RNA polymerase, RBP11-like |
| AT1G51400.1 | Photosystem II 5 kD protein |
| AT4G02770.1 | photosystem I subunit D-1 |
| AT1G20340.1 | Cupredoxin superfamily protein |
| AT1G74670.1 | Gibberellin-regulated family protein |
| AT1G23090.1 | sulfate transporter 91 |
| AT3G08770.1 | lipid transfer protein 6 |
| AT5G23820.1 | MD-2-related lipid recognition domain-containing protein |
| AT2G43550.1 | Scorpion toxin-like knottin superfamily protein |
| AT1G03260.1 | SNARE associated Golgi protein family |
| AT1G14380.1 | IQ-domain 28 |
| AT1G33270.2 | Acyl transferase/acyl hydrolase/lysophospholipase superfamily protein |
| AT5G33280.1 | Voltage-gated chloride channel family protein |
| AT1G16300.1 | glyceraldehyde-3-phosphate dehydrogenase of plastid 2 |
| AT5G21930.1 | P-type ATPase of Arabidopsis 2 |
| AT1G10830.1 | 15-cis-zeta-carotene isomerase |
| AT2G45180.1 | Bifunctional inhibitor/lipid-transfer protein/seed storage 2S albumin superfamily |
| AT2G39480.1 | P-glycoprotein 6 |
| AT5G50160.1 | ferric reduction oxidase 8 |

f wild type and *trm5-1* .

| trm/WT | t test p value |
| --- | --- |
| 3.128146 | 0.02925421 |
| 1.605573 | 0.00240321 |
| 1.487743 | 0.02049437 |
| 1.439604 | 0.00014798 |
| 1.433018 | 0.00550718 |
| 1.425184 | 0.01514819 |
| 1.407337 | 0.00420212 |
| 1.3837 | 0.00878016 |
| 1.378422 | 0.00099998 |
| 1.375129 | 0.04191905 |
| 1.374408 | 0.00124163 |
| 1.367435 | 0.00625017 |
| 1.366468 | 0.01628921 |
| 1.357341 | 0.00118745 |
| 1.354601 | 0.04866243 |
| 1.346094 | 0.0027498 |
| 1.33586 | 0.00015117 |
| 1.32419 | 0.0028351 |
| 1.320625 | 0.0063224 |
| 1.311569 | 0.00085025 |
| 1.307407 | 0.00682643 |
| 1.306586 | 0.00689735 |
| 1.305414 | 0.0493091 |
| 1.303122 | 0.00012724 |
| 1.302678 | 0.00176723 |
| 1.302347 | 0.04382104 |
| 1.297292 | 0.00604892 |
| 1.294226 | 0.04340927 |
| 1.291283 | 0.00126227 |
| 1.291041 | 0.00400034 |
| 1.290292 | 0.02891399 |
| 1.289236 | 0.00653661 |
| 1.285623 | 0.01454794 |
| 1.284872 | 0.00143874 |
| 1.280874 | 0.01898141 |
| 1.277439 | 0.00551195 |
| 1.276558 | 0.02841547 |
| 1.26934 | 0.00501884 |
| 1.265969 | 0.00153152 |
| 1.265867 | 0.04631422 |
| 1.262156 | 0.00126554 |
| 1.261536 | 0.00048284 |

| | |
|---|---|
| 1.260307 | 0.00013914 |
| 1.260091 | 0.00028995 |
| 1.260027 | 0.00245915 |
| 1.255393 | 0.00378779 |
| 1.252927 | 0.01239905 |
| 1.252812 | 0.00477631 |
| 1.250601 | 0.01400981 |
| 1.248513 | 0.01039376 |
| 1.247285 | 0.04339738 |
| 1.245202 | 0.04696226 |
| 1.244363 | 0.00442949 |
| 1.243568 | 0.00025482 |
| 1.241947 | 0.00655161 |
| 1.241103 | 0.03359592 |
| 1.238913 | 0.00161893 |
| 1.238362 | 0.00023115 |
| 1.237631 | 0.00073219 |
| 1.235158 | 0.00546651 |
| 1.234693 | 0.02141542 |
| 1.234479 | 0.00822626 |
| 1.233449 | 0.00020361 |
| 1.233315 | 0.0024433 |
| 1.232086 | 3.3052E-05 |
| 1.231663 | 0.00514625 |
| 1.23157 | 0.01146516 |
| 1.231336 | 0.00093221 |
| 1.231083 | 0.01335797 |
| 1.230543 | 0.00066484 |
| 1.229705 | 0.00077189 |
| 1.227956 | 0.00807558 |
| 1.227417 | 0.00575733 |
| 1.226093 | 0.01512936 |
| 1.220995 | 0.0017665 |
| 1.220918 | 0.04405277 |
| 1.219765 | 0.00392091 |
| 1.218487 | 0.00844595 |
| 1.217771 | 0.00657841 |
| 1.216026 | 0.0196669 |
| 1.215901 | 0.0013631 |
| 1.212652 | 0.00056352 |
| 1.211752 | 0.01704347 |
| 1.211489 | 0.00205014 |
| 1.209537 | 0.00956453 |
| 1.208959 | 0.00522263 |
| 1.208036 | 0.04924578 |
| 1.207788 | 0.01419044 |

| | |
|---|---|
| 1.207577 | 0.00014492 |
| 1.20698 | 0.02520614 |
| 1.206597 | 0.0026503 |
| 1.205993 | 0.00538829 |
| 1.205948 | 0.0257066 |
| 1.205896 | 0.00405732 |
| 1.205189 | 0.00279504 |
| 1.204725 | 0.00514546 |
| 1.202571 | 0.01880134 |
| 1.20153 | 0.00318711 |
| 1.200937 | 0.00081727 |
| 1.20071 | 0.03370771 |
| 1.200528 | 0.00264501 |
| 1.20021 | 0.00059104 |

| trm/WT | t test p value |
|---|---|
| 0.833245 | 0.00147407 |
| 0.831921 | 0.00078813 |
| 0.831595 | 0.00670951 |
| 0.830699 | 0.00805514 |
| 0.830208 | 0.03080168 |
| 0.830024 | 0.00307216 |
| 0.829714 | 0.02285203 |
| 0.829707 | 0.00038822 |
| 0.828955 | 0.00410523 |
| 0.827813 | 6.8645E-05 |
| 0.827677 | 0.00020799 |
| 0.827663 | 0.00227293 |
| 0.82754 | 0.00561431 |
| 0.82723 | 0.00233149 |
| 0.826334 | 0.00139418 |
| 0.825857 | 0.00795867 |
| 0.825747 | 0.00041789 |
| 0.825607 | 0.00607077 |
| 0.825451 | 0.01081488 |
| 0.825105 | 0.01816426 |
| 0.824536 | 0.00170156 |
| 0.824384 | 0.03075649 |
| 0.823871 | 0.00025707 |
| 0.82338 | 0.01502523 |
| 0.822811 | 0.0149186 |
| 0.822557 | 0.02744129 |
| 0.822411 | 0.00070296 |
| 0.821771 | 0.00458194 |
| 0.821617 | 0.00496443 |

| | |
|---|---|
| 0.821316 | 0.00031238 |
| 0.821297 | 0.00745838 |
| 0.82122 | 0.00315175 |
| 0.82062 | 0.00093388 |
| 0.819802 | 0.02310749 |
| 0.818947 | 0.00809534 |
| 0.817891 | 0.02437538 |
| 0.81737 | 3.2217E-05 |
| 0.816582 | 0.00091969 |
| 0.816441 | 0.044379 |
| 0.814498 | 0.00044604 |
| 0.814323 | 6.2556E-05 |
| 0.81416 | 0.00637381 |
| 0.813993 | 0.01263968 |
| 0.81205 | 0.00011732 |
| 0.811796 | 0.00044343 |
| 0.811507 | 0.00358071 |
| 0.811441 | 0.0019323 |
| 0.810928 | 0.0058056 |
| 0.810367 | 0.00170316 |
| 0.810142 | 4.8671E-05 |
| 0.809802 | 0.00842367 |
| 0.809476 | 0.04651566 |
| 0.809459 | 0.03570374 |
| 0.809423 | 0.00610107 |
| 0.809194 | 0.00211864 |
| 0.808924 | 0.04100295 |
| 0.808859 | 0.00091136 |
| 0.807811 | 0.00026816 |
| 0.807716 | 0.0032325 |
| 0.80761 | 0.00279878 |
| 0.80756 | 0.04316129 |
| 0.806968 | 0.0253169 |
| 0.806264 | 0.00796265 |
| 0.806116 | 0.00092419 |
| 0.804394 | 0.0037746 |
| 0.803919 | 0.04648451 |
| 0.802841 | 0.00132951 |
| 0.801535 | 0.00107098 |
| 0.800542 | 0.00193793 |
| 0.799618 | 0.01646216 |
| 0.799609 | 0.00234614 |
| 0.799408 | 0.00396806 |
| 0.79879 | 0.04393597 |
| 0.797977 | 0.02430016 |
| 0.797677 | 0.04438347 |

| | |
|---|---|
| 0.797114 | 0.00380108 |
| 0.795974 | 0.00083246 |
| 0.794769 | 0.00216607 |
| 0.79457 | 0.00036683 |
| 0.794359 | 0.00134238 |
| 0.793529 | 0.00313294 |
| 0.793474 | 0.00282537 |
| 0.792318 | 0.00183858 |
| 0.792182 | 0.00384693 |
| 0.791903 | 0.01417048 |
| 0.791332 | 0.00622567 |
| 0.790491 | 0.00444516 |
| 0.78923 | 0.00502752 |
| 0.789204 | 0.00101108 |
| 0.787687 | 0.03491329 |
| 0.78625 | 0.00090496 |
| 0.785739 | 0.00043276 |
| 0.785542 | 0.00123438 |
| 0.784267 | 0.00367039 |
| 0.783705 | 0.00413216 |
| 0.783472 | 0.01104991 |
| 0.781785 | 0.01616049 |
| 0.779988 | 2.8501E-05 |
| 0.779129 | 0.00123973 |
| 0.77902 | 9.0168E-05 |
| 0.777658 | 0.00096054 |
| 0.777511 | 0.00147436 |
| 0.776176 | 0.00856818 |
| 0.775918 | 0.0032929 |
| 0.77499 | 0.0021116 |
| 0.772077 | 0.03374962 |
| 0.767323 | 0.00191666 |
| 0.767261 | 0.00415623 |
| 0.765066 | 0.0032091 |
| 0.761806 | 0.00043209 |
| 0.757293 | 0.0490751 |
| 0.757224 | 0.00045822 |
| 0.757205 | 0.00057441 |
| 0.757166 | 0.0166265 |
| 0.75712 | 0.00058207 |
| 0.756668 | 0.00070513 |
| 0.75568 | 0.00132648 |
| 0.755265 | 0.00495591 |
| 0.755242 | 0.00028388 |
| 0.751754 | 0.03748085 |
| 0.749991 | 5.076E-05 |

| | |
|---|---|
| 0.748188 | 0.00535278 |
| 0.746688 | 4.05E-05 |
| 0.74206 | 0.00011513 |
| 0.738392 | 0.00157664 |
| 0.737394 | 0.00036367 |
| 0.73648 | 0.00239148 |
| 0.73519 | 0.03297299 |
| 0.734294 | 0.00106006 |
| 0.73298 | 0.0024009 |
| 0.73249 | 0.01344246 |
| 0.728666 | 0.00090183 |
| 0.728066 | 0.00092709 |
| 0.725549 | 0.02024324 |
| 0.724826 | 0.01011777 |
| 0.724627 | 0.0012696 |
| 0.724245 | 0.01473885 |
| 0.722921 | 0.04926844 |
| 0.71713 | 0.00043897 |
| 0.714391 | 0.04212258 |
| 0.713625 | 0.00273744 |
| 0.699223 | 0.00128459 |
| 0.698368 | 0.01986979 |
| 0.696331 | 0.02372759 |
| 0.695775 | 0.00774736 |
| 0.681851 | 0.00596063 |
| 0.681159 | 0.01215182 |
| 0.677991 | 0.02271375 |
| 0.677106 | 4.0231E-05 |
| 0.670863 | 0.00363038 |
| 0.668527 | 0.01040704 |
| 0.667472 | 0.04006534 |
| 0.663206 | 0.01038626 |
| 0.642924 | 0.04949485 |
| 0.627352 | 0.03662174 |
| 0.626816 | 0.00041034 |
| 0.619952 | 0.00502183 |
| 0.584101 | 0.04838368 |
| 0.561363 | 0.00182215 |
| 0.437341 | 0.0134861 |
| 0.404351 | 0.02641821 |

**Supplementary Table S3.** Differentially expressed genes identified by RNA-seq analysis of \

| Accession | Description | log2FoldChange | pvalue | padj |
|---|---|---|---|---|
| AT3G26210 | putative cytochrome P450 T | 2.193326792 | 1.77E-55 | 1.94E-51 |
| AT3G13080 | encodes an ATP-dependent | 3.118787071 | 7.52E-45 | 5.50E-41 |
| AT5G65080 | Is upregulated during verna | 3.385542438 | 1.07E-37 | 5.87E-34 |
| AT5G38900 | Thioredoxin superfamily prc | 4.713704447 | 1.24E-36 | 5.42E-33 |
| AT1G51890 | Leucine-rich repeat protein | 4.384425211 | 2.24E-32 | 6.13E-29 |
| AT3G25010 | receptor like protein 41;(sou | 3.868480298 | 2.32E-31 | 5.66E-28 |
| AT2G37130 | Peroxidase superfamily pro | 2.637540317 | 2.91E-31 | 6.37E-28 |
| AT5G38250 | Protein kinase family protei | 3.505802622 | 2.28E-30 | 4.54E-27 |
| AT5G13080 | WRKY75 is one of several tı | 6.521117457 | 1.27E-27 | 1.99E-24 |
| AT2G14610 | PR1 gene expression is indu | 5.828361305 | 1.41E-27 | 2.06E-24 |
| AT1G26420 | FAD-binding Berberine fam | 3.624306337 | 2.40E-27 | 3.10E-24 |
| AT1G26380 | Functions in the biosynthesi | 3.696614434 | 6.22E-27 | 7.57E-24 |
| AT4G28390 | Encodes a mitochondrial AD | 2.279836486 | 3.10E-25 | 3.40E-22 |
| AT5G42830 | HXXXD-type acyl-transferas | 3.781004753 | 3.43E-24 | 3.42E-21 |
| AT3G01345 | Expressed protein;(source:A | 12.25274396 | 3.93E-24 | 3.74E-21 |
| AT1G33030 | O-methyltransferase family | 4.080041015 | 3.88E-23 | 2.83E-20 |
| AT3G22600 | Bifunctional inhibitor/lipid- | 4.724903488 | 2.41E-21 | 1.51E-18 |
| AT3G15536 | Unknown gene The mRNA i | 5.489505231 | 2.99E-21 | 1.82E-18 |
| AT5G52740 | Copper transport protein fa | 4.360376154 | 4.15E-21 | 2.39E-18 |
| AT5G35935 | transposable_element_gen | 5.176996956 | 3.93E-20 | 2.05E-17 |
| AT1G67520 | lectin protein kinase family | 3.152369868 | 5.82E-20 | 2.90E-17 |
| AT5G44990 | Glutathione S-transferase f | 4.204176876 | 6.22E-20 | 3.03E-17 |
| AT5G46295 | transmembrane protein;(so | 6.679660961 | 6.58E-20 | 3.14E-17 |
| AT2G43570 | chitinase;(source:Araport11 | 5.999316616 | 7.03E-20 | 3.28E-17 |
| AT1G72240 | hypothetical protein;(source | 2.152258719 | 1.18E-19 | 5.40E-17 |
| AT4G13890 | Pyridoxal phosphate (PLP)-c | 5.303231277 | 2.12E-18 | 8.79E-16 |
| AT1G47890 | receptor like protein 7;(sour | 3.231915707 | 2.48E-18 | 1.01E-15 |
| AT3G28510 | P-loop containing nucleosid | 5.446571248 | 3.99E-18 | 1.59E-15 |
| AT1G33960 | Identified as a gene that is | 3.434211736 | 4.33E-18 | 1.70E-15 |
| AT2G04050 | MATE efflux family protein; | 5.211134936 | 4.57E-18 | 1.76E-15 |
| AT5G09290 | Inositol monophosphatase f | 4.040978897 | 8.06E-18 | 3.05E-15 |
| AT5G22570 | member of WRKY Transcrip | 4.681134118 | 1.19E-17 | 4.29E-15 |
| AT2G36780 | UDP-Glycosyltransferase su | 2.624109499 | 1.28E-17 | 4.52E-15 |
| AT2G30750 | putative cytochrome P450 | 3.516574547 | 1.46E-17 | 5.09E-15 |
| AT1G23840 | transmembrane protein;(so | 2.292727408 | 1.86E-17 | 6.18E-15 |
| AT4G12490 | Encodes a member of the A | 2.985203168 | 2.39E-17 | 7.69E-15 |
| AT2G40740 | member of WRKY Transcrip | 5.335637338 | 2.85E-17 | 8.94E-15 |
| AT4G21850 | methionine sulfoxide reduc | 3.203803114 | 3.00E-17 | 9.26E-15 |
| AT1G19250 | FMO1 is required for full ex | 7.13728771 | 3.86E-17 | 1.14E-14 |
| AT2G25440 | receptor like protein 20;(sou | 4.660620576 | 5.86E-17 | 1.67E-14 |
| AT5G53110 | RING/U-box superfamily pr | 2.939782688 | 1.12E-16 | 3.11E-14 |
| AT5G46050 | Encodes a di- and tri-peptid | 3.349394023 | 1.26E-16 | 3.45E-14 |
| AT5G24200 | alpha/beta-Hydrolases supe | 4.352145762 | 2.38E-16 | 6.14E-14 |

| | | | | |
|---|---|---|---|---|
| AT1G52040 | Encodes myrosinase-bindin | 3.137788812 | 2.69E-16 | 6.85E-14 |
| AT3G53210 | nodulin MtN21-like transpo | 2.170998042 | 3.48E-16 | 8.78E-14 |
| AT2G18660 | Encodes PNP-A (Plant Natri | 4.029486782 | 5.24E-16 | 1.31E-13 |
| AT3G26440 | transmembrane protein, pu | 3.384889282 | 5.36E-16 | 1.32E-13 |
| AT1G51860 | Leucine-rich repeat protein | 4.634586086 | 1.37E-15 | 3.28E-13 |
| AT3G07520 | member of Putative ligand- | 2.704083086 | 2.78E-15 | 6.29E-13 |
| AT1G66960 | Terpenoid cyclases family p | 4.78937787 | 3.02E-15 | 6.76E-13 |
| AT1G67980 | Encodes S-adenosyl-L-meth | 4.552109014 | 4.85E-15 | 1.08E-12 |
| AT3G13950 | ankyrin;(source:Araport11) | 3.968471056 | 5.82E-15 | 1.28E-12 |
| AT3G23120 | receptor like protein 38;(so | 2.015888327 | 8.47E-15 | 1.77E-12 |
| AT1G30220 | Inositol transporter present | 4.576842878 | 8.94E-15 | 1.85E-12 |
| AT1G65690 | Encodes NHL6 (NDR1/HIN1- | 3.070845945 | 1.05E-14 | 2.13E-12 |
| AT4G10500 | 2-oxoglutarate (2OG) and F | 6.148208666 | 1.18E-14 | 2.36E-12 |
| AT3G13610 | Encodes a Fe(II)- and 2-oxo | 3.904221306 | 1.36E-14 | 2.69E-12 |
| AT4G03320 | Encodes a component of th | 2.868340494 | 3.09E-14 | 6.00E-12 |
| AT1G72960 | Root hair defective 3 GTP-b | 2.407039121 | 3.82E-14 | 7.35E-12 |
| AT1G13520 | hypothetical protein (DUF1 | 7.656233047 | 7.25E-14 | 1.34E-11 |
| AT4G08093 | pseudogene of dihydroorota | 7.680716257 | 9.25E-14 | 1.66E-11 |
| AT1G68765 | Encodes a small protein of | 4.003357446 | 1.03E-13 | 1.82E-11 |
| AT1G15520 | ABC transporter family invo | 3.520260065 | 1.48E-13 | 2.51E-11 |
| AT3G25510 | disease resistance protein ( | 2.003822315 | 1.81E-13 | 2.98E-11 |
| AT3G24900 | receptor like protein 39;(so | 3.268642425 | 1.84E-13 | 3.01E-11 |
| AT1G26390 | FAD-binding Berberine fam | 6.12860148 | 4.08E-13 | 6.33E-11 |
| AT1G30900 | VACUOLAR SORTING RECEI | 3.075129677 | 4.10E-13 | 6.33E-11 |
| AT2G30770 | putative cytochrome P450 | 3.164015079 | 4.14E-13 | 6.34E-11 |
| AT4G11070 | member of WRKY Transcrip | 4.384550472 | 6.98E-13 | 1.05E-10 |
| AT4G17660 | Protein kinase superfamily | 3.897487229 | 7.84E-13 | 1.16E-10 |
| AT3G53150 | UDP-glucosyl transferase 7 | 4.238490014 | 8.02E-13 | 1.18E-10 |
| AT4G15417 | RNAse II-like 1;(source:Ara | 3.47826412 | 8.35E-13 | 1.22E-10 |
| AT1G02790 | encodes a exopolygalacturo | 4.863887492 | 8.77E-13 | 1.27E-10 |
| AT1G75040 | Thaumatin-like protein invo | 3.943830102 | 1.09E-12 | 1.56E-10 |
| AT1G52000 | Mannose-binding lectin sup | 3.32573686 | 1.36E-12 | 1.90E-10 |
| AT1G04600 | member of Myosin-like pro | 2.97716094 | 1.50E-12 | 2.07E-10 |
| AT2G29350 | senescence-associated gen | 5.197128123 | 1.56E-12 | 2.14E-10 |
| AT2G45220 | Plant invertase/pectin met | 6.383830698 | 1.70E-12 | 2.31E-10 |
| AT1G09080 | Heat shock protein 70 (Hsp | 2.783763091 | 2.12E-12 | 2.86E-10 |
| AT5G24540 | beta glucosidase 31;(source | 5.238693344 | 2.43E-12 | 3.21E-10 |
| AT4G18360 | Encodes a glycolate oxidase | 3.937284614 | 2.58E-12 | 3.37E-10 |
| AT3G60170 | transposable_element_gen | 8.457393682 | 3.66E-12 | 4.70E-10 |
| AT3G47480 | Calcium-binding EF-hand fa | 2.114051426 | 4.57E-12 | 5.76E-10 |
| AT4G04500 | Encodes a cysteine-rich rec | 5.127667031 | 4.76E-12 | 5.96E-10 |
| AT5G18350 | Disease resistance protein ( | 8.684170088 | 5.40E-12 | 6.69E-10 |
| AT5G64000 | 3'(2'),5'-bisphosphate nucle | 3.672328373 | 5.48E-12 | 6.75E-10 |
| AT1G74710 | Encodes a protein with isoc | 2.994792319 | 7.48E-12 | 9.06E-10 |
| AT2G33080 | receptor like protein 28;(so | 5.002567618 | 7.48E-12 | 9.06E-10 |
| AT4G25110 | Encodes a type I metacaspa | 2.027267541 | 8.07E-12 | 9.67E-10 |

| | | | | |
|---|---|---|---|---|
| AT4G15610 | Uncharacterized protein far | 3.420360324 | 8.79E-12 | 1.05E-09 |
| AT3G28580 | P-loop containing nucleosid | 4.05784862 | 1.00E-11 | 1.18E-09 |
| AT3G26470 | Powdery mildew resistance | 2.886608743 | 1.10E-11 | 1.28E-09 |
| AT4G23280 | Encodes a cysteine-rich rece | 2.757689421 | 1.12E-11 | 1.29E-09 |
| AT1G32960 | Subtilase family protein;(so | 3.577405998 | 1.31E-11 | 1.51E-09 |
| AT5G66890 | Leucine-rich repeat (LRR) fa | 3.802509582 | 1.72E-11 | 1.91E-09 |
| AT1G52400 | encodes a member of glyco | 3.218426257 | 1.82E-11 | 2.02E-09 |
| AT2G45510 | member of CYP704A | 2.084153946 | 1.84E-11 | 2.03E-09 |
| AT3G48020 | hypothetical protein;(source | 2.474395345 | 2.15E-11 | 2.35E-09 |
| AT4G21840 | methionine sulfoxide reduc | 5.332926133 | 2.92E-11 | 3.06E-09 |
| AT5G02780 | Encodes a member of the la | 6.204419899 | 3.51E-11 | 3.58E-09 |
| AT4G00700 | C2 calcium/lipid-binding pla | 4.298791491 | 3.54E-11 | 3.59E-09 |
| AT5G11210 | member of Putative ligand- | 3.912935746 | 3.55E-11 | 3.59E-09 |
| AT4G22960 | FAM63A-like protein (DUF5 | 2.78642947 | 4.97E-11 | 4.84E-09 |
| AT2G43000 | Encodes a NAC transcriptio | 3.596691871 | 5.70E-11 | 5.51E-09 |
| AT1G67650 | SRP72 RNA-binding domair | 2.419482493 | 7.45E-11 | 7.10E-09 |
| AT3G28540 | P-loop containing nucleosid | 2.151614679 | 1.14E-10 | 1.04E-08 |
| AT2G29120 | member of Putative ligand- | 2.452182463 | 1.24E-10 | 1.12E-08 |
| AT2G25000 | Pathogen-induced transcrip | 2.645248507 | 1.29E-10 | 1.15E-08 |
| AT1G71390 | receptor like protein 11;(sou | 6.110729239 | 1.80E-10 | 1.55E-08 |
| AT1G37150 | Although HCS2 is predicted | 3.387132968 | 2.31E-10 | 1.94E-08 |
| AT5G23160 | transmembrane protein;(so | 3.811587065 | 2.59E-10 | 2.14E-08 |
| AT3G51860 | cation exchanger 3;(source: | 3.283370565 | 3.10E-10 | 2.53E-08 |
| AT4G23150 | Encodes a cysteine-rich rece | 4.268307375 | 3.79E-10 | 3.06E-08 |
| AT4G22950 | MADS-box protein AGL19 | 3.019561079 | 3.98E-10 | 3.20E-08 |
| AT3G45330 | Concanavalin A-like lectin p | 6.859268111 | 4.06E-10 | 3.25E-08 |
| AT5G45090 | phloem protein 2-A7;(sourc | 6.748282603 | 4.53E-10 | 3.59E-08 |
| AT4G14630 | germin-like protein with N- | 5.760540945 | 6.37E-10 | 4.97E-08 |
| AT1G21310 | Encodes extensin 3. | 3.311447221 | 6.47E-10 | 5.03E-08 |
| AT5G60280 | Plasma membrane localize | 2.333722471 | 6.80E-10 | 5.23E-08 |
| AT5G24530 | Encodes a putative 2OG-Fe | 2.642463446 | 7.05E-10 | 5.40E-08 |
| AT5G44430 | Predicted to encode a PR (p | 4.137699507 | 7.09E-10 | 5.41E-08 |
| AT4G04510 | Encodes a cysteine-rich rece | 3.58377637 | 8.38E-10 | 6.32E-08 |
| AT5G65980 | Auxin efflux carrier family p | 4.211507816 | 8.73E-10 | 6.55E-08 |
| AT4G21903 | MATE efflux family protein; | 3.153312211 | 9.04E-10 | 6.67E-08 |
| AT1G56600 | GolS2 is a galactinol syntha | 3.086935469 | 9.82E-10 | 7.20E-08 |
| AT2G26400 | Encodes a protein predicted | 4.08695219 | 1.08E-09 | 7.90E-08 |
| AT5G19880 | Peroxidase superfamily pro | 6.910308559 | 1.27E-09 | 9.00E-08 |
| AT1G55790 | ferredoxin-fold anticodon-b | 7.374697684 | 1.35E-09 | 9.43E-08 |
| AT5G01900 | member of WRKY Transcrip | 4.294163849 | 1.65E-09 | 1.13E-07 |
| AT1G21240 | encodes a wall-associated l | 4.077352308 | 2.23E-09 | 1.46E-07 |
| AT1G32350 | alternative oxidase 1D;(sou | 5.003527025 | 2.61E-09 | 1.69E-07 |
| AT4G21120 | Encodes a member of the c | 2.747882711 | 2.88E-09 | 1.84E-07 |
| AT5G10760 | Eukaryotic aspartyl protease | 2.710702895 | 3.01E-09 | 1.91E-07 |
| AT5G25260 | SPFH/Band 7/PHB domain-c | 3.031691165 | 3.35E-09 | 2.09E-07 |
| AT1G09935 | Phosphoglycerate mutase fa | 6.494424633 | 3.53E-09 | 2.18E-07 |

| | | | | |
|---|---|---|---|---|
| AT1G44130 | Eukaryotic aspartyl proteas | 5.537936041 | 4.26E-09 | 2.58E-07 |
| AT2G30140 | Encodes a putative glycosyl | 2.004855526 | 4.49E-09 | 2.70E-07 |
| AT1G17615 | TN2 is an atypical TIR-NBS | 4.14108581 | 4.56E-09 | 2.73E-07 |
| AT4G18250 | receptor Serine/Threonine l | 2.636799558 | 4.57E-09 | 2.73E-07 |
| AT5G58940 | Arabidopsis thaliana calmo | 2.471055909 | 5.54E-09 | 3.24E-07 |
| AT3G18610 | Encodes ATNUC-L2 (NUCLE | 2.234923925 | 6.20E-09 | 3.57E-07 |
| AT1G68740 | Encodes PHO1;H1, a memb | 2.228321627 | 7.53E-09 | 4.22E-07 |
| AT3G50770 | calmodulin-like 41;(source:/ | 2.607555975 | 1.17E-08 | 6.18E-07 |
| AT3G47090 | Leucine-rich repeat protein | 2.417017891 | 1.19E-08 | 6.27E-07 |
| AT5G40990 | Component of plant resista | 7.269350813 | 1.33E-08 | 6.95E-07 |
| AT5G64810 | member of WRKY Transcrip | 3.901310858 | 1.58E-08 | 8.00E-07 |
| AT5G65600 | Concanavalin A-like lectin p | 2.105251451 | 1.59E-08 | 8.03E-07 |
| AT3G26220 | cytochrome P450 monooxyg | 2.130483214 | 1.59E-08 | 8.04E-07 |
| AT5G52390 | PAR1 protein;(source:Arapo | 3.80664532 | 1.69E-08 | 8.34E-07 |
| AT4G16260 | Encodes a putative beta-1,3 | 3.861528417 | 1.72E-08 | 8.43E-07 |
| AT1G09480 | similar to Eucalyptus gunnii | 2.889465213 | 1.76E-08 | 8.60E-07 |
| AT1G61550 | S-locus lectin protein kinase | 2.113023251 | 1.86E-08 | 8.99E-07 |
| AT1G51920 | transmembrane protein;(so | 3.683867485 | 2.06E-08 | 9.74E-07 |
| AT1G58225 | hypothetical protein;(source | 4.03676321 | 2.10E-08 | 9.91E-07 |
| AT1G07900 | LOB domain-containing pro | 3.399468371 | 2.17E-08 | 1.02E-06 |
| AT4G23310 | Encodes a cysteine-rich rec | 4.200821253 | 2.19E-08 | 1.02E-06 |
| AT5G01550 | Encodes LecRKA4.2, a mem | 2.742834103 | 2.41E-08 | 1.11E-06 |
| AT4G04490 | Encodes a cysteine-rich rec | 3.342720557 | 2.58E-08 | 1.18E-06 |
| AT5G13320 | Encodes an enzyme capable | 3.738041439 | 2.64E-08 | 1.20E-06 |
| AT3G44300 | Encodes an enzyme that ca | 4.186259038 | 2.94E-08 | 1.31E-06 |
| AT1G13830 | Carbohydrate-binding X8 do | 3.260800097 | 3.57E-08 | 1.58E-06 |
| AT2G44480 | beta glucosidase 17;(source | 2.46917699 | 3.82E-08 | 1.67E-06 |
| AT4G20000 | VQ motif-containing protei | 2.989624915 | 5.21E-08 | 2.23E-06 |
| AT2G47030 | Plant invertase/pectin meth | 4.500333694 | 5.70E-08 | 2.43E-06 |
| AT1G65481 | transmembrane protein;(so | 4.020334492 | 6.04E-08 | 2.54E-06 |
| AT3G05630 | Encodes a member of the P | 2.827114173 | 7.61E-08 | 3.11E-06 |
| AT4G37520 | Peroxidase superfamily pro | 2.15588701 | 8.23E-08 | 3.32E-06 |
| AT5G47960 | Encodes a small molecular | 2.051630885 | 8.40E-08 | 3.39E-06 |
| AT1G65500 | transmembrane protein;(so | 2.696407927 | 8.42E-08 | 3.39E-06 |
| AT5G25820 | Exostosin family protein;(so | 2.574122148 | 9.27E-08 | 3.66E-06 |
| AT3G57260 | beta 1,3-glucanase | 4.004670471 | 9.56E-08 | 3.72E-06 |
| AT3G11640 | transmembrane protein;(so | 3.133369049 | 9.73E-08 | 3.78E-06 |
| AT1G65510 | transmembrane protein;(so | 3.301395794 | 1.02E-07 | 3.95E-06 |
| AT2G15490 | UDP-glycosyltransferase 73 | 3.76652337 | 1.04E-07 | 3.98E-06 |
| AT3G54000 | TIP41-like protein;(source:A | 2.381961266 | 1.25E-07 | 4.74E-06 |
| AT3G28830 | mucin-like protein, putative | 4.251305793 | 1.27E-07 | 4.80E-06 |
| AT3G49340 | Cysteine proteinases superf | 3.887835351 | 1.28E-07 | 4.82E-06 |
| AT1G73010 | Encodes PPsPase1, a pyroph | 2.779768215 | 1.29E-07 | 4.85E-06 |
| AT2G47040 | Share high homologies with | 4.040476886 | 1.37E-07 | 5.07E-06 |
| AT3G13433 | transmembrane protein;(so | 4.72836311 | 1.45E-07 | 5.33E-06 |
| AT4G23160 | Encodes a cysteine-rich rec | 2.769320521 | 1.46E-07 | 5.35E-06 |

| | | | | |
|---|---|---|---|---|
| AT3G09270 | Encodes glutathione transfe | 2.698084739 | 1.54E-07 | 5.57E-06 |
| AT1G68320 | putative transcription factor | 7.472974568 | 1.54E-07 | 5.58E-06 |
| AT5G64790 | O-Glycosyl hydrolases famil | 5.494978748 | 1.81E-07 | 6.39E-06 |
| AT4G11480 | Encodes a cysteine-rich rece | 5.011141157 | 1.82E-07 | 6.42E-06 |
| AT5G37490 | ARM repeat superfamily pro | 2.8162295 | 1.89E-07 | 6.57E-06 |
| AT4G05590 | Encodes NRGA1, a putative | 2.349827028 | 1.95E-07 | 6.76E-06 |
| AT5G40000 | P-loop containing nucleosid | 4.535585689 | 1.98E-07 | 6.86E-06 |
| AT1G10550 | Encodes a membrane-locali | 3.67695241 | 2.31E-07 | 7.85E-06 |
| AT2G29110 | member of Putative ligand- | 3.868645367 | 2.55E-07 | 8.55E-06 |
| AT5G38240 | Protein kinase family protei | 2.022694195 | 2.58E-07 | 8.61E-06 |
| AT1G66880 | Protein kinase superfamily | 2.07528009 | 2.72E-07 | 8.99E-06 |
| AT1G13480 | hypothetical protein (DUF12 | 6.967201499 | 2.78E-07 | 9.19E-06 |
| AT4G14640 | encodes a divergent memb | 2.548510694 | 2.84E-07 | 9.34E-06 |
| AT5G52720 | Copper transport protein fa | 4.660055445 | 2.86E-07 | 9.36E-06 |
| AT3G48630 | hypothetical protein;(source | 3.422346853 | 2.90E-07 | 9.44E-06 |
| AT1G17745 | encodes a 3-Phosphoglycera | 2.216688275 | 3.06E-07 | 9.88E-06 |
| AT1G65484 | transmembrane protein;(so | 2.174785992 | 3.56E-07 | 1.12E-05 |
| AT5G48410 | member of Putative ligand- | 2.968548173 | 3.71E-07 | 1.17E-05 |
| AT5G39390 | Leucine-rich repeat protein | 4.935880356 | 3.92E-07 | 1.22E-05 |
| AT1G17020 | Encodes a novel member of | 3.446565463 | 4.17E-07 | 1.29E-05 |
| AT2G29100 | member of Putative ligand- | 4.325534824 | 4.25E-07 | 1.31E-05 |
| AT1G80130 | Tetratricopeptide repeat (T | 3.037037093 | 5.00E-07 | 1.50E-05 |
| AT4G11170 | Encodes RMG1 (Resistance | 3.936499711 | 5.08E-07 | 1.52E-05 |
| AT4G30270 | encodes a protein similar to | 2.247747765 | 5.33E-07 | 1.59E-05 |
| AT1G61566 | Member of a diversely expr | 4.979762341 | 5.50E-07 | 1.62E-05 |
| AT4G04540 | Encodes a cysteine-rich rece | 3.620441556 | 6.11E-07 | 1.76E-05 |
| AT1G63245 | Member of a large family o | 3.215253706 | 7.03E-07 | 1.97E-05 |
| AT5G40010 | Encodes a mitochondrial AT | 3.778891248 | 7.06E-07 | 1.97E-05 |
| AT5G59320 | Predicted to encode a PR (p | 3.993099733 | 7.11E-07 | 1.98E-05 |
| AT3G46900 | encodes a member of coppe | 2.811921088 | 7.17E-07 | 2.00E-05 |
| AT5G47850 | CRINKLY4 related 4;(source | 3.091670895 | 7.28E-07 | 2.02E-05 |
| AT5G44420 | Encodes an ethylene- and ja | 2.277835458 | 7.32E-07 | 2.03E-05 |
| AT2G32680 | receptor like protein 23;(sou | 3.497956012 | 7.42E-07 | 2.06E-05 |
| AT1G60830 | RNA-binding (RRM/RBD/RN | 6.849093932 | 8.00E-07 | 2.21E-05 |
| AT1G06137 | transmembrane protein;(so | 5.444929356 | 8.52E-07 | 2.33E-05 |
| AT1G34180 | NAC domain containing pro | 2.08204262 | 8.75E-07 | 2.38E-05 |
| AT4G19370 | chitin synthase, putative (D | 2.236750601 | 9.19E-07 | 2.47E-05 |
| AT5G11920 | Encodes a protein with fruc | 4.463495164 | 9.20E-07 | 2.47E-05 |
| AT5G64905 | elicitor peptide 3 precursor; | 2.978499358 | 9.91E-07 | 2.63E-05 |
| AT5G62480 | Encodes glutathione transfe | 3.475922721 | 1.05E-06 | 2.76E-05 |
| AT3G01240 | splicing regulatory glutamin | 4.634589126 | 1.14E-06 | 2.95E-05 |
| AT5G10380 | Encodes a RING finger dom | 2.635988209 | 1.31E-06 | 3.33E-05 |
| AT5G45380 | urea-proton symporter DEG | 2.514066681 | 1.31E-06 | 3.33E-05 |
| AT2G02340 | phloem protein 2-B8;(sourc | 3.822479798 | 1.37E-06 | 3.46E-05 |
| AT3G13432 | transmembrane protein;(so | 4.368267679 | 1.38E-06 | 3.48E-05 |
| AT5G48400 | member of Putative ligand- | 3.819441268 | 1.44E-06 | 3.62E-05 |

| | | | | |
|---|---|---|---|---|
| AT2G37870 | Bifunctional inhibitor/lipid-t | 3.938248985 | 1.49E-06 | 3.73E-05 |
| AT2G43580 | Chitinase family protein;(so | 5.667897286 | 1.61E-06 | 4.00E-05 |
| AT3G01420 | Encodes an alpha-dioxygena | 7.399094422 | 1.67E-06 | 4.12E-05 |
| AT3G29250 | NAD(P)-binding Rossmann- | 5.119440228 | 1.76E-06 | 4.31E-05 |
| AT3G26830 | Mutations in pad3 are defe | 3.606111507 | 1.92E-06 | 4.62E-05 |
| AT3G60164 | Pseudogene of AT3G60966; | 3.695518467 | 2.00E-06 | 4.80E-05 |
| AT1G69480 | EXS (ERD1/XPR1/SYG1) fan | 2.989176779 | 2.05E-06 | 4.89E-05 |
| AT2G46150 | Late embryogenesis abunda | 2.457084923 | 2.67E-06 | 6.13E-05 |
| AT3G04717 | Based on qRT-PCR data, thi | 3.841564295 | 2.83E-06 | 6.43E-05 |
| AT3G28890 | receptor like protein 43;(sou | 2.104840255 | 2.86E-06 | 6.50E-05 |
| AT1G02310 | Glycosyl hydrolase superfar | 3.223388831 | 2.99E-06 | 6.74E-05 |
| AT3G61390 | RING/U-box superfamily pr | 3.15069374 | 3.07E-06 | 6.90E-05 |
| AT3G07820 | Pectin lyase-like superfamil | 4.327484355 | 3.27E-06 | 7.23E-05 |
| AT4G11910 | Acts antagonistically with S | 2.104820876 | 3.33E-06 | 7.37E-05 |
| AT3G11402 | Cysteine/Histidine-rich C1 d | 2.851980278 | 3.82E-06 | 8.30E-05 |
| AT1G52410 | Contains a novel calcium-bi | 2.410849382 | 4.03E-06 | 8.64E-05 |
| AT3G14060 | hypothetical protein;(source | 2.263087673 | 4.05E-06 | 8.66E-05 |
| AT3G22860 | member of eIF3c - eukaryot | 3.715576879 | 4.05E-06 | 8.66E-05 |
| AT2G44240 | NEP-interacting protein (DU | 9.057066942 | 4.10E-06 | 8.73E-05 |
| AT1G66460 | Protein kinase superfamily p | 2.066996226 | 4.35E-06 | 9.15E-05 |
| AT1G54575 | hypothetical protein;(source | 3.560781443 | 4.35E-06 | 9.15E-05 |
| AT4G14390 | Ankyrin repeat family prote | 2.528372323 | 4.43E-06 | 9.25E-05 |
| AT4G05540 | P-loop containing nucleosid | 3.396617361 | 4.96E-06 | 0.000101647 |
| AT3G28750 | hypothetical protein;(source | 2.215565767 | 5.08E-06 | 0.000103808 |
| AT1G10040 | alpha/beta-Hydrolases supe | 2.063324593 | 5.17E-06 | 0.000105395 |
| AT5G44480 | mutant has Altered lateral r | 2.968646597 | 5.18E-06 | 0.000105401 |
| AT1G53790 | F-box and associated intera | 2.061970222 | 5.36E-06 | 0.000108601 |
| AT2G32830 | Encodes Pht1;5, a member | 3.457098428 | 5.48E-06 | 0.00011068 |
| AT5G44380 | FAD-binding Berberine fam | 3.687779997 | 5.77E-06 | 0.000115686 |
| AT2G35070 | transmembrane protein;(so | 2.402117538 | 5.85E-06 | 0.000116644 |
| AT5G06740 | Concanavalin A-like lectin p | 4.494554642 | 5.94E-06 | 0.000118158 |
| AT2G04460 | transposable_element_gen | 6.147138742 | 6.05E-06 | 0.000120005 |
| AT4G38560 | phospholipase-like protein ( | 2.627758971 | 6.06E-06 | 0.000120173 |
| AT3G01270 | Pectate lyase family proteir | 3.674092537 | 6.22E-06 | 0.000122492 |
| AT3G13600 | calmodulin-binding family p | 2.183700326 | 6.52E-06 | 0.000127887 |
| AT3G21520 | Encodes a protein is directly | 3.433580165 | 6.52E-06 | 0.000127887 |
| AT3G14280 | LL-diaminopimelate aminot | 2.064728085 | 6.62E-06 | 0.000129756 |
| AT3G29727 | transposable_element_gen | 4.819418632 | 6.82E-06 | 0.000133273 |
| AT1G66390 | production of anthocyanin p | 5.414166314 | 6.93E-06 | 0.00013507 |
| AT1G13470 | hypothetical protein (DUF12 | 2.875839498 | 7.04E-06 | 0.000136315 |
| AT2G04100 | MATE efflux family protein; | 2.792307355 | 7.14E-06 | 0.000137871 |
| AT1G33840 | LURP-one-like protein (DUF | 3.470426122 | 7.24E-06 | 0.000139158 |
| AT5G39720 | avirulence induced protein 2 | 7.008635885 | 7.39E-06 | 0.000141336 |
| AT1G29715 | pseudogene of Leucine-rich | 2.691248298 | 8.09E-06 | 0.000152953 |
| AT1G71910 | hypothetical protein;(source | 2.477250419 | 8.37E-06 | 0.000156603 |
| AT3G46690 | UDP-Glycosyltransferase su | 2.865981179 | 8.36E-06 | 0.000156603 |

| | | | | |
|---|---|---|---|---|
| AT4G23140 | Arabidopsis thaliana recept | 3.475953715 | 8.40E-06 | 0.000157011 |
| AT5G07430 | Pectin lyase-like superfamil | 4.025292601 | 8.63E-06 | 0.000159998 |
| AT1G74590 | Encodes glutathione transfe | 2.719333742 | 9.06E-06 | 0.000166594 |
| AT1G60740 | Thioredoxin superfamily prc | 4.430380151 | 9.67E-06 | 0.000175242 |
| AT2G29065 | GRAS family transcription f | 2.130012856 | 9.86E-06 | 0.00017811 |
| AT1G18710 | Member of the R2R3 factor | 2.192542477 | 1.04E-05 | 0.000187104 |
| AT1G66920 | Protein kinase superfamily | 2.158252424 | 1.13E-05 | 0.000200865 |
| AT3G62170 | VANGUARD-like protein;(sc | 4.760413499 | 1.27E-05 | 0.000221617 |
| AT1G22990 | Heavy metal transport/detc | 6.502918629 | 1.29E-05 | 0.000223862 |
| AT3G56970 | Encodes a member of the b | 3.969870376 | 1.29E-05 | 0.000224251 |
| AT2G30760 | hypothetical protein;(source | 4.977868402 | 1.36E-05 | 0.000234835 |
| AT4G15280 | UDP-glucosyl transferase 7: | 3.662119192 | 1.44E-05 | 0.000247054 |
| AT3G09960 | Calcineurin-like metallo-pho | 4.393372523 | 1.57E-05 | 0.00026639 |
| AT5G64890 | elicitor peptide 2 precursor; | 3.434887262 | 1.59E-05 | 0.000268628 |
| AT5G13330 | encodes a member of the E | 2.582775323 | 1.67E-05 | 0.000280655 |
| AT2G26310 | Encodes a plastid stroma lo | 2.049338137 | 1.90E-05 | 0.000312576 |
| AT5G12880 | proline-rich family protein;( | 3.135244088 | 2.08E-05 | 0.000337981 |
| AT2G35980 | Encodes a protein whose se | 3.447278987 | 2.12E-05 | 0.00034377 |
| AT1G30160 | hypothetical protein (DUF2 | 2.301412171 | 2.18E-05 | 0.000351511 |
| AT4G08770 | Encodes a putative apoplas | 5.64984999 | 2.36E-05 | 0.000375254 |
| AT1G19610 | Predicted to encode a PR (p | 2.269163492 | 2.38E-05 | 0.000377981 |
| AT1G23140 | Expression is upregulated ir | 2.389325796 | 2.43E-05 | 0.000382974 |
| AT4G25000 | Predicted to be secreted prc | 3.38663699 | 2.43E-05 | 0.000383592 |
| AT5G24090 | Chitinase A (class III) expres | 5.054364475 | 2.44E-05 | 0.00038384 |
| AT2G31910 | member of Putative Na+/H | 2.678156461 | 2.49E-05 | 0.000391844 |
| AT5G46871 | Encodes a defensin-like (DE | 2.196383693 | 2.49E-05 | 0.000391844 |
| AT4G35010 | putative beta-galactosidase | 3.35292026 | 2.78E-05 | 0.000428633 |
| AT1G12940 | member of High affinity nit | 4.5914947 | 2.79E-05 | 0.000428665 |
| AT3G45860 | Encodes a cysteine-rich rece | 2.685776023 | 2.97E-05 | 0.000450126 |
| AT3G18250 | Putative membrane lipopro | 2.950513997 | 3.06E-05 | 0.000460815 |
| AT3G54800 | Pleckstrin homology (PH) ar | 5.147975832 | 3.10E-05 | 0.00046522 |
| AT5G39520 | hypothetical protein (DUF1 | 3.132810629 | 3.13E-05 | 0.000469056 |
| AT5G50030 | Plant invertase/pectin meth | 3.167645478 | 3.17E-05 | 0.000472741 |
| AT1G13340 | Regulator of Vps4 activity ir | 2.97527194 | 3.24E-05 | 0.000480834 |
| AT5G60285 | pre-tRNA tRNA-Leu (antico | 3.819257304 | 3.23E-05 | 0.000480834 |
| AT2G17740 | VACUOLELESS GAMETOPHY | 2.607440937 | 3.26E-05 | 0.000482007 |
| AT5G46350 | member of WRKY Transcrip | 2.300492312 | 3.38E-05 | 0.000496698 |
| AT1G05680 | Encodes a UDP-glucosyltrar | 3.678492959 | 3.46E-05 | 0.000505984 |
| AT5G57480 | P-loop containing nucleosid | 2.290925609 | 3.57E-05 | 0.000518469 |
| AT5G07410 | Encodes a pectin methylest | 4.823137858 | 3.62E-05 | 0.000524936 |
| AT1G13540 | hypothetical protein (DUF1 | 2.31621001 | 3.70E-05 | 0.000534582 |
| AT2G04070 | Expression in rosette leaves | 3.867802817 | 3.76E-05 | 0.000541585 |
| AT1G58340 | Encodes a plant MATE (mul | 2.049337286 | 3.96E-05 | 0.000563166 |
| AT4G15270 | glucosyltransferase-like prc | 4.06943636 | 3.99E-05 | 0.000566821 |
| AT3G29725 | pseudogene of HXXXD-type | 3.46503467 | 4.08E-05 | 0.000575489 |
| AT5G22560 | transmembrane protein, pu | 5.7813991 | 4.32E-05 | 0.000603778 |

| | | | | |
|---|---|---|---|---|
| AT2G25260 | Hyp O-arabinosyltransferas | 2.254092405 | 4.42E-05 | 0.000617187 |
| AT4G24000 | encodes a protein similar to | 3.790668008 | 4.46E-05 | 0.000621429 |
| AT3G60961 | P-loop containing nucleosid | 3.361795618 | 4.75E-05 | 0.00065388 |
| AT2G23830 | PapD-like superfamily prote | 3.554623322 | 4.82E-05 | 0.000661223 |
| AT2G39530 | Uncharacterized protein far | 4.217754463 | 4.95E-05 | 0.000675198 |
| AT4G33550 | Bifunctional inhibitor/lipid- | 3.583869844 | 5.17E-05 | 0.00069595 |
| AT5G15130 | member of WRKY Transcrip | 4.26520156 | 5.67E-05 | 0.000754246 |
| AT4G09770 | TRAF-like family protein;(sc | 3.183831527 | 5.69E-05 | 0.000756444 |
| AT4G15150 | glycine-rich protein;(source | 3.021525234 | 6.37E-05 | 0.000828855 |
| AT2G21490 | dehydrin LEA;(source:Arapo | 2.527307382 | 6.57E-05 | 0.000849913 |
| AT5G15840 | Encodes a protein showing | 2.847156441 | 6.68E-05 | 0.000859587 |
| AT4G18990 | xyloglucan endotransglucos | 3.257744493 | 6.77E-05 | 0.000868724 |
| AT2G32660 | receptor like protein 22;(sou | 3.8975994 | 7.20E-05 | 0.000914382 |
| AT2G26450 | Plant invertase/pectin meth | 4.002642943 | 7.70E-05 | 0.000966906 |
| AT3G30770 | Eukaryotic aspartyl protease | 6.387832975 | 7.76E-05 | 0.000971729 |
| AT2G23110 | Late embryogenesis abunda | 3.570639834 | 8.77E-05 | 0.001072717 |
| AT1G33950 | Avirulence induced gene (A | 5.449157722 | 9.06E-05 | 0.001102307 |
| AT4G22070 | member of WRKY Transcrip | 3.924233576 | 9.28E-05 | 0.001125056 |
| AT3G60540 | Preprotein translocase Sec, | 2.262963984 | 9.59E-05 | 0.001155391 |
| AT5G28680 | Receptor-like kinase require | 5.669745251 | 9.63E-05 | 0.001158729 |
| AT1G61563 | Member of a diversely expr | 3.800175153 | 0.000101554 | 0.001207179 |
| AT1G65730 | Arabidopsis thaliana metal- | 3.42380253 | 0.00010359 | 0.001226415 |
| AT2G22470 | Encodes arabinogalactan-pr | 2.887132727 | 0.000106754 | 0.001257083 |
| AT3G17280 | F-box and associated intera | 2.015614201 | 0.000111873 | 0.001305439 |
| AT4G02250 | Plant invertase/pectin meth | 5.394757414 | 0.000112479 | 0.001311119 |
| AT1G14080 | Encodes an alpha-(1,2)-fucc | 2.893622278 | 0.000114016 | 0.001326923 |
| AT2G41380 | S-adenosyl-L-methionine-de | 2.212235181 | 0.000117087 | 0.001353317 |
| AT3G45130 | lanosterol synthase 1;(sourc | 4.486078691 | 0.000117586 | 0.001356939 |
| AT4G35380 | Encodes one of the function | 2.370787133 | 0.000119644 | 0.001374186 |
| AT4G18430 | RAB GTPase homolog A1E;( | 2.513793898 | 0.000120953 | 0.001386315 |
| AT2G35970 | Late embryogenesis abunda | 3.189065936 | 0.000122991 | 0.001403799 |
| AT2G23270 | transmembrane protein;(so | 4.006340816 | 0.000125271 | 0.001427594 |
| AT1G55560 | SKU5 similar 14;(source:Ara | 5.233491558 | 0.000128256 | 0.001453298 |
| AT3G25882 | encodes a kinase that physi | 2.072354873 | 0.000128843 | 0.001459203 |
| AT1G26970 | Protein kinase superfamily | 3.550114091 | 0.000130437 | 0.001472688 |
| AT4G37530 | Peroxidase superfamily pro | 2.462336162 | 0.000137055 | 0.001530858 |
| AT5G06510 | nuclear factor Y, subunit A1 | 2.371516073 | 0.00013722 | 0.001531914 |
| AT1G54020 | GDSL-motif esterase/acylt | 3.20476481 | 0.000139784 | 0.001551923 |
| AT2G36540 | Haloacid dehalogenase-like | 2.726026361 | 0.000139871 | 0.001552034 |
| AT5G03545 | Expressed in response to ph | 2.440241466 | 0.000141458 | 0.001566082 |
| AT4G19970 | nucleotide-diphospho-sugar | 2.990051247 | 0.000142255 | 0.001571325 |
| AT5G38350 | Disease resistance protein ( | 2.78058278 | 0.00015085 | 0.001639503 |
| AT5G19470 | nudix hydrolase homolog 24 | 4.163344859 | 0.000157793 | 0.001693475 |
| AT1G66700 | A member of the Arabidops | 3.345749991 | 0.000159017 | 0.001704941 |
| AT1G68620 | alpha/beta-Hydrolases supe | 2.920515186 | 0.000166051 | 0.001763968 |
| AT4G11650 | osmotin-like protein | 5.005878165 | 0.000174309 | 0.001830407 |

| | | | | |
|---|---|---|---|---|
| AT5G39880 | transmembrane protein;(so | 3.760462003 | 0.000186031 | 0.001927654 |
| AT1G24520 | Male fertility gene acting o | 4.141626048 | 0.000189535 | 0.001957486 |
| AT3G54150 | Encodes a DNA methyltrans | 2.124756524 | 0.000195807 | 0.002004929 |
| AT1G29140 | Pollen Ole e 1 allergen and | 5.511023414 | 0.000196232 | 0.002007209 |
| AT1G66570 | sucrose-proton symporter 7 | 5.478879569 | 0.000201041 | 0.002048336 |
| AT3G60120 | beta glucosidase 27;(source | 3.660959962 | 0.000201346 | 0.002049539 |
| AT5G45880 | Pollen Ole e 1 allergen and | 3.854432715 | 0.000204444 | 0.002075298 |
| AT1G79900 | encodes a mitochondrial or | 2.155192306 | 0.000207286 | 0.002094453 |
| AT5G38344 | Toll-Interleukin-Resistance | 4.34372768 | 0.000209803 | 0.002117931 |
| AT1G64360 | SAQR is a clade specific pro | 3.014823416 | 0.000221298 | 0.002212581 |
| AT5G06720 | Encodes a peroxidase with | 3.231062577 | 0.000222207 | 0.002219641 |
| AT2G42360 | RING/U-box superfamily pr | 2.59414125 | 0.000235679 | 0.002325479 |
| AT4G23030 | MATE efflux family protein; | 2.084455275 | 0.000239239 | 0.002351212 |
| AT4G22030 | F-box protein with a domai | 3.175258084 | 0.000242448 | 0.0023753 |
| AT2G19190 | Encodes a receptor-like pro | 2.252632271 | 0.000242595 | 0.002375676 |
| AT5G49200 | WD-40 repeat family protei | 3.947899564 | 0.000256091 | 0.002479051 |
| AT4G27480 | Core-2/I-branching beta-1,6 | 2.116020461 | 0.000265766 | 0.002556293 |
| AT5G04150 | Encodes a member of the b | 2.331711492 | 0.000269122 | 0.002582392 |
| AT1G64110 | Target promoter of the mal | 2.387216978 | 0.000273101 | 0.002615999 |
| AT3G44350 | NAC domain containing pro | 3.095045058 | 0.000289952 | 0.002754543 |
| AT1G35490 | bZIP family transcription fa | 5.016370609 | 0.000296317 | 0.002802872 |
| AT1G67920 | hypothetical protein;(source | 2.408378843 | 0.00030721 | 0.002887222 |
| AT1G70390 | F-box and associated intera | 4.364866398 | 0.000315355 | 0.002949335 |
| AT5G59680 | Leucine-rich repeat protein | 2.436915475 | 0.000316441 | 0.002957494 |
| AT2G28710 | C2H2-type zinc finger family | 2.554056663 | 0.0003202 | 0.002984992 |
| AT4G13900 | pseudogene of receptor like | 2.219205517 | 0.000336678 | 0.003110834 |
| AT2G02010 | glutamate decarboxylase 4; | 2.466155177 | 0.000339617 | 0.003132708 |
| AT5G48140 | Pectin lyase-like superfamil | 5.632206119 | 0.000347012 | 0.003186177 |
| AT5G17330 | Encodes one of two isoform | 2.68557752 | 0.000350604 | 0.003213773 |
| AT3G33528 | hypothetical protein;(source | 3.262358669 | 0.000358386 | 0.003271431 |
| AT1G01980 | member of Reticuline oxida | 5.567017431 | 0.000358545 | 0.003271519 |
| AT5G47000 | Peroxidase superfamily pro | 4.708944886 | 0.000360791 | 0.003286909 |
| AT2G15480 | UDP-glucosyl transferase 73 | 2.055998976 | 0.000365662 | 0.003313022 |
| AT1G13530 | hypothetical protein (DUF12 | 2.729586663 | 0.000367537 | 0.003328635 |
| AT2G38940 | Encodes Pht1;4, a member | 2.56733342 | 0.00037315 | 0.003371111 |
| AT4G11880 | AGL12, AGL14, and AGL17 a | 5.37572704 | 0.000384162 | 0.003453379 |
| AT3G07850 | Pectin lyase-like superfamil | 3.848046272 | 0.000385372 | 0.00346156 |
| AT3G47780 | member of ATH subfamily | 2.033928879 | 0.000397027 | 0.003541582 |
| AT1G28570 | SGNH hydrolase-type estera | 2.097705944 | 0.000404519 | 0.00358943 |
| AT1G73260 | Encodes a trypsin inhibitor i | 5.269453025 | 0.000407674 | 0.003613039 |
| AT1G44318 | Aldolase superfamily protei | 5.385147657 | 0.000417925 | 0.003687493 |
| AT3G30120 | pseudogene of mediator of | 3.313235236 | 0.00041791 | 0.003687493 |
| AT2G44380 | Cysteine/Histidine-rich C1 d | 2.354218838 | 0.000428258 | 0.003754489 |
| AT1G36622 | transmembrane protein;(so | 2.726270786 | 0.000437242 | 0.00381647 |
| AT1G61800 | glucose6-Phosphate/phospl | 2.837759944 | 0.000439116 | 0.00382673 |
| AT4G12735 | Encodes a peroxisomal prot | 3.072945931 | 0.000440815 | 0.003838488 |

| | | | | |
|---|---|---|---|---|
| AT5G51440 | HSP20-like chaperones supe | 2.125238488 | 0.000445153 | 0.003870109 |
| AT4G31970 | member of CYP82C | 6.52359938 | 0.000472559 | 0.004058494 |
| AT5G15110 | Pectate lyase family proteir | 4.479008288 | 0.000474994 | 0.004075561 |
| AT1G33730 | member of CYP76C | 3.225572091 | 0.000483122 | 0.004134634 |
| AT3G28790 | transmembrane protein, pu | 4.831549329 | 0.000491735 | 0.004187097 |
| AT5G22550 | transmembrane protein, pu | 4.649426266 | 0.000494993 | 0.004209935 |
| AT3G13400 | SKU5 similar 13;(source:Ara | 2.134181565 | 0.000499126 | 0.004235231 |
| AT2G17500 | Auxin efflux carrier family p | 2.205909031 | 0.000500253 | 0.00423996 |
| AT3G57100 | transmembrane protein, pu | 2.516132166 | 0.000510771 | 0.004319001 |
| AT4G39610 | MIZU-KUSSEI-like protein (I | 4.372951026 | 0.000514849 | 0.004345101 |
| AT2G05850 | serine carboxypeptidase-like | 5.911229935 | 0.000522004 | 0.004395339 |
| AT5G36970 | NDR1/HIN1-like protein, exp | 5.245788205 | 0.000526447 | 0.004425951 |
| AT3G25490 | Protein kinase family protei | 5.278222409 | 0.000537565 | 0.004505603 |
| AT5G24110 | member of WRKY Transcrip | 2.27382612 | 0.000554093 | 0.004617655 |
| AT1G09930 | oligopeptide transporter | 2.184434131 | 0.000560817 | 0.00466482 |
| AT3G60970 | member of MRP subfamily | 3.314480401 | 0.000563314 | 0.00468382 |
| AT3G48850 | Encodes a mitochondrial ph | 2.929762735 | 0.000586217 | 0.004842466 |
| AT3G44540 | Encodes a member of the e | 5.40537424 | 0.000587217 | 0.004847635 |
| AT1G22830 | Tetratricopeptide repeat (T | 2.068224402 | 0.000595707 | 0.004901112 |
| AT3G50940 | P-loop containing nucleosid | 2.179642092 | 0.000642898 | 0.005205387 |
| AT3G13435 | transmembrane protein;(so | 4.288674442 | 0.000649482 | 0.005247065 |
| AT2G26390 | Serine protease inhibitor (S | 2.720956692 | 0.000652144 | 0.005257322 |
| AT5G12000 | kinase with adenine nucleo | 2.949347522 | 0.000693887 | 0.005508387 |
| AT5G39670 | Calcium-binding EF-hand fa | 2.285698195 | 0.000695506 | 0.005519242 |
| AT1G04470 | hypothetical protein (DUF8: | 4.283027029 | 0.000708481 | 0.005601928 |
| AT3G28980 | mediator of RNA polymeras | 5.618791514 | 0.000760717 | 0.005933646 |
| AT2G37750 | hypothetical protein;(source | 2.394138753 | 0.00077589 | 0.006015619 |
| AT1G26410 | FAD-binding Berberine fam | 3.482544192 | 0.000787028 | 0.006080468 |
| AT5G22530 | hypothetical protein;(source | 2.807771121 | 0.000826712 | 0.006309519 |
| AT2G26850 | F-box family protein;(source | 5.836763038 | 0.0008315 | 0.006339481 |
| AT4G03290 | EF hand calcium-binding pr | 4.286442301 | 0.000900181 | 0.006743205 |
| AT4G23320 | Encodes a cysteine-rich rece | 2.843128493 | 0.000912072 | 0.0068067 |
| AT2G39030 | Encodes a protein that acts | 3.368181233 | 0.000924106 | 0.006870789 |
| AT3G56275 | pseudogene of expressed pr | 2.520266821 | 0.000966832 | 0.007125635 |
| AT1G51850 | Leucine-rich repeat protein | 2.242535826 | 0.000984827 | 0.007241219 |
| AT1G34060 | Pyridoxal phosphate (PLP)-c | 2.127776227 | 0.001010618 | 0.007413452 |
| AT3G28210 | Encodes a putative zinc fing | 2.116204346 | 0.00102487 | 0.007505442 |
| AT4G23700 | member of Putative Na+/H- | 2.531294612 | 0.001081677 | 0.007874118 |
| AT3G43120 | SAUR-like auxin-responsive | 5.145297352 | 0.001086754 | 0.007900586 |
| AT2G33100 | encodes a gene similar to c | 2.33978016 | 0.001130404 | 0.008155723 |
| AT3G29730 | transposable_element_gen | 3.767995661 | 0.001171454 | 0.008393886 |
| AT4G14450 | A member of a small famil | 2.301560872 | 0.001207833 | 0.008601153 |
| AT4G11655 | Uncharacterized protein fa | 3.257227866 | 0.00139079 | 0.009631857 |
| AT3G57950 | cotton fiber protein;(source | 3.450868088 | 0.001408766 | 0.009737893 |
| AT4G35180 | LYS/HIS transporter 7;(sour | 2.665096049 | 0.001467778 | 0.01005703 |
| AT3G60966 | RING/U-box superfamily pr | 2.281131281 | 0.00149915 | 0.010230426 |

| | | | | |
|---|---|---|---|---|
| AT1G35230 | Encodes arabinogalactan-pı | 2.06289086 | 0.001505702 | 0.010263774 |
| AT1G70170 | mutant has Late flowering; | 2.417625496 | 0.001525086 | 0.01035906 |
| AT3G11430 | sn-glycerol-3-phosphate 2-( | 5.066085068 | 0.001525007 | 0.01035906 |
| AT3G43860 | glycosyl hydrolase 9A4;(sou | 4.706094011 | 0.001537646 | 0.010432986 |
| AT5G63225 | glycosyl hydrolase family pr | 4.579002385 | 0.001551483 | 0.010502569 |
| AT3G45660 | Encodes a member of the N | 2.842591944 | 0.001560292 | 0.010555682 |
| AT1G05580 | member of Putative Na+/H | 3.153536428 | 0.001568987 | 0.010600504 |
| AT1G47880 | pseudogene of receptor like | 5.720965344 | 0.001582456 | 0.010643227 |
| AT3G62180 | Plant invertase/pectin meth | 4.820637526 | 0.001601323 | 0.010750338 |
| AT2G13810 | AGD2-like defense respons | 2.704174658 | 0.00161616 | 0.010826738 |
| AT1G47395 | hypothetical protein;(source | 2.919610214 | 0.001634046 | 0.010912548 |
| AT3G14040 | Pectin lyase-like superfamil | 4.341591527 | 0.001640077 | 0.010946831 |
| AT5G18270 | NAC domain containing pro | 2.324090844 | 0.001662476 | 0.011062654 |
| AT3G62710 | Glycosyl hydrolase family pı | 2.74536162 | 0.001702552 | 0.011281402 |
| AT2G04135 | transposable_element_gen | 2.238553871 | 0.001729731 | 0.01142008 |
| AT1G13609 | Encodes a defensin-like (DE | 3.009351605 | 0.00177117 | 0.011651569 |
| AT4G17460 | Encodes a class II HD-ZIP pr | 2.203194536 | 0.00186472 | 0.012117919 |
| AT3G02810 | Encodes a receptor-like cyto | 3.027553281 | 0.001932491 | 0.012480648 |
| AT1G78780 | pathogenesis-related famil | 2.018631814 | 0.001948325 | 0.012557015 |
| AT4G16820 | Encodes a lipase that hydro | 2.108932342 | 0.001990884 | 0.012778725 |
| AT2G18193 | P-loop containing nucleosid | 2.517248002 | 0.002005149 | 0.012851308 |
| AT4G24640 | Encodes AppB protein (App | 5.089450555 | 0.002048855 | 0.013074268 |
| AT3G56980 | Encodes a member of the b | 3.607511371 | 0.002076556 | 0.013208755 |
| AT3G01175 | transmembrane protein;(so | 2.46529338 | 0.002126016 | 0.013425986 |
| AT5G35575 | transposable_element_gen | 2.812098824 | 0.002173919 | 0.013649872 |
| AT1G70260 | nodulin MtN21-like transpo | 2.547491274 | 0.002187251 | 0.013711596 |
| AT1G66600 | A member of WRKY Transcı | 2.962996806 | 0.002235189 | 0.013918986 |
| AT1G76960 | transmembrane protein;(so | 2.3189965 | 0.002452309 | 0.014990404 |
| AT5G61730 | Encodes an ER-localized AB | 2.433607722 | 0.002495675 | 0.015142755 |
| AT3G13390 | SKU5 similar 11;(source:Ara | 2.373504244 | 0.002586608 | 0.01558504 |
| AT2G31865 | poly(ADP-ribose) glycohydro | 2.074269127 | 0.002658242 | 0.01595126 |
| AT3G60470 | transmembrane protein, pu | 2.686204789 | 0.002668298 | 0.015993737 |
| AT3G25165 | Member of a diversely expr | 4.929815015 | 0.0026831 | 0.01605019 |
| AT3G28810 | mediator of RNA polymeras | 2.97273618 | 0.002684304 | 0.01605019 |
| AT1G55230 | proteinase inhibitor I4, serp | 3.649963069 | 0.002691593 | 0.016076238 |
| AT1G76470 | NAD(P)-binding Rossmann- | 2.2610677 | 0.002690883 | 0.016076238 |
| AT4G18980 | Encodes a nuclear-targeted | 3.482693136 | 0.002854121 | 0.016786334 |
| AT5G57010 | calmodulin-binding family p | 2.026112037 | 0.002882288 | 0.016915697 |
| AT3G49540 | hypothetical protein;(source | 2.245984182 | 0.002905625 | 0.017011678 |
| AT1G67000 | Protein kinase superfamily | 2.46697387 | 0.003038714 | 0.017588925 |
| AT1G65610 | Six-hairpin glycosidases sup | 2.886208115 | 0.00306715 | 0.017719348 |
| AT3G56891 | Heavy metal transport/deto | 3.825067748 | 0.003325639 | 0.018828163 |
| AT4G28460 | transmembrane protein;(so | 2.489481459 | 0.003325884 | 0.018828163 |
| AT2G21820 | seed maturation protein;(so | 2.55260242 | 0.003394504 | 0.019128215 |
| AT1G07620 | GTP-binding protein Obg/Cg | 2.19568423 | 0.003434795 | 0.019282273 |
| AT1G66870 | Carbohydrate-binding X8 do | 3.664279114 | 0.003583595 | 0.019967956 |

| | | | | |
|---|---|---|---|---|
| AT3G25170 | Member of a diversely expr | 3.690577678 | 0.003811719 | 0.020930564 |
| AT2G02330 | pseudogene of phloem prot | 3.909886147 | 0.003813089 | 0.020932844 |
| AT5G52730 | Copper transport protein fa | 4.279706471 | 0.00394548 | 0.02150885 |
| AT5G27870 | Plant invertase/pectin meth | 4.537702264 | 0.003973197 | 0.02162232 |
| AT2G47770 | Encodes a membrane-boun | 2.656568189 | 0.004041729 | 0.021906003 |
| AT2G16730 | putative beta-galactosidase | 3.304736039 | 0.004097723 | 0.022156471 |
| AT2G24450 | FASCICLIN-like arabinogala | 2.280507165 | 0.004135519 | 0.022295402 |
| AT3G20580 | COBRA-like protein 10 prec | 3.141716628 | 0.004222375 | 0.022624581 |
| AT1G05000 | Encodes an atypical dual-sp | 2.102136161 | 0.004395695 | 0.02328582 |
| AT5G54165 | Avr9/Cf-9 rapidly elicited pr | 2.009912836 | 0.004460624 | 0.023510493 |
| AT3G29034 | transmembrane protein;(so | 2.103038287 | 0.004469075 | 0.023532404 |
| AT1G61480 | S-locus lectin protein kinase | 2.28338103 | 0.004531971 | 0.02376656 |
| AT5G06730 | Peroxidase superfamily pro | 2.997285811 | 0.004637398 | 0.024206926 |
| AT2G02140 | Predicted to encode a PR (p | 4.258344808 | 0.00468504 | 0.024371102 |
| AT2G03360 | Glycosyltransferase family | 6.279307715 | 0.004787604 | 0.024702914 |
| AT1G47400 | hypothetical protein;(source | 2.419283348 | 0.004792271 | 0.024706167 |
| AT5G22380 | NAC domain containing pro | 2.542720019 | 0.004967857 | 0.025388822 |
| AT3G46770 | AP2/B3-like transcriptional | 3.987666416 | 0.005022584 | 0.025592264 |
| AT3G57690 | Encodes a putative arabino | 2.63590792 | 0.005052736 | 0.02571641 |
| AT5G46960 | Pectin methylesterase inhib | 6.220723961 | 0.005202531 | 0.026337177 |
| AT4G10250 | Columbia endomembrane-l | 2.69313862 | 0.005242403 | 0.02650944 |
| AT5G12960 | proline-tRNA ligase (DUF16 | 3.162354427 | 0.005406505 | 0.027182536 |
| AT3G50460 | Homolog of RPW8 | 3.034299732 | 0.00545211 | 0.027349111 |
| AT4G26200 | Member of a family of prot | 2.323557341 | 0.005612683 | 0.028026345 |
| AT5G20390 | Glycosyl hydrolase superfar | 4.565242968 | 0.00585411 | 0.029046666 |
| AT1G60970 | SNARE-like superfamily pro | 2.2099167 | 0.005899321 | 0.0292313 |
| AT3G48400 | Cysteine/Histidine-rich C1 d | 2.630714546 | 0.005927353 | 0.029323816 |
| AT2G07040 | Pollen receptor kinase. Coe | 3.573539712 | 0.006050488 | 0.02977845 |
| AT1G01680 | plant U-box 54;(source:Ara | 2.99875937 | 0.006125802 | 0.030054672 |
| AT1G03050 | ENTH/ANTH/VHS superfam | 2.482403143 | 0.006142715 | 0.03010097 |
| AT1G69930 | Encodes glutathione transfe | 3.245062725 | 0.006198024 | 0.030320811 |
| AT4G28420 | Tyrosine transaminase fam | 3.280779312 | 0.00628176 | 0.030614332 |
| AT5G55410 | Bifunctional inhibitor/lipid- | 5.061994106 | 0.006309401 | 0.030709508 |
| AT5G67340 | ARM repeat superfamily pr | 2.018047207 | 0.006370516 | 0.03091632 |
| AT1G29290 | B-cell lymphoma 6 protein; | 2.039617494 | 0.006446167 | 0.031221265 |
| AT3G60140 | Encodes a protein similar to | 4.310081207 | 0.006662432 | 0.032007119 |
| AT4G18540 | transmembrane protein;(so | 2.383438245 | 0.006743598 | 0.032276824 |
| AT1G51913 | transmembrane protein;(so | 4.361520571 | 0.006762513 | 0.032346175 |
| AT3G26110 | Anther-specific protein agp | 3.570834364 | 0.006893776 | 0.032791016 |
| AT4G07940 | pre-mRNA-splicing factor C | 4.636677046 | 0.006989638 | 0.03315051 |
| AT3G24580 | F-box and associated intera | 4.106163902 | 0.007151358 | 0.033815113 |
| AT1G12950 | root hair specific 2;(source: | 2.014765386 | 0.007206183 | 0.034030319 |
| AT2G21930 | A paternally expressed imp | 4.025723683 | 0.007330357 | 0.034459842 |
| AT3G01760 | Encodes an amino acid tran | 2.146055672 | 0.00733427 | 0.034464468 |
| AT2G35765 | hypothetical protein;(source | 4.518463938 | 0.007393975 | 0.034685557 |
| AT1G69940 | Encodes a protein with pect | 3.990247976 | 0.007399802 | 0.034698045 |

| | | | | |
|---|---|---|---|---|
| AT2G04450 | Encodes a protein with NAD | 2.061937811 | 0.007418239 | 0.034747343 |
| AT2G26410 | IQ-domain 4;(source:Arapor | 3.121741197 | 0.007468932 | 0.034887898 |
| AT3G10320 | MUCI21 is a GT61 protein r | 3.131936484 | 0.00762243 | 0.03537879 |
| AT3G27710 | RING/U-box superfamily pr | 2.08451019 | 0.007884523 | 0.036260814 |
| AT5G61720 | hypothetical protein (DUF12 | 2.372781197 | 0.007938854 | 0.036484449 |
| AT3G05610 | Plant invertase/pectin meth | 2.939234028 | 0.008010739 | 0.036737808 |
| AT1G76965 | Encodes a Protease inhibito | 2.039811679 | 0.008173224 | 0.037311287 |
| AT3G04220 | Disease resistance protein ( | 2.098012844 | 0.00833884 | 0.037893764 |
| AT3G10815 | RING/U-box superfamily pr | 2.477949282 | 0.008368703 | 0.038013712 |
| AT1G02580 | Encodes a putative transcrip | 4.80793042 | 0.008411951 | 0.038146936 |
| AT2G36440 | hypothetical protein;(source | 2.683798844 | 0.008523763 | 0.038494755 |
| AT1G48400 | F-box/RNI-like/FBD-like dor | 2.267229871 | 0.009049631 | 0.040236141 |
| AT1G66210 | Subtilisin-like serine endope | 3.011863888 | 0.009148369 | 0.040597005 |
| AT1G15670 | Encodes a member of a fan | 2.035477743 | 0.009178265 | 0.040704415 |
| AT3G19690 | CAP (Cysteine-rich secretory | 2.257058153 | 0.009209629 | 0.040810494 |
| AT2G36020 | HVA22-like protein J;(source | 2.725824161 | 0.0093123 | 0.041198849 |
| AT5G49310 | Putative importin alpha isof | 3.550415029 | 0.009416388 | 0.041533638 |
| AT4G37990 | Encodes an aromatic alcoho | 2.449275961 | 0.00956311 | 0.042016289 |
| AT4G14368 | Regulator of chromosome c | 2.658389014 | 0.009775245 | 0.042721154 |
| AT1G52810 | 2-oxoglutarate (2OG) and F | 2.044396747 | 0.009796475 | 0.04278835 |
| AT2G35742 | snoRNA;(source:Araport11) | 2.154158203 | 0.010466852 | 0.044972798 |
| AT4G11500 | pseudogene of cysteine-rich | 3.002621146 | 0.010487918 | 0.045054485 |
| AT1G36640 | transmembrane protein;(so | 4.551177345 | 0.010557607 | 0.045305331 |
| AT1G08140 | member of Putative Na+/H+ | 4.915159897 | 0.010580598 | 0.045386231 |
| AT5G18910 | Protein kinase superfamily p | 4.810510888 | 0.010709984 | 0.045798882 |
| AT3G01700 | Encodes an arabinogalactan | 3.022996867 | 0.011180496 | 0.047453263 |
| AT3G59620 | Mannose-binding lectin sup | 3.677978943 | 0.011198269 | 0.047488405 |
| AT2G47200 | hypothetical protein;(source | 3.839617318 | 0.011698365 | 0.049147031 |
| AT2G38380 | Peroxidase superfamily prot | 4.923875921 | 0.011716937 | 0.049215626 |
| AT5G20670 | DUF1677 family protein (DU | 2.46861214 | 0.011721326 | 0.049224631 |
| AT3G44510 | alpha/beta-Hydrolases supe | 3.338478488 | 0.011926384 | 0.049919329 |

| **Accession** | **DescriptionDescription** | **log2FoldChange** | **pvalue** | **padj** |
|---|---|---|---|---|
| AT5G52310 | cold regulated gene, the 5' | -2.280968657 | 4.00E-83 | 8.77E-79 |
| AT1G33811 | GDSL-motif esterase/acylt | -2.502119903 | 1.96E-32 | 6.13E-29 |
| AT5G20630 | Encodes a germin-like prote | -4.258970021 | 2.40E-29 | 4.39E-26 |
| AT3G06880 | Transducin/WD40 repeat-li | -2.222235951 | 8.30E-29 | 1.40E-25 |
| AT2G43550 | Encodes a defensin-like (DE | -3.102308263 | 4.32E-24 | 3.95E-21 |
| AT5G63140 | purple acid phosphatase 29 | -2.172265013 | 1.19E-23 | 1.04E-20 |
| AT5G39320 | UDP-glucose 6-dehydrogena | -2.163754332 | 1.28E-23 | 1.08E-20 |
| AT2G32990 | glycosyl hydrolase 9B8;(sou | -2.964979449 | 2.23E-23 | 1.68E-20 |
| AT1G58370 | Encodes a protein with xyla | -2.773996681 | 8.78E-23 | 6.21E-20 |
| AT2G05660 | transposable_element_gen | -11.43709743 | 1.41E-21 | 9.37E-19 |
| AT1G66100 | Predicted to encode a PR (p | -3.891943476 | 1.99E-21 | 1.28E-18 |
| AT1G04800 | glycine-rich protein;(source | -4.242570902 | 3.32E-21 | 1.97E-18 |
| AT1G67750 | Pectate lyase family protein | -3.777733376 | 8.11E-21 | 4.56E-18 |

| | | | | |
|---|---|---|---|---|
| AT3G05800 | AtBS1(activation-tagged BF | -2.665471509 | 4.42E-20 | 2.25E-17 |
| AT1G04680 | Pectin lyase-like superfamil | -2.545635426 | 8.74E-19 | 3.76E-16 |
| AT1G31690 | Copper amine oxidase fami | -6.199604353 | 2.12E-18 | 8.79E-16 |
| AT4G38770 | Encodes one of four proline | -3.295708756 | 8.32E-18 | 3.09E-15 |
| AT1G55330 | Encodes a putative arabinog | -2.343184114 | 1.67E-17 | 5.74E-15 |
| AT1G19940 | glycosyl hydrolase 9B5;(sou | -3.198440073 | 4.39E-17 | 1.28E-14 |
| AT2G30010 | Encodes a member of the T | -2.129416294 | 5.58E-17 | 1.61E-14 |
| AT5G28290 | Encodes AtNek3, a membe | -2.054762529 | 1.38E-16 | 3.68E-14 |
| AT4G29610 | Cytidine/deoxycytidylate de | -2.147215992 | 1.18E-15 | 2.84E-13 |
| AT4G25780 | CAP (Cysteine-rich secretory | -4.177114441 | 2.29E-15 | 5.34E-13 |
| AT5G23820 | ML3 can be modified by NE | -3.707746269 | 2.45E-15 | 5.66E-13 |
| AT5G40390 | Encodes a protein which mi | -2.214467065 | 2.55E-15 | 5.82E-13 |
| AT5G36910 | Encodes a thionin that is ex | -5.349764918 | 6.82E-15 | 1.48E-12 |
| AT1G31710 | Copper amine oxidase fami | -4.996672946 | 8.42E-15 | 1.77E-12 |
| AT5G15230 | Encodes gibberellin-regulat | -3.561126039 | 8.47E-15 | 1.77E-12 |
| AT2G30420 | In a tandem repeat with AT | -4.556834485 | 1.28E-14 | 2.55E-12 |
| AT4G36360 | putative beta-galactosidase | -2.594874883 | 1.44E-14 | 2.81E-12 |
| AT3G53650 | Histone superfamily proteir | -3.747187837 | 6.31E-14 | 1.18E-11 |
| AT1G65450 | Contains dual transcription | -2.217526134 | 1.08E-13 | 1.89E-11 |
| AT5G01015 | transmembrane protein;(so | -4.654523824 | 1.28E-13 | 2.22E-11 |
| AT5G55730 | Encodes fasciclin-like arabi | -3.244923931 | 1.75E-13 | 2.93E-11 |
| AT5G07030 | Eukaryotic aspartyl proteas | -4.128302642 | 1.78E-13 | 2.96E-11 |
| AT5G67070 | Member of a diversely expr | -2.496288049 | 2.20E-13 | 3.57E-11 |
| AT2G34620 | Mitochondrial transcription | -3.146609474 | 3.14E-13 | 4.99E-11 |
| AT1G41830 | SKU5-similar 6;(source:Ara | -2.448895925 | 3.18E-13 | 5.02E-11 |
| AT1G53700 | The WAG1 and its homolog | -4.170392685 | 3.35E-13 | 5.24E-11 |
| AT2G10940 | Bifunctional inhibitor/lipid- | -4.654248189 | 1.04E-12 | 1.49E-10 |
| AT3G46490 | 2-oxoglutarate (2OG) and F | -4.422916365 | 1.17E-12 | 1.66E-10 |
| AT1G29920 | Encodes lhcb1.1 a compone | -8.08686096 | 1.22E-12 | 1.72E-10 |
| AT4G12730 | AF333971 Arabidopsis thali | -2.741852377 | 1.26E-12 | 1.77E-10 |
| AT1G60590 | Pectin lyase-like superfamil | -3.488170028 | 1.37E-12 | 1.91E-10 |
| AT3G23810 | S-adenosyl-l-homocysteine | -2.917623208 | 2.23E-12 | 2.97E-10 |
| AT5G18460 | carboxyl-terminal peptidase | -2.133827511 | 2.23E-12 | 2.97E-10 |
| AT1G55260 | Bifunctional inhibitor/lipid- | -2.240949517 | 2.94E-12 | 3.80E-10 |
| AT3G53730 | Histone superfamily proteir | -2.389082383 | 4.34E-12 | 5.54E-10 |
| AT3G06130 | Heavy metal transport/detc | -2.10952717 | 5.06E-12 | 6.30E-10 |
| AT5G45670 | GDSL-motif esterase/acylt | -6.233395609 | 7.41E-12 | 9.06E-10 |
| AT3G49260 | IQ-domain 21;(source:Arap | -3.070378084 | 7.58E-12 | 9.13E-10 |
| AT2G36870 | Encodes a xyloglucan endot | -3.75367272 | 9.00E-12 | 1.07E-09 |
| AT1G53520 | Encodes a plastid stroma lo | -3.429593254 | 1.05E-11 | 1.23E-09 |
| AT1G12090 | extensin-like protein (ELP) | -3.636365684 | 1.58E-11 | 1.78E-09 |
| AT2G02780 | Leucine-rich repeat protein | -2.258702651 | 1.60E-11 | 1.78E-09 |
| AT2G32690 | Glycine-rich protein similar | -2.01468103 | 1.87E-11 | 2.05E-09 |
| AT5G08000 | Encodes a member of the X | -6.451210676 | 2.26E-11 | 2.45E-09 |
| AT1G01900 | Encodes AtSBT1.1, a subtili | -2.02631756 | 2.29E-11 | 2.48E-09 |
| AT4G20940 | Encodes a plasma-membra | -3.547500503 | 2.51E-11 | 2.70E-09 |

| | | | | |
|---|---|---|---|---|
| AT1G14700 | purple acid phosphatase 3;( | -2.287124183 | 2.85E-11 | 3.03E-09 |
| AT3G04290 | Li-tolerant lipase 1;(source: | -6.060398309 | 2.98E-11 | 3.11E-09 |
| AT3G16670 | Pollen Ole e 1 allergen and | -4.20418224 | 3.29E-11 | 3.40E-09 |
| AT1G64390 | glycosyl hydrolase 9C2;(sou | -4.127759623 | 3.44E-11 | 3.53E-09 |
| AT2G20635 | protein kinase and Mad3-BU | -2.430186743 | 3.73E-11 | 3.75E-09 |
| AT2G35190 | plant-specific SNARE locate | -2.583108366 | 4.62E-11 | 4.54E-09 |
| AT4G34580 | Encodes COW1 (can of wor | -3.190039042 | 4.97E-11 | 4.84E-09 |
| AT5G15960 | cold and ABA inducible prot | -2.174294159 | 5.34E-11 | 5.19E-09 |
| AT1G33170 | S-adenosyl-L-methionine-de | -4.366022073 | 7.58E-11 | 7.19E-09 |
| AT2G18890 | Protein kinase superfamily | -3.972457323 | 7.68E-11 | 7.26E-09 |
| AT1G45130 | beta-galactosidase 5;(sourc | -2.825136034 | 9.29E-11 | 8.70E-09 |
| AT5G20740 | Plant invertase/pectin meth | -4.564393799 | 9.63E-11 | 8.99E-09 |
| AT1G11545 | xyloglucan endotransglucos | -4.194364608 | 1.03E-10 | 9.52E-09 |
| AT3G56120 | S-adenosyl-L-methionine-de | -2.239234588 | 1.09E-10 | 1.00E-08 |
| AT5G26670 | Pectinacetylesterase family | -4.171829967 | 1.17E-10 | 1.07E-08 |
| AT3G27360 | Histone superfamily protein | -2.897350162 | 1.20E-10 | 1.09E-08 |
| AT4G26530 | Aldolase superfamily protei | -2.149775436 | 1.25E-10 | 1.12E-08 |
| AT2G34060 | Peroxidase superfamily pro | -3.722914337 | 1.51E-10 | 1.33E-08 |
| AT4G28250 | putative beta-expansin/alle | -3.445291801 | 1.77E-10 | 1.53E-08 |
| AT2G27385 | Pollen Ole e 1 allergen and | -3.049148596 | 1.79E-10 | 1.54E-08 |
| AT1G13250 | Encodes a protein with put | -2.738795776 | 1.84E-10 | 1.58E-08 |
| AT5G05890 | Encodes a nicotinate-N-glyc | -4.807499192 | 2.08E-10 | 1.78E-08 |
| AT5G65730 | xyloglucan endotransglucos | -2.821157452 | 2.16E-10 | 1.83E-08 |
| AT2G39900 | Encodes a member of the A | -2.48872313 | 2.30E-10 | 1.94E-08 |
| AT3G53190 | Pectin lyase-like superfamil | -3.594777504 | 2.36E-10 | 1.97E-08 |
| AT4G34760 | SAUR-like auxin-responsive | -3.047190882 | 2.54E-10 | 2.11E-08 |
| AT2G28470 | putative beta-galactosidase | -2.21926978 | 2.92E-10 | 2.41E-08 |
| AT4G30610 | Encodes a secreted glycosyl | -3.353763292 | 3.48E-10 | 2.84E-08 |
| AT5G47500 | predicted to encode a pecti | -5.983837132 | 5.49E-10 | 4.33E-08 |
| AT2G39700 | putative expansin. Naming | -2.168929059 | 5.81E-10 | 4.56E-08 |
| AT3G27400 | Encodes a pectate lyase inv | -2.135780554 | 5.98E-10 | 4.68E-08 |
| AT3G23890 | Encodes a topoisomerase II | -4.511432401 | 6.53E-10 | 5.06E-08 |
| AT4G33467 | hypothetical protein;(source | -5.110982102 | 7.55E-10 | 5.75E-08 |
| AT5G66230 | Chalcone-flavanone isomera | -5.171674786 | 7.89E-10 | 5.98E-08 |
| AT3G23670 | Microtubule motor kinesin I | -5.343373311 | 8.75E-10 | 6.55E-08 |
| AT5G61290 | Flavin-binding monooxygen | -2.987617278 | 8.79E-10 | 6.55E-08 |
| AT2G21540 | SEC14-like 3;(source:Arapor | -2.803753281 | 9.43E-10 | 6.94E-08 |
| AT4G35350 | tracheary element vacuolar | -2.027046813 | 1.11E-09 | 8.06E-08 |
| AT5G45820 | Encodes a CBL-interacting s | -7.30985952 | 1.16E-09 | 8.38E-08 |
| AT5G48900 | Pectin lyase-like superfamil | -3.696192397 | 1.17E-09 | 8.38E-08 |
| AT1G45191 | beta-glucosidase related pr | -2.238776956 | 1.18E-09 | 8.48E-08 |
| AT5G17160 | aspartic/glutamic acid-rich | -4.43863585 | 1.20E-09 | 8.54E-08 |
| AT3G51080 | Encodes a member of the G | -2.164115392 | 1.23E-09 | 8.79E-08 |
| AT3G25980 | Encodes MAD2 (MITOTIC AI | -3.910166655 | 1.31E-09 | 9.17E-08 |
| AT3G14170 | hypothetical protein (DUF93 | -2.378629042 | 1.44E-09 | 9.98E-08 |
| AT3G10080 | RmlC-like cupins superfami | -2.507527903 | 1.44E-09 | 9.99E-08 |

| | | | | |
|---|---|---|---|---|
| AT1G20390 | transposable_element_gen | -2.499526653 | 1.49E-09 | 1.03E-07 |
| AT2G23560 | Encodes a protein shown to | -2.387822914 | 1.74E-09 | 1.18E-07 |
| AT2G28740 | histone 4histone 4 | -2.971967352 | 1.74E-09 | 1.18E-07 |
| AT4G32000 | Protein kinase superfamily | -2.067048484 | 1.78E-09 | 1.21E-07 |
| AT3G05470 | Actin-binding FH2 (formin h | -4.864711207 | 1.85E-09 | 1.25E-07 |
| AT2G47500 | P-loop nucleoside triphosph | -2.21981304 | 1.89E-09 | 1.27E-07 |
| AT1G60390 | polygalacturonase 1;(source | -3.516838238 | 1.97E-09 | 1.31E-07 |
| AT1G72230 | Cupredoxin superfamily pro | -4.570392933 | 2.01E-09 | 1.33E-07 |
| AT1G26770 | Encodes an expansin. Nami | -2.113280806 | 2.10E-09 | 1.39E-07 |
| AT2G05100 | Lhcb2.1 protein encoding a | -5.768711166 | 2.14E-09 | 1.41E-07 |
| AT4G30650 | Low temperature and salt r | -2.395430107 | 2.14E-09 | 1.41E-07 |
| AT5G12050 | rho GTPase-activating prote | -3.429655716 | 2.35E-09 | 1.53E-07 |
| AT1G76240 | DUF241 domain protein (DU | -3.742689776 | 2.38E-09 | 1.55E-07 |
| AT5G50740 | Heavy metal transport/detc | -3.124453143 | 2.41E-09 | 1.57E-07 |
| AT5G04160 | UUAT1 is a UDP-Uronic acid | -2.044306349 | 2.73E-09 | 1.75E-07 |
| AT4G02290 | glycosyl hydrolase 9B13;(so | -7.782112245 | 2.86E-09 | 1.83E-07 |
| AT1G02820 | Late embryogenesis abunda | -2.637370674 | 2.98E-09 | 1.89E-07 |
| AT2G45470 | FASCICLIN-like arabinogalac | -2.308795457 | 3.03E-09 | 1.91E-07 |
| AT2G34430 | Photosystem II type I chloro | -5.672890778 | 3.10E-09 | 1.96E-07 |
| AT1G20190 | member of Alpha-Expansin | -4.980420638 | 3.26E-09 | 2.04E-07 |
| AT2G36200 | P-loop containing nucleosid | -3.803973861 | 3.55E-09 | 2.20E-07 |
| AT1G29910 | member of Chlorophyll a/b- | -5.345764751 | 3.73E-09 | 2.29E-07 |
| AT3G51740 | encodes a leucine-repeat re | -3.539694486 | 4.06E-09 | 2.48E-07 |
| AT2G28620 | Mutants have radially swoll | -3.381587082 | 4.26E-09 | 2.58E-07 |
| AT5G46600 | aluminum activated malate | -4.260948059 | 4.29E-09 | 2.59E-07 |
| AT1G44110 | Cyclin A1;(source:Araport11 | -5.389852667 | 4.84E-09 | 2.87E-07 |
| AT5G55820 | Encodes a plant ortholog of | -3.977470142 | 4.87E-09 | 2.88E-07 |
| AT3G44990 | Encodes a xyloglucan endot | -6.514366887 | 5.00E-09 | 2.95E-07 |
| AT5G26850 | Uncharacterized protein;(sc | -2.535451387 | 5.45E-09 | 3.21E-07 |
| AT3G16240 | Delta tonoplast intrinsic pro | -3.440931906 | 5.49E-09 | 3.22E-07 |
| AT1G76310 | core cell cycle genes | -4.398239144 | 5.59E-09 | 3.26E-07 |
| AT2G33330 | Encodes a plasmodesmal p | -3.239419996 | 5.62E-09 | 3.27E-07 |
| AT1G72970 | Originally identified as a m | -2.463196896 | 6.12E-09 | 3.53E-07 |
| AT1G72670 | IQ-domain 8;(source:Arapor | -5.026143766 | 6.85E-09 | 3.90E-07 |
| AT1G80080 | Encodes a transmembrane | -4.47706138 | 6.95E-09 | 3.95E-07 |
| AT1G20060 | ATP binding microtubule m | -3.299930954 | 7.45E-09 | 4.20E-07 |
| AT2G38620 | Encodes a member of a pla | -4.621419955 | 7.47E-09 | 4.20E-07 |
| AT1G17190 | Encodes glutathione transfe | -2.38044323 | 7.74E-09 | 4.33E-07 |
| AT1G10640 | Pectin lyase-like superfamil | -6.281584953 | 8.42E-09 | 4.65E-07 |
| AT2G16850 | plasma membrane intrinsic | -2.877703039 | 8.50E-09 | 4.68E-07 |
| AT2G35860 | FASCICLIN-like arabinogalac | -2.265796662 | 9.23E-09 | 5.06E-07 |
| AT2G15970 | encodes an alpha form of a | -2.255475995 | 9.47E-09 | 5.16E-07 |
| AT2G30820 | aspartyl/glutamyl-tRNA(Asn | -3.540079947 | 9.51E-09 | 5.16E-07 |
| AT4G13710 | Pectin lyase-like superfamil | -2.423338392 | 1.02E-08 | 5.52E-07 |
| AT2G27402 | plastid transcriptionally acti | -3.359496016 | 1.12E-08 | 5.97E-07 |
| AT2G02100 | Predicted to encode a PR (p | -3.886488524 | 1.13E-08 | 6.01E-07 |

| | | | | |
|---|---|---|---|---|
| AT5G51720 | Encodes a protein with bioc | -3.82536702 | 1.23E-08 | 6.47E-07 |
| AT1G23800 | Encodes a mitochondrial ald | -2.218740735 | 1.34E-08 | 6.99E-07 |
| AT4G14330 | P-loop containing nucleosid | -5.326430772 | 1.37E-08 | 7.14E-07 |
| AT5G45960 | GDSL-motif esterase/acyltr | -6.600831754 | 1.38E-08 | 7.19E-07 |
| AT5G03760 | encodes a beta-mannan syr | -2.029258832 | 1.41E-08 | 7.30E-07 |
| AT4G33810 | Glycosyl hydrolase superfar | -2.093903837 | 1.43E-08 | 7.37E-07 |
| AT5G16250 | transmembrane protein;(sc | -3.868513972 | 1.46E-08 | 7.50E-07 |
| AT3G53680 | Acyl-CoA N-acyltransferase | -2.534659488 | 1.47E-08 | 7.55E-07 |
| AT1G23205 | Plant invertase/pectin meth | -2.977747803 | 1.61E-08 | 8.08E-07 |
| AT1G78120 | Encodes one of the 36 carbo | -2.359535666 | 1.64E-08 | 8.20E-07 |
| AT5G55520 | kinesin-like protein;(source: | -5.503713589 | 1.65E-08 | 8.24E-07 |
| AT4G33270 | Encodes a CDC20 protein th | -4.661953026 | 1.67E-08 | 8.33E-07 |
| AT1G74890 | Encodes a nuclear response | -4.928386979 | 1.68E-08 | 8.34E-07 |
| AT4G05190 | ATK5 encodes a kinesin pro | -4.260237039 | 1.69E-08 | 8.34E-07 |
| AT4G27595 | Encodes a microtubule-asso | -2.143849818 | 1.70E-08 | 8.34E-07 |
| AT4G29020 | glycine-rich protein;(source | -5.501382296 | 1.76E-08 | 8.61E-07 |
| AT2G30432 | Encodes TRICHOMELESS1 (T | -4.657195161 | 1.85E-08 | 8.96E-07 |
| AT2G19170 | Encodes a novel subtilisin-li | -2.52305568 | 1.85E-08 | 8.96E-07 |
| AT1G18250 | encodes a thaumatin-like pr | -4.265980329 | 1.89E-08 | 9.11E-07 |
| AT5G23910 | ATP binding microtubule m | -3.654538174 | 1.93E-08 | 9.30E-07 |
| AT1G14350 | Encodes a putative MYB tra | -2.128661557 | 1.98E-08 | 9.51E-07 |
| AT5G60020 | LAC17 appears to have lacc | -2.025250721 | 1.99E-08 | 9.53E-07 |
| AT1G50010 | Encodes alpha-2,4 tubulin. | -2.596814317 | 2.02E-08 | 9.63E-07 |
| AT1G25510 | Eukaryotic aspartyl protease | -4.489782493 | 2.03E-08 | 9.65E-07 |
| AT1G70710 | endo-1,4-beta-glucanase. In | -2.509055341 | 2.05E-08 | 9.72E-07 |
| AT3G53900 | Encodes UPP, a plastidial ur | -2.135365721 | 2.16E-08 | 1.02E-06 |
| AT5G67270 | encodes a homolog of anim | -3.671479197 | 2.20E-08 | 1.03E-06 |
| AT5G46700 | Encodes a transmembrane | -2.888996709 | 2.20E-08 | 1.03E-06 |
| AT2G05790 | O-Glycosyl hydrolases famil | -2.447438253 | 2.22E-08 | 1.03E-06 |
| AT4G14940 | atao1 gene of Arabidopsis t | -3.801296271 | 2.42E-08 | 1.11E-06 |
| AT5G15780 | Pollen Ole e 1 allergen and | -4.758061263 | 2.43E-08 | 1.11E-06 |
| AT5G55950 | Nucleotide/sugar transport | -2.880486364 | 2.57E-08 | 1.17E-06 |
| AT5G60150 | hypothetical protein;(source | -4.596353962 | 2.60E-08 | 1.18E-06 |
| AT1G78820 | D-mannose binding lectin p | -2.797235307 | 2.65E-08 | 1.20E-06 |
| AT1G57750 | Encodes a CYP96A15, midch | -9.551979495 | 2.84E-08 | 1.28E-06 |
| AT3G55660 | Encodes a member of KPP-l | -5.354439597 | 2.92E-08 | 1.31E-06 |
| AT3G20260 | DUF1666 family protein (DU | -4.525883064 | 2.98E-08 | 1.33E-06 |
| AT1G02640 | encodes a protein similar to | -2.413165147 | 3.04E-08 | 1.35E-06 |
| AT5G18430 | GDSL-motif esterase/acyltr | -2.504233117 | 3.60E-08 | 1.59E-06 |
| AT1G15570 | A2-type cyclin. Negatively r | -3.321895258 | 3.62E-08 | 1.59E-06 |
| AT4G16740 | Encodes an (E,E)-alpha-farr | -3.614240883 | 4.45E-08 | 1.95E-06 |
| AT3G20150 | Kinesin motor family protei | -3.510753285 | 4.51E-08 | 1.97E-06 |
| AT3G51720 | WEB family protein (DUF82 | -3.279631492 | 4.61E-08 | 2.00E-06 |
| AT1G51060 | Encodes HTA10, a histone H | -2.122390917 | 4.65E-08 | 2.01E-06 |
| AT5G23940 | Encodes PERMEABLE LEAVE | -5.62558021 | 5.17E-08 | 2.22E-06 |
| AT5G02760 | Encodes a phosphatase tha | -5.244663081 | 5.96E-08 | 2.52E-06 |

| | | | | |
|---|---|---|---|---|
| AT2G42380 | Encodes a member of the B | -3.245182675 | 6.13E-08 | 2.57E-06 |
| AT3G45430 | Concanavalin A-like lectin p | -4.697735658 | 6.14E-08 | 2.57E-06 |
| AT1G58070 | WEB family protein;(source | -2.339118113 | 6.29E-08 | 2.62E-06 |
| AT1G02730 | Encodes a gene similar to c | -4.222731501 | 6.40E-08 | 2.66E-06 |
| AT3G45930 | Histone superfamily proteir | -2.515860102 | 6.47E-08 | 2.68E-06 |
| AT2G40670 | response regulator 16 | -2.254335136 | 6.62E-08 | 2.74E-06 |
| AT5G51600 | Mutant has defective roots. | -4.516955021 | 6.83E-08 | 2.82E-06 |
| AT1G12845 | transmembrane protein;(so | -3.172507589 | 7.04E-08 | 2.90E-06 |
| AT1G18370 | Encodes a kinesin HINKEL. F | -4.298840416 | 7.09E-08 | 2.91E-06 |
| AT1G50240 | The FUSED (FU) gene belor | -3.714253394 | 7.11E-08 | 2.92E-06 |
| AT4G36240 | Encodes a member of the G | -4.031012214 | 7.81E-08 | 3.18E-06 |
| AT3G15550 | trichohyalin;(source:Araport | -4.401789274 | 8.14E-08 | 3.29E-06 |
| AT1G21540 | AMP-dependent synthetase | -2.231379274 | 8.48E-08 | 3.40E-06 |
| AT4G31290 | ChaC-like family protein;(so | -2.629208132 | 8.74E-08 | 3.48E-06 |
| AT5G11590 | encodes a member of the D | -3.257733678 | 8.74E-08 | 3.48E-06 |
| AT5G49800 | Polyketide cyclase/dehydras | -2.245345006 | 8.92E-08 | 3.54E-06 |
| AT5G54270 | Lhcb3 protein is a compone | -4.520997717 | 9.10E-08 | 3.60E-06 |
| AT3G50410 | Arabidopsis Dof protein con | -2.065684998 | 9.49E-08 | 3.72E-06 |
| AT3G15950 | Similar to TSK-associating p | -2.538725957 | 9.55E-08 | 3.72E-06 |
| AT5G56120 | RNA polymerase II elongati | -4.576127504 | 9.56E-08 | 3.72E-06 |
| AT4G19380 | Long-chain fatty alcohol del | -5.794297085 | 1.00E-07 | 3.87E-06 |
| AT1G78430 | Encodes RIP2 (ROP interact | -4.556158648 | 1.02E-07 | 3.93E-06 |
| AT1G09350 | Predicted to encode a galac | -5.816865159 | 1.04E-07 | 3.98E-06 |
| AT3G55580 | TCF1 encodes a member of | -8.89778691 | 1.04E-07 | 3.98E-06 |
| AT3G63200 | PATATIN-like protein 9;(sou | -4.370698121 | 1.04E-07 | 3.98E-06 |
| AT1G31330 | Encodes subunit F of photos | -2.20440661 | 1.05E-07 | 4.01E-06 |
| AT2G28870 | cyclin-dependent kinase inh | -2.423838269 | 1.15E-07 | 4.38E-06 |
| AT5G59970 | Histone superfamily proteir | -3.051313068 | 1.25E-07 | 4.72E-06 |
| AT5G54190 | light-dependent NADPH:pro | -6.226506787 | 1.32E-07 | 4.93E-06 |
| AT3G12610 | Plays role in DNA-damage r | -2.947352514 | 1.32E-07 | 4.93E-06 |
| AT2G46760 | D-arabinono-1,4-lactone oxi | -2.514231195 | 1.36E-07 | 5.05E-06 |
| AT5G25090 | early nodulin-like protein 13 | -4.305433802 | 1.37E-07 | 5.07E-06 |
| AT5G25810 | encodes a member of the D | -3.786346108 | 1.38E-07 | 5.11E-06 |
| AT1G75640 | Encodes a Leucine-Rich Rep | -4.405992179 | 1.44E-07 | 5.32E-06 |
| AT2G05070 | Encodes Lhcb2.2. Belongs t | -5.248103204 | 1.45E-07 | 5.33E-06 |
| AT3G47470 | Encodes a chlorophyll a/b-b | -3.603623528 | 1.46E-07 | 5.36E-06 |
| AT3G20670 | Encodes HTA13, a histone H | -2.654627761 | 1.48E-07 | 5.42E-06 |
| AT1G05835 | PHD finger protein;(source:/ | -2.585419957 | 1.55E-07 | 5.59E-06 |
| AT4G22860 | Cell cycle regulated microtu | -3.917828132 | 1.55E-07 | 5.60E-06 |
| AT3G05730 | Encodes a defensin-like (DE | -3.592154623 | 1.63E-07 | 5.82E-06 |
| AT4G23020 | hypothetical protein;(source | -3.105101323 | 1.67E-07 | 5.97E-06 |
| AT5G13140 | Pollen Ole e 1 allergen and | -2.57621234 | 1.67E-07 | 5.98E-06 |
| AT2G34170 | hypothetical protein (DUF68 | -2.702214069 | 1.72E-07 | 6.11E-06 |
| AT5G64040 | Encodes the only subunit of | -2.279260183 | 1.80E-07 | 6.36E-06 |
| AT2G39705 | ROTUNDIFOLIA like 8;(sour | -2.810193896 | 1.83E-07 | 6.44E-06 |
| AT1G56720 | Protein kinase superfamily | -2.381149539 | 1.86E-07 | 6.51E-06 |

| | | | | |
|---|---|---|---|---|
| AT1G76540 | Encodes a cyclin-dependent | -3.360997769 | 2.02E-07 | 6.98E-06 |
| AT2G42570 | Encodes a member of the T | -3.428596169 | 2.07E-07 | 7.14E-06 |
| AT1G01600 | Encodes a member of the C | -5.403842615 | 2.10E-07 | 7.26E-06 |
| AT1G12080 | Vacuolar calcium-binding p | -3.471630277 | 2.22E-07 | 7.61E-06 |
| AT3G21950 | S-adenosyl-L-methionine-de | -2.137343584 | 2.29E-07 | 7.80E-06 |
| AT4G18970 | GDSL-motif esterase/acylt | -2.700429938 | 2.49E-07 | 8.40E-06 |
| AT1G08560 | member of SYP11 syntaxin | -3.599095729 | 2.52E-07 | 8.48E-06 |
| AT3G58650 | Encodes a member of the T | -4.737919884 | 2.53E-07 | 8.50E-06 |
| AT4G02850 | DAAR1 encodes a PLP-indep | -3.65889868 | 2.56E-07 | 8.56E-06 |
| AT3G17360 | PHRAGMOPLAST ORIENTIN | -3.910703554 | 2.59E-07 | 8.62E-06 |
| AT1G61520 | PSI type III chlorophyll a/b-b | -2.231925136 | 2.69E-07 | 8.90E-06 |
| AT4G03100 | Rho GTPase activating prot | -4.614519754 | 2.86E-07 | 9.36E-06 |
| AT1G69700 | Part of the AtHVA22 family. | -2.4764041 | 2.90E-07 | 9.44E-06 |
| AT3G57060 | binding protein;(source:Ara | -2.45415045 | 3.00E-07 | 9.71E-06 |
| AT1G10682 | Has been identified as a tra | -2.552552763 | 3.27E-07 | 1.05E-05 |
| AT5G56490 | D-arabinono-1,4-lactone oxi | -3.825998144 | 3.34E-07 | 1.07E-05 |
| AT5G57123 | hypothetical protein;(source | -3.603354338 | 3.33E-07 | 1.07E-05 |
| AT4G25480 | encodes a member of the D | -5.104990307 | 3.40E-07 | 1.09E-05 |
| AT4G11320 | Papain family cysteine prot | -3.6974208 | 3.48E-07 | 1.11E-05 |
| AT2G28790 | Pathogenesis-related thaum | -6.210896243 | 3.50E-07 | 1.11E-05 |
| AT2G19920 | RNA-dependent RNA polym | -3.543372954 | 3.56E-07 | 1.12E-05 |
| AT2G21650 | RSM1 is a member of a sm | -3.175109785 | 3.58E-07 | 1.13E-05 |
| AT4G21760 | beta-glucosidase 47;(source | -2.01828553 | 3.72E-07 | 1.17E-05 |
| AT4G17000 | neurofilament heavy protei | -3.419480416 | 3.73E-07 | 1.17E-05 |
| AT5G62700 | encodes tubulin beta-2/bet | -2.047705427 | 3.75E-07 | 1.17E-05 |
| AT1G16630 | transmembrane protein;(so | -4.058005186 | 3.98E-07 | 1.24E-05 |
| AT2G45190 | Encodes a member of the Y | -2.201013811 | 4.12E-07 | 1.28E-05 |
| AT2G07170 | ARM repeat superfamily pro | -4.100756438 | 4.23E-07 | 1.31E-05 |
| AT2G03090 | member of Alpha-Expansin | -6.272104359 | 4.25E-07 | 1.31E-05 |
| AT5G48310 | portal protein;(source:Arap | -3.964832989 | 4.44E-07 | 1.36E-05 |
| AT2G04780 | fasciclin-like arabinogalacta | -2.193974059 | 4.50E-07 | 1.38E-05 |
| AT5G49160 | Encodes a cytosine methylt | -2.163011857 | 4.66E-07 | 1.42E-05 |
| AT5G02890 | Encodes a protein with simi | -2.331120117 | 4.84E-07 | 1.46E-05 |
| AT5G27360 | Encodes a sugar-porter fam | -3.422419434 | 4.85E-07 | 1.47E-05 |
| AT4G08150 | A member of class I knottec | -6.978442537 | 5.15E-07 | 1.54E-05 |
| AT3G14890 | phosphoesterase;(source:Ar | -2.164212222 | 5.16E-07 | 1.54E-05 |
| AT3G58120 | Encodes a member of the B | -3.518463905 | 5.24E-07 | 1.56E-05 |
| AT1G20610 | Cyclin B2;(source:Araport11 | -3.93775981 | 5.44E-07 | 1.61E-05 |
| AT1G03780 | Homolog of vertebrate TPX2 | -3.650621556 | 5.61E-07 | 1.65E-05 |
| AT1G63100 | GRAS family transcription f | -6.604687125 | 5.73E-07 | 1.66E-05 |
| AT2G44740 | cyclin p4;(source:Araport11) | -2.603379572 | 5.94E-07 | 1.72E-05 |
| AT5G54585 | hypothetical protein;(source | -2.370744101 | 6.11E-07 | 1.76E-05 |
| AT5G62550 | microtubule-associated futs | -3.525648856 | 6.21E-07 | 1.79E-05 |
| AT1G65710 | serine/arginine repetitive m | -4.891205388 | 6.60E-07 | 1.88E-05 |
| AT1G77110 | Rate-limiting factor in satu | -2.330960904 | 6.89E-07 | 1.94E-05 |
| AT1G75090 | DNA glycosylase superfamil | -2.598770281 | 6.92E-07 | 1.95E-05 |

| | | | | |
|---|---|---|---|---|
| AT1G54385 | At1G54385 encodes the pla | -4.333866992 | 7.07E-07 | 1.98E-05 |
| AT1G47670 | Transmembrane amino aci | -2.119606397 | 8.07E-07 | 2.22E-05 |
| AT2G14890 | putative proline-rich protei | -2.372158142 | 8.16E-07 | 2.25E-05 |
| AT1G75780 | beta tubulin gene downreg | -3.743914935 | 8.30E-07 | 2.28E-05 |
| AT2G42840 | Encodes a putative extracel | -6.645595575 | 8.32E-07 | 2.28E-05 |
| AT5G13840 | FIZZY-related 3;(source:Ara | -2.908069353 | 8.44E-07 | 2.31E-05 |
| AT3G03130 | lisH domain-like protein;(so | -5.00502369 | 8.46E-07 | 2.32E-05 |
| AT4G21820 | binding / calmodulin bindin | -3.616090744 | 8.66E-07 | 2.36E-05 |
| AT3G17680 | Kinase interacting (KIP1-like | -3.593207882 | 8.82E-07 | 2.39E-05 |
| AT4G38860 | SAUR-like auxin-responsive | -4.607105228 | 8.81E-07 | 2.39E-05 |
| AT4G26660 | kinesin-like protein;(source: | -3.963563165 | 8.89E-07 | 2.40E-05 |
| AT2G25060 | early nodulin-like protein 14 | -4.212532306 | 9.13E-07 | 2.46E-05 |
| AT4G21270 | Encodes a kinesin-like moto | -4.835551049 | 9.15E-07 | 2.46E-05 |
| AT1G78865 | other_RNA;(source:Araport | -5.84441975 | 9.30E-07 | 2.49E-05 |
| AT1G21500 | hypothetical protein;(source | -2.623724456 | 9.46E-07 | 2.53E-05 |
| AT4G23800 | Encodes a protein containin | -3.933103991 | 9.64E-07 | 2.57E-05 |
| AT2G37640 | member of Alpha-Expansin | -2.969766609 | 9.87E-07 | 2.63E-05 |
| AT5G41140 | Myosin heavy chain-related | -2.297800867 | 1.01E-06 | 2.67E-05 |
| AT4G34220 | Encodes a receptor like kina | -2.255855151 | 1.02E-06 | 2.70E-05 |
| AT2G34420 | Photosystem II type I chloro | -3.116587989 | 1.03E-06 | 2.73E-05 |
| AT5G39550 | Encodes the VIM3/ORTH1 p | -2.973913908 | 1.06E-06 | 2.79E-05 |
| AT4G12980 | Auxin-responsive family pro | -2.435839883 | 1.12E-06 | 2.92E-05 |
| AT5G56580 | Encodes a member of the M | -2.165746511 | 1.12E-06 | 2.92E-05 |
| AT2G45080 | cyclin p3;(source:Araport11 | -3.771177695 | 1.14E-06 | 2.95E-05 |
| AT1G25450 | Encodes KCS5, a member of | -2.056244751 | 1.16E-06 | 3.00E-05 |
| AT5G35970 | P-loop containing nucleosid | -2.853645724 | 1.18E-06 | 3.06E-05 |
| AT3G02110 | serine carboxypeptidase-lik | -2.007863298 | 1.20E-06 | 3.10E-05 |
| AT5G60490 | Encodes a member of fasci | -3.74378708 | 1.23E-06 | 3.15E-05 |
| AT2G21140 | Proline-rich protein express | -3.984706045 | 1.24E-06 | 3.18E-05 |
| AT4G15830 | ARM repeat superfamily pr | -4.724380162 | 1.25E-06 | 3.21E-05 |
| AT5G45700 | Haloacid dehalogenase-like | -4.909228589 | 1.30E-06 | 3.32E-05 |
| AT1G10060 | encodes a mitochondrial bra | -2.961880417 | 1.37E-06 | 3.46E-05 |
| AT3G20820 | Leucine-rich repeat (LRR) fa | -2.278449531 | 1.42E-06 | 3.57E-05 |
| AT3G10180 | P-loop containing nucleosid | -3.015667138 | 1.44E-06 | 3.62E-05 |
| AT4G29030 | Putative membrane lipopro | -8.497564252 | 1.45E-06 | 3.63E-05 |
| AT4G02800 | GRIP/coiled-coil protein;(so | -4.448276333 | 1.54E-06 | 3.84E-05 |
| AT3G48970 | Heavy metal transport/detc | -4.198032827 | 1.62E-06 | 4.02E-05 |
| AT5G57130 | Encodes a member of an ei | -2.166281954 | 1.68E-06 | 4.14E-05 |
| AT5G33370 | GDSL-motif esterase/acylt | -7.547023817 | 1.69E-06 | 4.16E-05 |
| AT1G07180 | Internal NAD(P)H dehydroge | -2.804162107 | 1.73E-06 | 4.25E-05 |
| AT1G71030 | Encodes a putative myb fan | -5.022837316 | 1.75E-06 | 4.29E-05 |
| AT5G60660 | A member of the plasma m | -2.36477995 | 1.82E-06 | 4.43E-05 |
| AT5G52290 | Encodes a protein with simi | -3.720397854 | 1.83E-06 | 4.44E-05 |
| AT5G65640 | bHLH093/NFL encodes a bH | -3.609663679 | 1.85E-06 | 4.49E-05 |
| AT1G09470 | NEAP3 is a member of a sm | -5.037041593 | 1.91E-06 | 4.62E-05 |
| AT2G21050 | Encodes LAX2 (LIKE AUXIN I | -2.687042083 | 1.97E-06 | 4.74E-05 |

| | | | | |
|---|---|---|---|---|
| AT4G38400 | member of EXPANSIN-LIKE. | -2.094532068 | 1.97E-06 | 4.74E-05 |
| AT4G34900 | xanthine dehydrogenase 2;( | -3.830498639 | 2.03E-06 | 4.86E-05 |
| AT2G25900 | Encodes a protein with two | -4.007500557 | 2.04E-06 | 4.88E-05 |
| AT1G10930 | DNA helicase involved in th | -2.226010228 | 2.15E-06 | 5.11E-05 |
| AT1G20010 | beta tubulinbeta tubulin | -2.053909674 | 2.19E-06 | 5.18E-05 |
| AT1G31173 | Encodes a microRNA that ta | -3.206117815 | 2.27E-06 | 5.37E-05 |
| AT2G36050 | ovate family protein 15;(sou | -4.182870923 | 2.31E-06 | 5.44E-05 |
| AT1G04110 | Initially identified as a muta | -3.203575391 | 2.35E-06 | 5.52E-05 |
| AT3G05600 | Encodes a cytosolic epoxide | -5.926427366 | 2.44E-06 | 5.69E-05 |
| AT5G49170 | hypothetical protein;(source | -5.121812085 | 2.45E-06 | 5.71E-05 |
| AT5G03390 | hypothetical protein (DUF29 | -2.088449807 | 2.46E-06 | 5.72E-05 |
| AT3G08770 | Predicted to encode a PR (p | -4.977340719 | 2.47E-06 | 5.74E-05 |
| AT1G70210 | Encodes a D-type cyclin that | -2.968740976 | 2.52E-06 | 5.85E-05 |
| AT3G19050 | PHRAGMOPLAST ORIENTIN | -3.675629583 | 2.53E-06 | 5.85E-05 |
| AT1G06080 | Encodes a protein homolog | -8.628554221 | 2.56E-06 | 5.92E-05 |
| AT1G49870 | myosin-2 heavy chain-like p | -3.743166176 | 2.59E-06 | 5.96E-05 |
| AT1G52270 | hypothetical protein;(source | -3.425373876 | 2.69E-06 | 6.17E-05 |
| AT3G02640 | transmembrane protein;(so | -4.033261925 | 2.71E-06 | 6.22E-05 |
| AT3G15720 | Pectin lyase-like superfamil | -2.644268205 | 2.77E-06 | 6.34E-05 |
| AT2G40820 | stomatal closure actin-bind | -2.162019925 | 2.80E-06 | 6.38E-05 |
| AT3G23085 | transposable_element_gen | -6.464610133 | 2.81E-06 | 6.41E-05 |
| AT1G08340 | Rho GTPase activating prot | -2.135274796 | 2.84E-06 | 6.46E-05 |
| AT3G08920 | Rhodanese/Cell cycle contr | -2.09362798 | 2.90E-06 | 6.58E-05 |
| AT1G24260 | Member of the MADs box t | -2.658365296 | 2.91E-06 | 6.59E-05 |
| AT3G54260 | Encodes a member of the T | -2.484651071 | 3.00E-06 | 6.76E-05 |
| AT5G02170 | Transmembrane amino acid | -2.376360752 | 3.02E-06 | 6.81E-05 |
| AT1G80370 | Encodes a A2-type cyclin. C | -4.110122371 | 3.07E-06 | 6.89E-05 |
| AT5G63180 | Pectin lyase-like superfamil | -2.087838647 | 3.13E-06 | 7.00E-05 |
| AT3G26125 | encodes a protein with cyto | -2.405093191 | 3.18E-06 | 7.10E-05 |
| AT2G22170 | Lipase/lipooxygenase, PLAT | -2.88800564 | 3.25E-06 | 7.20E-05 |
| AT1G78170 | E3 ubiquitin-protein ligase;( | -3.484759594 | 3.45E-06 | 7.61E-05 |
| AT1G09450 | Encodes a protein kinase th | -3.750177788 | 3.45E-06 | 7.61E-05 |
| AT2G40475 | hypothetical protein;(source | -2.830682852 | 3.51E-06 | 7.70E-05 |
| AT4G18640 | Required for root hair elong | -2.307942476 | 3.57E-06 | 7.81E-05 |
| AT5G05940 | Encodes a member of KPP-l | -3.280842323 | 3.60E-06 | 7.87E-05 |
| AT3G44940 | enabled-like protein (DUF16 | -2.694607847 | 3.85E-06 | 8.33E-05 |
| AT4G14770 | TESMIN/TSO1-like CXC 2;(s | -2.816054992 | 3.89E-06 | 8.41E-05 |
| AT5G04820 | ovate family protein 13;(sou | -2.518406958 | 4.00E-06 | 8.57E-05 |
| AT5G67280 | receptor-like kinase;(source | -2.035280415 | 4.09E-06 | 8.73E-05 |
| AT4G03010 | RNI-like superfamily protein | -2.075327661 | 4.10E-06 | 8.73E-05 |
| AT5G17780 | alpha/beta-Hydrolases supe | -3.062191421 | 4.21E-06 | 8.95E-05 |
| AT2G22610 | Di-glucose binding protein v | -3.71473184 | 4.24E-06 | 9.00E-05 |
| AT3G13175 | transmembrane protein;(so | -4.521364652 | 4.28E-06 | 9.08E-05 |
| AT3G05890 | Low temperature and salt r | -3.471337179 | 4.32E-06 | 9.12E-05 |
| AT1G52910 | fiber (DUF1218);(source:Ara | -2.696586706 | 4.39E-06 | 9.21E-05 |
| AT1G55200 | kinase with adenine nucleot | -3.896371427 | 4.40E-06 | 9.21E-05 |

| | | | | |
|---|---|---|---|---|
| AT2G13820 | Bifunctional inhibitor/lipid-1 | -3.649141935 | 4.58E-06 | 9.53E-05 |
| AT2G04235 | hypothetical protein;(source | -2.153850878 | 4.65E-06 | 9.67E-05 |
| AT4G31840 | early nodulin-like protein 15 | -3.649821231 | 4.82E-06 | 9.95E-05 |
| AT2G32280 | Encodes a member of a pla | -2.58262684 | 4.88E-06 | 0.000100612 |
| AT5G55830 | Concanavalin A-like lectin p | -3.378840202 | 4.88E-06 | 0.000100612 |
| AT5G54200 | Transducin/WD40 repeat-li | -2.371624442 | 4.92E-06 | 0.000101251 |
| AT2G06850 | endoxyloglucan transferase | -2.12839006 | 4.94E-06 | 0.00010147 |
| AT1G06980 | 6,7-dimethyl-8-ribityllumaz | -4.999434269 | 5.03E-06 | 0.000102951 |
| AT1G23790 | dicer-like protein (DUF936) | -6.221906042 | 5.12E-06 | 0.000104439 |
| AT3G22760 | CXC domain containing TSO | -2.183227574 | 5.21E-06 | 0.000105863 |
| AT4G20430 | Subtilase family protein;(so | -2.792267588 | 5.27E-06 | 0.000106984 |
| AT2G04570 | GDSL-motif esterase/acylt | -2.024979121 | 5.41E-06 | 0.000109457 |
| AT3G15450 | aluminum induced protein v | -2.464038672 | 5.47E-06 | 0.00011068 |
| AT1G76740 | hypothetical protein;(source | -3.334434034 | 5.61E-06 | 0.00011285 |
| AT1G78440 | Encodes a gibberellin 2-oxic | -7.093793606 | 5.61E-06 | 0.00011285 |
| AT4G38070 | transcription factor bHLH13 | -3.945523182 | 5.60E-06 | 0.00011285 |
| AT2G16270 | transmembrane protein;(so | -3.496820879 | 5.76E-06 | 0.000115552 |
| AT5G06150 | Encodes a cyclin whose exp | -4.166120301 | 5.78E-06 | 0.000115819 |
| AT1G78970 | Lupeol synthase. Converts o | -2.055178182 | 5.81E-06 | 0.000116243 |
| AT4G33666 | hypothetical protein;(source | -2.871449326 | 5.83E-06 | 0.000116369 |
| AT5G57410 | Encodes a microtubule-asso | -2.143392229 | 5.99E-06 | 0.000119016 |
| AT1G13130 | Cellulase (glycosyl hydrolas | -4.470158866 | 6.15E-06 | 0.000121535 |
| AT4G28780 | GDSL-motif esterase/acylt | -3.662463535 | 6.17E-06 | 0.000121714 |
| AT5G19730 | Pectin lyase-like superfamil | -2.368048677 | 6.49E-06 | 0.00012746 |
| AT4G28310 | microtubule-associated pro | -3.044530048 | 6.88E-06 | 0.000134235 |
| AT1G35290 | Thioesterase superfamily p | -7.468859151 | 6.95E-06 | 0.000135304 |
| AT2G38110 | bifunctional sn-glycerol-3-p | -2.052466374 | 6.97E-06 | 0.000135685 |
| AT2G36570 | Leucine-rich repeat protein | -2.7863914 | 6.99E-06 | 0.000135711 |
| AT2G41540 | Encodes a protein with NAD | -2.304528427 | 7.23E-06 | 0.000139045 |
| AT1G53140 | Encodes DRP5A, a dynamin | -3.773166186 | 7.30E-06 | 0.000139964 |
| AT5G62230 | Encodes a receptor-like kina | -2.690829606 | 7.30E-06 | 0.000139964 |
| AT3G57500 | fission regulator-like protei | -6.361922847 | 7.60E-06 | 0.000144978 |
| AT5G40630 | Ubiquitin-like superfamily p | -5.176013929 | 7.61E-06 | 0.00014512 |
| AT3G18850 | lysophosphatidyl acyltransfe | -2.551477563 | 7.70E-06 | 0.000146696 |
| AT1G48100 | Pectin lyase-like superfamil | -3.977016317 | 7.73E-06 | 0.000147097 |
| AT5G51750 | subtilase 1.3;(source:Arapo | -2.494707593 | 7.90E-06 | 0.000149932 |
| AT1G79530 | Encodes one of the chloropl | -2.689385053 | 7.99E-06 | 0.000151537 |
| AT3G61920 | UvrABC system protein C;(s | -3.52421324 | 8.11E-06 | 0.000153164 |
| AT1G77270 | hypothetical protein;(source | -2.001585363 | 8.28E-06 | 0.000155643 |
| AT1G27120 | Encodes a Golgi-localized h | -2.623111896 | 8.33E-06 | 0.000156288 |
| AT4G35620 | Cyclin B2;(source:Araport11 | -3.935433308 | 8.37E-06 | 0.000156603 |
| AT1G09200 | Histone superfamily proteir | -2.699137607 | 8.44E-06 | 0.000157544 |
| AT3G06030 | NPK1-related protein kinase | -3.320986083 | 8.46E-06 | 0.000157701 |
| AT1G33930 | P-loop containing nucleosid | -4.793315331 | 8.56E-06 | 0.00015911 |
| AT2G32590 | condensin complex subunit; | -2.795958214 | 8.58E-06 | 0.000159222 |
| AT4G24780 | Encodes a pectate lyase inv | -2.53958879 | 8.73E-06 | 0.000161682 |

| | | | | |
|---|---|---|---|---|
| AT3G54400 | Eukaryotic aspartyl protease | -3.302774808 | 8.93E-06 | 0.000164742 |
| AT1G18360 | alpha/beta-Hydrolases supe | -2.445129368 | 8.95E-06 | 0.000164823 |
| AT5G40610 | NAD-dependent glycerol-3-| -2.318883789 | 9.16E-06 | 0.000168193 |
| AT4G14150 | Microtubule motor kinesin I | -3.57653202 | 9.17E-06 | 0.000168211 |
| AT4G31620 | Transcriptional factor B3 fa | -3.223450771 | 9.19E-06 | 0.000168398 |
| AT4G18960 | Floral homeotic gene encoc | -3.222496717 | 9.22E-06 | 0.000168868 |
| AT5G05240 | cation-transporting ATPase, | -2.013106059 | 9.26E-06 | 0.000169454 |
| AT4G01730 | DHHC-type zinc finger famil | -3.953106493 | 9.43E-06 | 0.000172154 |
| AT3G57780 | nucleolar-like protein;(sourc | -2.033141392 | 9.46E-06 | 0.000172369 |
| AT1G10780 | F-box/RNI-like superfamily | -3.562292114 | 9.48E-06 | 0.000172533 |
| AT4G38950 | ATP binding microtubule m | -2.256741313 | 9.50E-06 | 0.000172662 |
| AT5G28640 | Encodes a protein with simi | -3.987416871 | 9.54E-06 | 0.000173156 |
| AT5G10400 | Histone superfamily proteir | -2.772332618 | 9.80E-06 | 0.000177091 |
| AT2G33560 | Encodes BUBR1. May have | -3.562324724 | 9.96E-06 | 0.000179576 |
| AT4G18670 | Leucine-rich repeat (LRR) fa | -3.176631461 | 1.01E-05 | 0.000182289 |
| AT1G53180 | hypothetical protein;(source | -3.850229143 | 1.01E-05 | 0.000182295 |
| AT2G22122 | hypothetical protein;(source | -2.062546118 | 1.03E-05 | 0.00018532 |
| AT4G37810 | EPIDERMAL PATTERNING F | -4.071975306 | 1.06E-05 | 0.000189152 |
| AT1G44830 | Encodes a nuclear-localized | -7.202078182 | 1.08E-05 | 0.000193732 |
| AT1G26920 | zinc finger CCHC domain pro | -2.870821278 | 1.11E-05 | 0.000197815 |
| AT5G19090 | Heavy metal transport/detc | -3.217976955 | 1.13E-05 | 0.000200865 |
| AT1G13650 | hypothetical protein;(source | -2.920549831 | 1.16E-05 | 0.000204684 |
| AT3G44450 | hypothetical protein;(source | -2.066899246 | 1.28E-05 | 0.000222762 |
| AT1G73830 | Encodes the brassinosteroid | -4.966189381 | 1.33E-05 | 0.000229835 |
| AT1G09750 | Eukaryotic aspartyl protease | -2.882025135 | 1.33E-05 | 0.000230095 |
| AT1G73620 | Pathogenesis-related thaun | -4.765395666 | 1.35E-05 | 0.00023357 |
| AT1G07790 | Encodes a histone 2B (H2B) | -2.010564488 | 1.41E-05 | 0.000242521 |
| AT4G38660 | Pathogenesis-related thaun | -2.19843384 | 1.44E-05 | 0.000246718 |
| AT3G11520 | Encodes a B-type mitotic cy | -3.431086786 | 1.48E-05 | 0.000251668 |
| AT3G46320 | Histone superfamily proteir | -3.343588686 | 1.50E-05 | 0.000254459 |
| AT3G01710 | TPX2 (targeting protein for | -3.328878538 | 1.53E-05 | 0.000259319 |
| AT5G47920 | transcription elongation fac | -4.174750549 | 1.57E-05 | 0.000265471 |
| AT1G21740 | DUF630 family protein, puta | -2.741743571 | 1.57E-05 | 0.000265847 |
| AT1G69120 | Floral homeotic gene encoc | -7.581412681 | 1.61E-05 | 0.00027139 |
| AT3G10570 | member of CYP77A | -2.89235812 | 1.62E-05 | 0.000272205 |
| AT1G47840 | Encodes a putative hexokina | -2.228368348 | 1.67E-05 | 0.000280515 |
| AT2G33400 | FK506-binding nuclear-like | | -2.729494489 | 1.68E-05 | 0.000281538 |
| AT3G11210 | SGNH hydrolase-type estera | -2.234817602 | 1.73E-05 | 0.000289507 |
| AT5G15430 | Plant calmodulin-binding pr | -4.338252185 | 1.73E-05 | 0.000289507 |
| AT5G23400 | Leucine-rich repeat (LRR) fa | -2.711606335 | 1.74E-05 | 0.000289559 |
| AT5G55250 | Encodes an enzyme which s | -6.718035647 | 1.75E-05 | 0.00029033 |
| AT1G01420 | UDP-glucosyl transferase 72 | -2.304855566 | 1.76E-05 | 0.000291496 |
| AT3G05980 | hypothetical protein;(source | -5.456884657 | 1.75E-05 | 0.000291496 |
| AT2G20750 | member of BETA-EXPANSIN | -4.186019606 | 1.85E-05 | 0.000305119 |
| AT3G55110 | ABC-2 type transporter fam | -2.005115259 | 1.85E-05 | 0.00030615 |
| AT2G19910 | RNA-dependent RNA polym | -4.873836022 | 1.88E-05 | 0.000310023 |

| | | | | |
|---|---|---|---|---|
| AT4G10340 | photosystem II encoding the | -2.081857622 | 1.92E-05 | 0.000315884 |
| AT3G08940 | Lhcb4.2 protein (Lhcb4.2, pr | -2.901314626 | 1.98E-05 | 0.000323643 |
| AT5G44260 | Encodes a Tandem CCCH Zi | -5.052971809 | 1.99E-05 | 0.000324504 |
| AT1G71020 | Encodes a nuclear localized | -2.481139264 | 2.02E-05 | 0.000329636 |
| AT3G22790 | Encodes a member of the N | -2.739994267 | 2.06E-05 | 0.000334552 |
| AT4G37750 | ANT is required for control | -3.408976109 | 2.12E-05 | 0.00034377 |
| AT1G14430 | glyoxal oxidase-related prot | -3.107959648 | 2.14E-05 | 0.000346867 |
| AT3G60550 | cyclin p3;(source:Araport11 | -6.296359334 | 2.17E-05 | 0.000350935 |
| AT5G59870 | Encodes HTA6, a histone H2 | -2.993967978 | 2.19E-05 | 0.000353212 |
| AT1G23000 | Heavy metal transport/detc | -4.254902857 | 2.20E-05 | 0.000354961 |
| AT1G15820 | Lhcb6 protein (Lhcb6), light | -2.638246753 | 2.22E-05 | 0.000356607 |
| AT1G72250 | Di-glucose binding protein v | -2.549425468 | 2.28E-05 | 0.000366058 |
| AT1G52240 | Encodes a member of KPP-l | -2.984120969 | 2.31E-05 | 0.000369998 |
| AT3G20460 | Major facilitator superfami | -5.966959801 | 2.35E-05 | 0.000374319 |
| AT5G22310 | trichohyalin-like protein;(so | -3.145295641 | 2.36E-05 | 0.000375254 |
| AT2G20870 | cell wall protein precursor;( | -7.45726811 | 2.37E-05 | 0.0003775 |
| AT1G75580 | SAUR-like auxin-responsive | -4.249497479 | 2.38E-05 | 0.000377981 |
| AT3G53380 | Concanavalin A-like lectin p | -3.027054289 | 2.39E-05 | 0.000378689 |
| AT4G24710 | Encodes an AAA+ ATPase th | -2.384787336 | 2.49E-05 | 0.000391754 |
| AT5G55620 | hypothetical protein;(source | -2.57867037 | 2.52E-05 | 0.000395706 |
| AT3G44050 | P-loop containing nucleosid | -3.603168672 | 2.55E-05 | 0.000398771 |
| AT3G16380 | polyadenylate-binding prote | -2.258613573 | 2.57E-05 | 0.000401872 |
| AT2G24230 | Leucine-rich repeat protein | -2.372978046 | 2.59E-05 | 0.000404606 |
| AT1G06360 | Fatty acid desaturase famil | -2.571839139 | 2.60E-05 | 0.000404812 |
| AT3G07540 | Actin-binding FH2 (formin h | -2.178932332 | 2.63E-05 | 0.000408545 |
| AT3G46370 | Leucine-rich repeat protein | -3.048740138 | 2.68E-05 | 0.000416613 |
| AT5G02540 | NAD(P)-binding Rossmann- | -2.118681252 | 2.71E-05 | 0.000420109 |
| AT1G01190 | member of CYP78A | -3.818584454 | 2.75E-05 | 0.0004248 |
| AT1G11730 | Galactosyltransferase famil | -2.548441801 | 2.79E-05 | 0.000428665 |
| AT2G26760 | Cyclin B1;(source:Araport11 | -3.471595142 | 2.79E-05 | 0.000428665 |
| AT5G38970 | Encodes a polypeptide invol | -7.323679172 | 2.81E-05 | 0.000431127 |
| AT1G73600 | Encodes a S-adenosyl-L-me | -2.473559063 | 2.84E-05 | 0.000434823 |
| AT3G51280 | Tetratricopeptide repeat (T | -4.48011179 | 2.86E-05 | 0.000436615 |
| AT5G44110 | Encodes a member of the N | -2.05443676 | 2.92E-05 | 0.000444264 |
| AT1G29930 | Subunit of light-harvesting | -2.456354425 | 2.94E-05 | 0.00044768 |
| AT3G28420 | Putative membrane lipopro | -4.712293525 | 2.95E-05 | 0.000448328 |
| AT5G67460 | O-Glycosyl hydrolases famil | -3.082334092 | 3.07E-05 | 0.000461484 |
| AT4G15960 | alpha/beta-Hydrolases supe | -2.251279091 | 3.08E-05 | 0.000463508 |
| AT4G23290 | Encodes a cysteine-rich rece | -2.643453939 | 3.15E-05 | 0.000471071 |
| AT5G61280 | Remorin family protein;(sou | -4.380247274 | 3.15E-05 | 0.000471489 |
| AT4G28430 | Reticulon family protein;(sc | -4.023376756 | 3.20E-05 | 0.000476935 |
| AT4G12540 | hypothetical protein;(source | -3.680026289 | 3.25E-05 | 0.000481519 |
| AT1G34245 | Encodes a secretory peptide | -3.626896235 | 3.31E-05 | 0.000488383 |
| AT2G01913 | hypothetical protein;(source | -2.348516205 | 3.30E-05 | 0.000488383 |
| AT1G02970 | Protein kinase that negative | -2.677353892 | 3.33E-05 | 0.000491307 |
| AT1G19320 | Pathogenesis-related thaun | -4.873461756 | 3.36E-05 | 0.000493712 |

| | | | | |
|---|---|---|---|---|
| AT1G62510 | Bifunctional inhibitor/lipid-· | -3.736526814 | 3.36E-05 | 0.000493712 |
| AT3G17130 | Plant invertase/pectin meth | -2.41955775 | 3.36E-05 | 0.000493712 |
| AT4G39790 | bZIP transcription factor, pu | -3.57234643 | 3.36E-05 | 0.000493907 |
| AT4G09060 | hypothetical protein;(source | -2.934374008 | 3.37E-05 | 0.000494985 |
| AT1G63650 | Mutant has reduced trichon | -7.030241952 | 3.40E-05 | 0.000498616 |
| AT3G29280 | hypothetical protein;(source | -2.128518323 | 3.44E-05 | 0.000502905 |
| AT2G24210 | terpene synthase 10;(source | -6.017999516 | 3.50E-05 | 0.000509122 |
| AT1G06350 | Fatty acid desaturase famil | -6.11594558 | 3.63E-05 | 0.00052628 |
| AT2G45970 | Encodes a member of the C | -2.27299518 | 3.66E-05 | 0.000529127 |
| AT3G54890 | Encodes a component of th | -2.44013244 | 3.76E-05 | 0.000541585 |
| AT1G80050 | Encodes an adenosine phos | -2.246434616 | 3.82E-05 | 0.000548631 |
| AT3G44960 | shugoshin;(source:Araport1 | -2.902591946 | 3.82E-05 | 0.00054878 |
| AT4G29270 | HAD superfamily, subfamily | -6.142654711 | 3.88E-05 | 0.000555111 |
| AT1G77960 | repressor ROX1-like protein | -2.784606835 | 3.88E-05 | 0.000555163 |
| AT5G38005 | other_RNA;(source:Araport | -3.482807584 | 3.98E-05 | 0.000565634 |
| AT3G27690 | Encodes Lhcb2.4. Belongs t | -2.736252011 | 4.01E-05 | 0.000568294 |
| AT5G08030 | Encodes a member of the g | -4.711214893 | 4.00E-05 | 0.000568294 |
| AT3G56940 | Encodes a putative ZIP prot | -2.283843445 | 4.07E-05 | 0.000574715 |
| AT2G29890 | Encodes a ubiquitously expr | -2.238880993 | 4.08E-05 | 0.000575489 |
| AT3G51590 | Encodes a member of the li | -6.185356018 | 4.11E-05 | 0.000578774 |
| AT3G22142 | Encodes a Protease inhibito | -4.716356034 | 4.19E-05 | 0.000588727 |
| AT2G20590 | Reticulon family protein;(sc | -3.458256217 | 4.22E-05 | 0.000592707 |
| AT5G03870 | Glutaredoxin family protein | -5.913208911 | 4.29E-05 | 0.000600085 |
| AT4G23720 | transmembrane protein, pu | -4.935246931 | 4.29E-05 | 0.00060035 |
| AT4G23820 | Pectin lyase-like superfamil | -2.283288439 | 4.41E-05 | 0.000615466 |
| AT2G37380 | Encodes a member of the N | -5.811717174 | 4.47E-05 | 0.000622061 |
| AT5G43890 | Encodes a YUCCA-like putat | -2.787276143 | 4.47E-05 | 0.000622061 |
| AT4G17860 | carboxyl-terminal proteinas | -5.131864806 | 4.49E-05 | 0.00062355 |
| AT5G24580 | Heavy metal transport/detc | -2.86271335 | 4.73E-05 | 0.000652468 |
| AT1G20930 | Cyclin-dependent kinase, ex | -2.419151167 | 4.75E-05 | 0.000653524 |
| AT2G45490 | Encodes a member of a fan | -2.903079951 | 4.75E-05 | 0.000653601 |
| AT1G52700 | alpha/beta-Hydrolases supe | -2.524230549 | 4.78E-05 | 0.000656949 |
| AT1G28100 | hypothetical protein;(source | -2.480966229 | 4.80E-05 | 0.000658626 |
| AT2G31160 | LIGHT-DEPENDENT SHORT | -3.637127366 | 4.93E-05 | 0.000672618 |
| AT1G79420 | C-type mannose receptor (D | -2.634059075 | 4.97E-05 | 0.000675413 |
| AT3G57040 | response regulator ARR9, A | -2.412685496 | 4.99E-05 | 0.00067763 |
| AT2G31270 | Encodes a cyclin-dependent | -2.272281923 | 5.03E-05 | 0.000682043 |
| AT1G01200 | RAB GTPase homolog A3;(s | -3.311628054 | 5.06E-05 | 0.000685013 |
| AT5G51670 | hypothetical protein (DUF66 | -3.62808282 | 5.12E-05 | 0.000690665 |
| AT4G34790 | SAUR-like auxin-responsive | -3.742693528 | 5.17E-05 | 0.00069595 |
| AT3G19270 | Encodes a protein with ABA | -3.254989418 | 5.22E-05 | 0.000701024 |
| AT4G00480 | MYC-related protein with a | -3.261388098 | 5.22E-05 | 0.000701024 |
| AT4G30250 | P-loop containing nucleosid | -4.089865465 | 5.24E-05 | 0.00070274 |
| AT1G31320 | LOB domain-containing pro | -3.637345861 | 5.47E-05 | 0.000731513 |
| AT1G15175 | Potential natural antisense | -2.167337093 | 5.53E-05 | 0.000737823 |
| AT5G51850 | hypothetical protein;(source | -4.581669858 | 5.72E-05 | 0.000759077 |

| | | | | |
|---|---|---|---|---|
| AT2G26330 | Homologous to receptor pro | -2.054087698 | 5.74E-05 | 0.000760854 |
| AT1G62520 | sulfated surface-like glycop | -2.372214704 | 5.77E-05 | 0.000763723 |
| AT2G22810 | key regulatory enzyme in th | -2.061282985 | 5.78E-05 | 0.000764009 |
| AT4G25420 | Encodes gibberellin 20-oxid | -3.420304224 | 5.82E-05 | 0.000768814 |
| AT4G27440 | light-dependent NADPH:pro | -3.039216861 | 5.88E-05 | 0.0007759 |
| AT3G60840 | Encodes MAP65-4, a non-m | -2.8736366 | 5.91E-05 | 0.000778614 |
| AT5G35740 | Carbohydrate-binding X8 do | -2.540096494 | 5.95E-05 | 0.000782622 |
| AT4G14650 | hypothetical protein;(source | -4.770247558 | 6.00E-05 | 0.000788069 |
| AT1G03010 | Phototropic-responsive NPH | -2.658917164 | 6.06E-05 | 0.000794885 |
| AT5G39024 | This gene encodes a small p | -2.184857941 | 6.07E-05 | 0.000795072 |
| AT5G24105 | Encodes a putative arabinog | -2.256907976 | 6.08E-05 | 0.000796295 |
| AT3G05400 | Major facilitator superfami | -2.419319686 | 6.14E-05 | 0.000802768 |
| AT3G12710 | DNA glycosylase superfami | -2.697327341 | 6.20E-05 | 0.000810291 |
| AT5G33300 | chromosome-associated kir | -2.956358171 | 6.24E-05 | 0.000814913 |
| AT1G04520 | Encodes a plasmodesmal pr | -2.277641951 | 6.26E-05 | 0.000816391 |
| AT1G54790 | GDSL-motif esterase/acyltr | -2.506241216 | 6.31E-05 | 0.000821393 |
| AT4G24050 | NAD(P)-binding Rossmann- | -2.362322819 | 6.51E-05 | 0.000844153 |
| AT5G27550 | P-loop containing nucleosid | -2.300725428 | 6.62E-05 | 0.000855291 |
| AT5G07800 | Flavin-binding monooxygen | -2.277357392 | 6.70E-05 | 0.000861081 |
| AT3G51220 | WEB family protein (DUF82 | -3.373026833 | 6.72E-05 | 0.000863471 |
| AT1G19620 | transmembrane protein;(so | -3.531535278 | 6.79E-05 | 0.00087099 |
| AT1G76410 | RING/U-box superfamily pr | -2.658363392 | 6.80E-05 | 0.00087099 |
| AT3G02120 | hydroxyproline-rich glycopro | -3.943968324 | 6.86E-05 | 0.000876646 |
| AT5G54630 | zinc finger protein-like prot | -2.822018572 | 6.90E-05 | 0.000880741 |
| AT1G62480 | Vacuolar calcium-binding pr | -2.074097085 | 6.97E-05 | 0.000888397 |
| AT4G39510 | member of CYP96A | -2.5145006 | 7.08E-05 | 0.000902443 |
| AT2G20670 | sugar phosphate exchanger | -2.965959039 | 7.15E-05 | 0.000910375 |
| AT1G67180 | zinc finger (C3HC4-type RIN | -2.712169087 | 7.19E-05 | 0.00091398 |
| AT1G25440 | B-box type zinc finger prote | -2.355314441 | 7.20E-05 | 0.000914531 |
| AT4G23496 | Belongs to a six-member ge | -6.456213236 | 7.25E-05 | 0.000919072 |
| AT5G60930 | P-loop containing nucleosid | -2.657970152 | 7.38E-05 | 0.000933314 |
| AT5G37010 | rho GTPase-activating prote | -3.154976017 | 7.46E-05 | 0.000943239 |
| AT4G14380 | cotton fiber protein;(source | -2.377427157 | 7.65E-05 | 0.000961163 |
| AT1G03130 | Encodes a protein predictec | -2.274240486 | 7.80E-05 | 0.000975429 |
| AT4G17680 | SBP (S-ribonuclease binding | -3.126601676 | 7.81E-05 | 0.00097657 |
| AT3G06100 | Encodes NIP7;1, an anther-s | -3.784708314 | 7.90E-05 | 0.000986759 |
| AT1G12860 | Encodes ICE2 (Inducer of CB | -2.455433828 | 7.93E-05 | 0.000988631 |
| AT4G25470 | Encodes a member of the D | -3.603652006 | 7.95E-05 | 0.000989857 |
| AT3G14760 | transmembrane protein;(so | -4.00138392 | 8.04E-05 | 0.000998202 |
| AT5G15510 | TPX2 (targeting protein for | -2.952125985 | 8.21E-05 | 0.00101848 |
| AT2G27740 | RAB6-interacting golgin (DU | -2.29361379 | 8.46E-05 | 0.001042487 |
| AT2G05540 | Glycine-rich protein family;( | -2.476300876 | 8.78E-05 | 0.001073281 |
| AT5G24470 | Encodes a pseudo-response | -2.267801623 | 8.79E-05 | 0.001073281 |
| AT2G28950 | Encodes an expansin. Nami | -2.277113822 | 8.93E-05 | 0.001089351 |
| AT4G28950 | A member of ROP GTPase g | -3.446510123 | 9.00E-05 | 0.001096107 |
| AT5G62920 | Encodes a Type-A response | -2.283733227 | 9.08E-05 | 0.00110473 |

| | | | | |
|---|---|---|---|---|
| AT5G53210 | N/AN/AN/AN/AN/AN/A | -5.591512434 | 9.18E-05 | 0.001114045 |
| AT3G14240 | Subtilase family protein;(so | -2.116041932 | 9.28E-05 | 0.001125056 |
| AT5G65350 | histone 3 11;(source:Arapor | -2.985330059 | 9.41E-05 | 0.001135955 |
| AT1G10850 | Leucine-rich repeat protein | -2.322041482 | 9.51E-05 | 0.001146453 |
| AT3G02310 | MADS-box protein, binds K | -3.399362237 | 9.51E-05 | 0.001146453 |
| AT5G27895 | transposable_element_gen | -3.446035541 | 9.66E-05 | 0.001160539 |
| AT1G02810 | Plant invertase/pectin metl | -2.380358533 | 9.84E-05 | 0.001177889 |
| AT4G24140 | alpha/beta-Hydrolases supe | -3.177371084 | 9.93E-05 | 0.001186757 |
| AT3G16490 | IQ-domain 26;(source:Arapo | -4.052629209 | 0.00010004 | 0.001193399 |
| AT5G20710 | beta-galactosidase 7;(sourc | -4.178794198 | 0.000100714 | 0.001199483 |
| AT5G40942 | Annotated as pseudogene c | -2.891007636 | 0.000102319 | 0.001215299 |
| AT3G28310 | hypothetical protein (DUF67 | -3.078564341 | 0.000102533 | 0.001216525 |
| AT4G27730 | oligopeptide transporter | -2.17054185 | 0.000102644 | 0.001217181 |
| AT5G04530 | Encodes KCS19, a member | -3.410704695 | 0.000104309 | 0.001233593 |
| AT1G67040 | DnaA initiator-associating p | -2.2634248 | 0.000105341 | 0.001243748 |
| AT4G30710 | QWRF motif protein (DUF5 | -2.542967534 | 0.000105849 | 0.001247095 |
| AT5G46690 | beta HLH protein 71;(source | -3.198524076 | 0.000107137 | 0.001259669 |
| AT1G53680 | Encodes glutathione transfe | -2.542066927 | 0.00010867 | 0.001273485 |
| AT5G62280 | DUF1442 family protein (DU | -3.531387729 | 0.000109193 | 0.001278043 |
| AT1G27460 | encodes a calmodulin-bindi | -2.10104613 | 0.000112027 | 0.001306539 |
| AT1G48260 | Encodes a member of the S | -2.697859705 | 0.000112784 | 0.001313972 |
| AT3G06840 | hypothetical protein;(source | -4.39312073 | 0.000114385 | 0.001330505 |
| AT4G39795 | hypothetical protein (DUF58 | -2.655231905 | 0.000116252 | 0.001346507 |
| AT3G16420 | The PBP1(PYK10-binding pr | -2.272387532 | 0.000117435 | 0.001355964 |
| AT1G35530 | Encodes FANCM, a highly c | -2.032489311 | 0.000119177 | 0.001371284 |
| AT1G77690 | Encodes an auxin influx carr | -2.1073058 | 0.000119073 | 0.001371284 |
| AT5G16000 | NSP-interacting kinase (NIK | -2.439279069 | 0.000119204 | 0.001371284 |
| AT1G18810 | phytochrome kinase substra | -2.536176206 | 0.000121351 | 0.001390148 |
| AT5G38300 | homeobox Hox-B3-like prot | -2.937648444 | 0.000121556 | 0.00139177 |
| AT5G57780 | Encodes a atypical member | -3.344408812 | 0.000122566 | 0.001401132 |
| AT3G21330 | basic helix-loop-helix (bHLH | -3.968855923 | 0.000123295 | 0.00140654 |
| AT3G32980 | Peroxidase superfamily pro | -2.925364244 | 0.000125995 | 0.001434618 |
| AT3G60530 | Encodes a member of the G | -2.179986446 | 0.000126018 | 0.001434618 |
| AT5G65360 | Histone superfamily protein | -2.33785059 | 0.000127579 | 0.001447881 |
| AT5G05510 | Mad3/BUB1 homology regi | -3.765100989 | 0.000131359 | 0.001481729 |
| AT1G26600 | Member of a large family o | -2.46752922 | 0.000132022 | 0.001485988 |
| AT5G03130 | hypothetical protein;(source | -5.430411468 | 0.000135003 | 0.001516432 |
| AT5G22430 | Pollen Ole e 1 allergen and | -4.606710937 | 0.000136555 | 0.00152994 |
| AT5G19060 | cytochrome P450 family pro | -2.304616233 | 0.000137939 | 0.001539164 |
| AT2G15090 | Encodes KCS8, a member o | -2.087346252 | 0.000138798 | 0.001544813 |
| AT4G10630 | Glutaredoxin family protein | -4.097047283 | 0.000140523 | 0.001557693 |
| AT4G01460 | basic helix-loop-helix (bHLH | -2.889687258 | 0.000141404 | 0.001566082 |
| AT5G19170 | NEP-interacting protein, pu | -4.526290008 | 0.000141605 | 0.001566519 |
| AT1G05470 | Encodes an inositol polypho | -2.036997779 | 0.000142633 | 0.001573124 |
| AT3G43960 | Encodes a putative cysteine | -2.431689912 | 0.000142574 | 0.001573124 |
| AT3G54340 | Floral homeotic gene encod | -6.323833386 | 0.000142727 | 0.001573161 |

| | | | | |
|---|---|---|---|---|
| AT1G21810 | Encodes a protein that local | -2.856581787 | 0.000144972 | 0.001594106 |
| AT4G13410 | encodes a gene similar to c | -2.036185724 | 0.000145104 | 0.001594764 |
| AT4G33260 | Encodes a CDC20 protein th | -3.239984569 | 0.000145313 | 0.001595262 |
| AT5G48490 | Encodes a protein with simi | -8.973181138 | 0.000145256 | 0.001595262 |
| AT3G23450 | transmembrane protein;(so | -2.492711735 | 0.000148425 | 0.001619088 |
| AT5G54510 | Encodes an IAA-amido synt | -2.398458461 | 0.000149166 | 0.001624748 |
| AT1G23340 | carboxyl-terminal proteinas | -3.882826906 | 0.000152837 | 0.001654223 |
| AT1G17030 | hypothetical protein;(source | -2.568099452 | 0.000153879 | 0.001661228 |
| AT5G14070 | Encodes glutaredoxin ROXY | -3.552115894 | 0.000155411 | 0.001674466 |
| AT1G13635 | DNA glycosylase superfami | -6.045164866 | 0.00015687 | 0.001686045 |
| AT1G20720 | RAD3-like DNA-binding heli | -2.253389156 | 0.000157253 | 0.001689338 |
| AT3G51470 | Protein phosphatase 2C fan | -2.054092786 | 0.000157781 | 0.001693475 |
| AT3G60220 | Encodes a putative RING-H2 | -3.028111001 | 0.000159975 | 0.001713936 |
| AT2G37280 | Encodes an ATP-binding cas | -4.382629781 | 0.000160145 | 0.001713984 |
| AT3G25100 | Required for normal meiosi | -2.440254577 | 0.000161502 | 0.001727357 |
| AT4G14310 | Transducin/WD40 repeat-lil | -2.596228558 | 0.000164827 | 0.001756872 |
| AT3G57830 | Leucine-rich repeat protein | -2.512819545 | 0.000165655 | 0.001762327 |
| AT2G05995 | other_RNA;(source:Araport | -2.208039487 | 0.000166504 | 0.001767043 |
| AT5G16600 | Encodes a putative transcrip | -3.012299751 | 0.000166793 | 0.001768422 |
| AT2G47930 | arabinogalactan protein 26; | -2.428768748 | 0.000168254 | 0.001779906 |
| AT4G37110 | Zinc-finger domain of monc | -2.182512984 | 0.000171327 | 0.001804282 |
| AT3G57010 | Calcium-dependent phosph | -3.620390956 | 0.000173014 | 0.00181849 |
| AT1G34355 | Encodes PS1 (Parallel Spind | -3.251355071 | 0.000176174 | 0.00184734 |
| AT4G25240 | Encodes GPI-anchored SKU5 | -3.282568678 | 0.000176562 | 0.00185052 |
| AT3G50570 | hydroxyproline-rich glycopro | -3.203568318 | 0.000176794 | 0.001852074 |
| AT1G34580 | Major facilitator superfami | -5.148548526 | 0.000179885 | 0.001881755 |
| AT1G69040 | ACT-domain containing pro | -2.071842773 | 0.000182805 | 0.001905027 |
| AT4G26540 | Leucine-rich repeat recepto | -2.796993763 | 0.000183439 | 0.001910729 |
| AT5G54148 | sarcosine dehydrogenase-2 | -2.500179722 | 0.000183905 | 0.001913768 |
| AT3G10310 | P-loop nucleoside triphosph | -2.545717519 | 0.00018444 | 0.001918425 |
| AT2G41990 | late embryogenesis abunda | -3.950124226 | 0.000184715 | 0.001918786 |
| AT3G26330 | putative cytochrome P450 | -5.804791276 | 0.000185872 | 0.001927364 |
| AT3G47500 | Dof-type zinc finger domain | -2.838436209 | 0.000187721 | 0.001943331 |
| AT4G24265 | homeobox protein;(source:A | -2.911160665 | 0.000190053 | 0.001959143 |
| AT3G61610 | Galactose mutarotase-like s | -2.116642902 | 0.000191874 | 0.001973278 |
| AT3G48540 | Cytidine/deoxycytidylate de | -2.347338715 | 0.000200487 | 0.002046503 |
| AT1G14250 | GDA1/CD39 nucleoside pho | -2.809657845 | 0.000202098 | 0.002053558 |
| AT5G64080 | Bifunctional inhibitor/lipid- | -2.287577466 | 0.000206603 | 0.002089477 |
| AT2G37070 | Encodes a microtubule-asso | -2.897186111 | 0.000214709 | 0.002161483 |
| AT5G22880 | Encodes a histone 2B (H2B) | -2.434718311 | 0.000214969 | 0.002163103 |
| AT3G01330 | Member of the E2F transcri | -2.969508354 | 0.000217714 | 0.002186719 |
| AT5G67390 | glycosyltransferase-like pro | -3.280140961 | 0.000220287 | 0.002207497 |
| AT1G64625 | Encodes a plant-specific ba | -4.16627753 | 0.000220438 | 0.00220801 |
| AT3G17350 | wall-associated receptor kin | -2.002231698 | 0.000225749 | 0.002248874 |
| AT3G61820 | Eukaryotic aspartyl protease | -2.048969566 | 0.000226307 | 0.002253411 |
| AT5G07690 | Encodes a putative transcrip | -2.128331186 | 0.00022704 | 0.002258655 |

| | | | | |
|---|---|---|---|---|
| AT5G13630 | Encodes magnesium chelat | -2.175656248 | 0.000231737 | 0.002297539 |
| AT1G20030 | Pathogenesis-related thaun | -2.923432745 | 0.000231978 | 0.002298392 |
| AT1G78260 | RNA-binding (RRM/RBD/RI | -3.638167425 | 0.000233374 | 0.002309093 |
| AT5G52930 | hypothetical protein (DUF2! | -3.452809713 | 0.000234514 | 0.002318282 |
| AT1G07270 | Cell division control, Cdc6;(s | -2.530958378 | 0.000234969 | 0.002321734 |
| AT4G01680 | Encodes a putative transcri| | -2.998162338 | 0.00023765 | 0.002339349 |
| AT3G54560 | Encodes HTA11, a histone H | -2.381891936 | 0.000242774 | 0.002376372 |
| AT1G13670 | hypothetical protein;(source | -5.621848829 | 0.00024347 | 0.002382115 |
| AT5G39220 | alpha/beta-Hydrolases supe | -4.86667856 | 0.000247232 | 0.002412082 |
| AT4G25490 | Transcriptional activator th | -3.30010459 | 0.00025 | 0.002434057 |
| AT5G42330 | hypothetical protein;(source | -2.769966781 | 0.000251329 | 0.002442658 |
| AT5G42230 | serine carboxypeptidase-lik | -5.478715805 | 0.000251478 | 0.002443029 |
| AT1G04425 | other_RNA;(source:Araport | -2.752217756 | 0.00025429 | 0.002463794 |
| AT1G65330 | Type I MADS-box protein, re | -5.694392092 | 0.00025537 | 0.002473163 |
| AT1G18330 | EARLY-PHYTOCHROME-RES | -2.972104726 | 0.000258834 | 0.002502294 |
| AT3G48260 | Encodes a member of the V | -4.137311047 | 0.000259347 | 0.002505044 |
| AT5G27000 | Encodes a kinesin-like prote | -2.18561944 | 0.000260248 | 0.002511526 |
| AT1G04310 | encodes an ethylene recept | -2.430625271 | 0.000260785 | 0.002514499 |
| AT5G42180 | Peroxidase superfamily pro | -5.416299935 | 0.000265936 | 0.002556293 |
| AT2G41340 | NRPE5-like protein of unknc | -3.394805192 | 0.000275765 | 0.002636907 |
| AT3G42800 | AF-like protein;(source:Araμ | -5.317955089 | 0.000276247 | 0.002640363 |
| AT4G28720 | Auxin biosynthetic gene reg | -5.107894089 | 0.000277881 | 0.002654822 |
| AT2G23690 | HTH-type transcriptional re | -2.454255927 | 0.000288791 | 0.002745896 |
| AT1G68800 | Encodes a TCP transcription | -5.342340167 | 0.000293584 | 0.002786635 |
| AT3G53530 | Chloroplast-targeted coppe | -3.159065003 | 0.000305869 | 0.002878318 |
| AT1G24020 | MLP-like protein 423;(sourc | -2.768641807 | 0.000312622 | 0.002928042 |
| AT2G46790 | Pseudo-response regulator | -4.341645466 | 0.000313159 | 0.00293182 |
| AT4G20230 | terpenoid synthase superfa | -3.934984568 | 0.000318818 | 0.002973378 |
| AT5G41820 | RAB geranylgeranyl transfe | -2.169402938 | 0.000320357 | 0.00298519 |
| AT5G48600 | member of SMC subfamily | -2.0152288 | 0.000321391 | 0.002993551 |
| AT2G17620 | Cyclin B2;(source:Araport11 | -3.46402088 | 0.000331227 | 0.003067002 |
| AT2G18060 | Encodes a NAC-domain trar | -2.575682536 | 0.000334786 | 0.00309596 |
| AT5G15580 | Encodes LONGIFOLIA1 (LNC | -2.015755314 | 0.000339824 | 0.003133298 |
| AT1G66050 | Encodes a protein that is sir | -3.672679105 | 0.000342026 | 0.003148839 |
| AT3G12870 | transmembrane protein;(so | -4.388788971 | 0.000350431 | 0.00321353 |
| AT3G49110 | Class III peroxidase Perx33. | -3.108095323 | 0.000351266 | 0.003218491 |
| AT1G71760 | hypothetical protein;(source | -2.962002865 | 0.000354395 | 0.003240394 |
| AT2G44690 | A member of ROP GTPase g | -2.210194003 | 0.000357715 | 0.003266664 |
| AT1G70560 | TAA1 is involved in the shac | -2.80622806 | 0.000359416 | 0.003278097 |
| AT3G26932 | dsRNA-binding protein 3;(s | -2.868583824 | 0.000362048 | 0.003295251 |
| AT1G05440 | C-8 sterol isomerase;(sourc | -4.293886245 | 0.000362216 | 0.003295412 |
| AT1G63820 | CCT motif family protein;(s | -2.350497451 | 0.000364446 | 0.00330691 |
| AT1G68050 | Encodes FKF1, a flavin-bind | -2.11215768 | 0.000364837 | 0.00330691 |
| AT3G49950 | GRAS family transcription f | -5.474216805 | 0.000364406 | 0.00330691 |
| AT4G05520 | Encodes AtEHD2, one of the | -2.41151287 | 0.000374888 | 0.003382636 |
| AT1G21060 | Serine/Threonine-kinase, pu | -2.084191418 | 0.000376588 | 0.003395179 |

| | | | | |
|---|---|---|---|---|
| AT1G51055 | FBD-like domain family pro | -5.923234248 | 0.000379718 | 0.003420586 |
| AT2G17630 | Pyridoxal phosphate (PLP)-c | -2.119481178 | 0.000381024 | 0.003429532 |
| AT3G02500 | mental retardation GTPase | -4.381909517 | 0.000384304 | 0.003453379 |
| AT4G32830 | Encodes a member of a fan | -2.349283282 | 0.000388816 | 0.003484275 |
| AT5G60880 | Encodes BASL (BREAKING ( | -6.75684581 | 0.000389015 | 0.003484275 |
| AT3G22120 | cell wall-plasma membrane | -2.60156741 | 0.000390088 | 0.003491049 |
| AT5G06940 | Leucine-rich repeat recepto | -2.826308304 | 0.000397939 | 0.003545263 |
| AT4G37490 | Cyclin-dependent protein ki | -3.056901692 | 0.000403813 | 0.00358462 |
| AT5G01910 | myelin transcription factor; | -3.834035265 | 0.000403715 | 0.00358462 |
| AT1G50280 | Phototropic-responsive NPH | -2.110787999 | 0.000416649 | 0.003680681 |
| AT5G01120 | hypothetical protein (DUF67 | -5.847958939 | 0.000418348 | 0.003688497 |
| AT1G49910 | Encodes a homolog of the y | -2.940054261 | 0.000420072 | 0.00370048 |
| AT3G28380 | P-glycoprotein 17;(source:A | -2.789432444 | 0.000420044 | 0.00370048 |
| AT1G61450 | CAP-gly domain linker;(sour | -4.034987695 | 0.000421726 | 0.003710576 |
| AT1G75590 | SAUR-like auxin-responsive | -4.130540033 | 0.00043237 | 0.003787512 |
| AT3G59480 | Encodes a member of the fi | -3.201909129 | 0.000435028 | 0.003801684 |
| AT3G18900 | ternary complex factor MIP | -2.154095467 | 0.000438976 | 0.00382673 |
| AT5G63540 | Encodes RMI1. Suppresses | -3.166431714 | 0.000442349 | 0.003849907 |
| AT2G01505 | Member of a large family o | -2.783125372 | 0.000446324 | 0.003878758 |
| AT2G39855 | plant/protein;(source:Arapc | -3.492015007 | 0.00045792 | 0.003970089 |
| AT4G13690 | RNA-binding protein;(sourc | -2.796997806 | 0.000458383 | 0.003970094 |
| AT3G59900 | Encodes ARGOS (Auxin-Reg | -2.049844075 | 0.0004593 | 0.003975761 |
| AT3G53250 | SAUR-like auxin-responsive | -5.783947066 | 0.000460739 | 0.003983503 |
| AT5G12360 | Encodes a protein that prot | -2.142118802 | 0.000462141 | 0.003992474 |
| AT1G10460 | germin-like protein (GLP7) | -2.20681602 | 0.000464245 | 0.004007494 |
| AT5G22500 | Encodes a member of the e | -3.179067773 | 0.000464644 | 0.004009358 |
| AT3G51400 | hypothetical protein (DUF24 | -2.203847243 | 0.000470423 | 0.004046489 |
| AT5G37950 | UDP-Glycosyltransferase su | -6.653005847 | 0.000483583 | 0.004136968 |
| AT3G24515 | ubiquitin-conjugating enzyn | -2.017452048 | 0.000489293 | 0.004172002 |
| AT2G25880 | Encodes a member of a fan | -3.100849463 | 0.00049669 | 0.004219456 |
| AT1G11080 | serine carboxypeptidase-like | -4.010245175 | 0.000505274 | 0.004277465 |
| AT3G22540 | hypothetical protein (DUF16 | -6.357770472 | 0.000510154 | 0.00431545 |
| AT1G53035 | transmembrane protein;(so | -2.522499763 | 0.000513222 | 0.004333043 |
| AT2G27970 | CDK-subunit 2;(source:Arap | -2.186240725 | 0.000515107 | 0.004345606 |
| AT4G21200 | Encodes a protein with gibb | -2.662237405 | 0.000518517 | 0.004367651 |
| AT3G15570 | Phototropic-responsive NPH | -2.340285717 | 0.000524037 | 0.004410761 |
| AT3G59400 | GUN, genomes uncoupled, | -2.468511089 | 0.000524441 | 0.004412468 |
| AT3G21090 | ABC-2 type transporter fam | -4.858473432 | 0.000531849 | 0.00446031 |
| AT4G32890 | Encodes a member of the G | -2.9739076 | 0.000550047 | 0.004592665 |
| AT5G59260 | Concanavalin A-like lectin p | -4.200222949 | 0.000560588 | 0.004664687 |
| AT2G20515 | pollen Ole e I family allerge | -3.162343087 | 0.000574087 | 0.004760756 |
| AT3G19200 | hypothetical protein;(source | -5.053983193 | 0.000574927 | 0.004764113 |
| AT5G05860 | Encodes a cytokinin N-gluco | -2.031336376 | 0.000584121 | 0.004830718 |
| AT1G75940 | encodes a protein similar to | -6.453425845 | 0.000604158 | 0.004963197 |
| AT1G09610 | glucuronoxylan 4-O-methylt | -2.079804011 | 0.000608726 | 0.004985774 |
| AT1G16070 | Member of TLP family | -3.491925808 | 0.000609746 | 0.004992264 |

| | | | | |
|---|---|---|---|---|
| AT2G42870 | Encodes PHYTOCHROME RA | -4.249884811 | 0.000616625 | 0.005041055 |
| AT1G28290 | Encodes an atypical arabinc | -4.096965941 | 0.000631216 | 0.005135747 |
| AT2G42900 | Plant basic secretory protei | -2.681488052 | 0.000637354 | 0.005171957 |
| AT5G45307 | Encodes a microRNA that ta | -2.665340749 | 0.000639511 | 0.005185166 |
| AT3G56100 | Protein kinase expressed in | -3.594199082 | 0.000642462 | 0.005205387 |
| AT1G05065 | Member of a large family o | -2.646093826 | 0.000649194 | 0.005246672 |
| AT1G16330 | core cell cycle genes | -2.655445272 | 0.000649756 | 0.005247346 |
| AT1G18350 | MAP kinase kinase7. Memb | -6.094130347 | 0.000651535 | 0.005257322 |
| AT5G56320 | member of Alpha-Expansin | -3.473626949 | 0.000653117 | 0.005260925 |
| AT2G22140 | Forms a complex with MUS | -2.689010757 | 0.000655839 | 0.005280911 |
| AT5G27890 | hypothetical protein;(source | -2.846478389 | 0.000659132 | 0.005301592 |
| AT5G11550 | ARM repeat superfamily pro | -2.08787554 | 0.000662064 | 0.005315425 |
| AT3G19210 | Encodes RAD54, a member | -2.017156553 | 0.000666609 | 0.005344979 |
| AT4G29140 | Encodes Activated Disease S | -2.236583933 | 0.000678955 | 0.005415503 |
| AT3G56300 | Cysteinyl-tRNA synthetase, | -2.246655659 | 0.000683414 | 0.005446938 |
| AT2G26700 | Encodes PID2, a homolog of | -2.932574644 | 0.000684398 | 0.005450823 |
| AT1G51780 | encodes a member of the s | -3.267055316 | 0.000684763 | 0.005451751 |
| AT4G21970 | senescence regulator (Prote | -3.324525562 | 0.000687823 | 0.005470152 |
| AT4G13495 | other_RNA;(source:Araport | -2.677329684 | 0.000690537 | 0.005489747 |
| AT1G26210 | AtSOFL1 acts redundantly w | -2.895159831 | 0.000699551 | 0.005547325 |
| AT2G38160 | hypothetical protein;(source | -2.163927092 | 0.000707194 | 0.00559579 |
| AT5G02440 | 60S ribosomal protein L36;( | -3.403169812 | 0.000712303 | 0.005626063 |
| AT3G52910 | Growth regulating factor en | -4.785850686 | 0.000713812 | 0.00563528 |
| AT2G35075 | hypothetical protein;(source | -4.711266488 | 0.000715613 | 0.005644073 |
| AT1G58170 | Disease resistance-responsi | -2.383153424 | 0.000718324 | 0.005661442 |
| AT1G22220 | F-box family protein;(source | -2.771689251 | 0.000718722 | 0.005662482 |
| AT1G63520 | hypothetical protein (DUF35 | -3.958366373 | 0.000734711 | 0.005776006 |
| AT1G05490 | chromatin remodeling 31;(s | -2.781426295 | 0.000736275 | 0.005784155 |
| AT1G20480 | AMP-dependent synthetase | -2.515543187 | 0.000758542 | 0.005920897 |
| AT4G34250 | Encodes KCS16, a member o | -2.201527947 | 0.000766863 | 0.005971502 |
| AT3G50870 | Encodes a GATA transcriptio | -3.712288729 | 0.000769236 | 0.005982783 |
| AT4G12440 | adenine phosphoribosyl tran | -4.844389053 | 0.000769472 | 0.005982783 |
| AT3G05140 | ROP binding protein kinases | -3.039529755 | 0.000770538 | 0.005987107 |
| AT4G29905 | hypothetical protein;(source | -2.231688047 | 0.000773822 | 0.006005954 |
| AT1G30040 | Encodes a gibberellin 2-oxic | -2.098364307 | 0.000776561 | 0.00601869 |
| AT2G42220 | Rhodanese/Cell cycle contro | -2.177309057 | 0.00078078 | 0.006040713 |
| AT5G24330 | Encodes a SET-domain prot | -3.425110178 | 0.000782627 | 0.006052871 |
| AT3G05415 | transposable_element_gen | -2.591361324 | 0.000789638 | 0.006090259 |
| AT4G28180 | hypothetical protein;(source | -2.183463277 | 0.000789889 | 0.006090259 |
| AT5G26230 | Encodes a member of the M | -2.988520151 | 0.000793654 | 0.006110131 |
| AT1G68190 | B-box zinc finger family pro | -2.090154749 | 0.000795397 | 0.006121399 |
| AT1G74480 | RWP-RK domain-containing | -2.732192053 | 0.000796096 | 0.006124629 |
| AT1G12570 | Ortholog of maize IPE1 gen | -3.558257602 | 0.000806756 | 0.006195773 |
| AT5G46570 | Encodes BR-signaling kinase | -2.107539893 | 0.000812916 | 0.00623435 |
| AT4G36450 | member of MAP Kinase | -2.489025983 | 0.000823698 | 0.006301607 |
| AT1G73590 | Encodes an auxin efflux carr | -2.039476924 | 0.000825588 | 0.006308063 |

| | | | | |
|---|---|---|---|---|
| AT3G14000 | Belongs to five-member BR | -2.183846284 | 0.00082598 | 0.006308063 |
| AT3G25130 | acidic leucine-rich nuclear p | -2.154088759 | 0.00083154 | 0.006339481 |
| AT1G14630 | XRI1-like protein;(source:Ar | -2.298274054 | 0.000833716 | 0.006351652 |
| AT5G47330 | alpha/beta-Hydrolases supe | -2.243058495 | 0.000838507 | 0.006377071 |
| AT1G78600 | light-regulated zinc finger p | -2.275982954 | 0.00083918 | 0.006377769 |
| AT2G41550 | Rho termination factor;(sou | -2.668179779 | 0.00083998 | 0.006381639 |
| AT5G04770 | Encodes a member of the c | -2.185334963 | 0.000844213 | 0.006409354 |
| AT3G28315 | transposable_element_gen | -5.813121098 | 0.000856255 | 0.006479457 |
| AT4G15990 | hypothetical protein;(source | -2.165908055 | 0.000860765 | 0.006503488 |
| AT1G44740 | hypothetical protein;(source | -3.356098581 | 0.000862168 | 0.006509609 |
| AT1G77870 | membrane-anchored ubiqui | -2.913584502 | 0.000873576 | 0.006580771 |
| AT2G26180 | Transient Expression of Pro | -2.455618818 | 0.000882868 | 0.00663488 |
| AT3G23150 | Involved in ethylene percep | -3.405564478 | 0.000882996 | 0.00663488 |
| AT5G51910 | TCP family transcription fac | -2.044818499 | 0.000884174 | 0.006639175 |
| AT5G44550 | Uncharacterized protein far | -2.714957168 | 0.00089095 | 0.006680909 |
| AT4G36850 | PQ-loop repeat family prote | -4.246690084 | 0.000897289 | 0.006723838 |
| AT2G04033 | Encodes a defensin-like (DE | -5.405592349 | 0.000912943 | 0.006810883 |
| AT3G27640 | Transducin/WD40 repeat-li | -2.11927953 | 0.000914561 | 0.006820631 |
| AT1G63710 | Encodes a member of the C | -3.095084896 | 0.000951951 | 0.007035388 |
| AT1G10750 | carboxyl-terminal peptidase | -2.919113493 | 0.000957413 | 0.007070477 |
| AT3G60390 | Encodes homeobox protein | -2.446609641 | 0.000965526 | 0.007120795 |
| AT3G18145 | pseudogene of carbamoyl p | -3.033949918 | 0.000966698 | 0.007125635 |
| AT5G51950 | Glucose-methanol-choline ( | -3.334324295 | 0.000970445 | 0.007149861 |
| AT1G67030 | Encodes a novel C2H2 zinc f | -6.87908615 | 0.001048068 | 0.007657426 |
| AT3G12830 | SAUR-like auxin-responsive | -2.063810562 | 0.001056087 | 0.007710876 |
| AT2G30400 | ovate family protein 2;(sour | -2.410523419 | 0.001069432 | 0.007795331 |
| AT4G16970 | Protein kinase superfamily p | -2.463673627 | 0.001071673 | 0.007806476 |
| AT5G25880 | The malic enzyme (EC 1.1.1 | -4.277503793 | 0.001095583 | 0.007951592 |
| AT5G19340 | hypothetical protein;(source | -5.282802548 | 0.001118399 | 0.008085072 |
| AT5G09970 | member of CYP78A | -6.867956994 | 0.001156643 | 0.008314939 |
| AT2G01275 | RING/FYVE/PHD zinc finger | -2.447883393 | 0.001169903 | 0.008388259 |
| AT1G79170 | transmembrane protein;(so | -5.220054079 | 0.00117721 | 0.008426874 |
| AT3G49650 | P-loop containing nucleosid | -2.307092205 | 0.001178335 | 0.008431491 |
| AT3G04445 | pseudogene of polyribonucl | -2.699505735 | 0.001180836 | 0.00844456 |
| AT3G25190 | The gene encodes nodulin-l | -6.163968908 | 0.001189013 | 0.008486427 |
| AT5G03960 | IQ-domain 12;(source:Arap | -2.268799389 | 0.001200735 | 0.008558943 |
| AT5G18060 | SAUR-like auxin-responsive | -2.14315898 | 0.001209335 | 0.008607322 |
| AT1G04360 | RING/U-box superfamily pr | -3.707445438 | 0.001210163 | 0.008609357 |
| AT5G07000 | Encodes a member of the s | -2.637265912 | 0.001214044 | 0.008634168 |
| AT4G39630 | translation initiation factor; | -2.898989358 | 0.001214687 | 0.008635939 |
| AT5G13790 | AGL15 (AGAMOUS-Like 15) | -2.347175571 | 0.001217443 | 0.008647121 |
| AT3G62740 | beta glucosidase 7;(source: | -5.334950823 | 0.001218854 | 0.008654339 |
| AT1G23410 | Ribosomal protein S27a / U | -2.291047661 | 0.00122102 | 0.008664105 |
| AT4G32780 | phosphoinositide binding pr | -3.585125653 | 0.001280978 | 0.009016607 |
| AT5G55240 | Catalyze hydroperoxide-dep | -4.608589309 | 0.001287498 | 0.009053777 |
| AT2G42170 | Actin family protein;(source | -3.281890838 | 0.001296255 | 0.009100763 |

| | | | | |
|---|---|---|---|---|
| AT1G23080 | Encodes a novel component | -2.445244427 | 0.001307632 | 0.009168895 |
| AT4G13493 | Encodes a microRNA of unk | -3.041784735 | 0.001330143 | 0.009302939 |
| AT3G19350 | Encodes a the C-terminal do | -5.596787704 | 0.001344138 | 0.009382862 |
| AT1G59920 | MADS-box family protein;(s | -3.270705837 | 0.00136537 | 0.009497762 |
| AT3G14740 | RING/FYVE/PHD zinc finger | -2.422907334 | 0.001366761 | 0.009499287 |
| AT2G37720 | Encodes a member of the T | -3.423996973 | 0.001376692 | 0.009543893 |
| AT2G33793 | DNA ligase-like protein;(sou | -3.390153837 | 0.001400614 | 0.00969377 |
| AT5G56540 | Encodes arabinogalactan pr | -2.519926879 | 0.001416599 | 0.009782786 |
| AT5G27330 | Prefoldin chaperone subuni | -2.761898969 | 0.001417724 | 0.009787476 |
| AT2G32179 | Potential natural antisense | -3.285536844 | 0.001419935 | 0.00979657 |
| AT3G14190 | Encodes a 193 amino acid p | -2.85956731 | 0.001430368 | 0.009853049 |
| AT4G00460 | Encodes a member of KPP-l | -2.648586715 | 0.001432624 | 0.009862393 |
| AT3G21050 | transposable_element_gen | -2.109531027 | 0.001449696 | 0.009961149 |
| AT2G30424 | In a tandem repeat with AT | -2.74712654 | 0.001463617 | 0.010034792 |
| AT1G14190 | Glucose-methanol-choline ( | -2.104797232 | 0.001470339 | 0.010065144 |
| AT4G16880 | Leucine-rich repeat (LRR) fa | -2.02546598 | 0.001494438 | 0.010207803 |
| AT2G37950 | RING/FYVE/PHD zinc finger | -2.624494075 | 0.001497436 | 0.010221913 |
| AT2G35700 | encodes a member of the D | -3.261858167 | 0.001518864 | 0.010338626 |
| AT5G18930 | Adenosylmethionine decarb | -2.456488994 | 0.001520131 | 0.01034142 |
| AT4G01580 | AP2/B3-like transcriptional | -3.257603602 | 0.001529416 | 0.010382034 |
| AT1G16390 | organic cation/carnitine tra | -4.106945486 | 0.00155401 | 0.010516424 |
| AT5G43080 | Cyclin A3;(source:Araport11 | -2.41350021 | 0.001581025 | 0.010636871 |
| AT5G49330 | Member of the R2R3 factor | -2.18541942 | 0.001591666 | 0.010701891 |
| AT1G30390 | transposable_element_gen | -3.514733107 | 0.001594507 | 0.010717708 |
| AT1G26945 | Encodes a basic helix-loop-l | -2.156797363 | 0.001606218 | 0.010770813 |
| AT1G46480 | Encodes WOX4, a WUSCHE | -2.825841644 | 0.001606829 | 0.010770813 |
| AT3G42725 | Putative membrane lipopro | -3.235840497 | 0.001634444 | 0.010912548 |
| AT2G24030 | zinc ion binding / nucleic ac | -2.224296515 | 0.0016631 | 0.011063448 |
| AT2G45480 | Growth regulating factor er | -4.500503296 | 0.00170897 | 0.011317085 |
| AT1G48330 | SsrA-binding protein;(sourc | -2.03214462 | 0.001721202 | 0.01137943 |
| AT1G36060 | encodes a member of the D | -3.903043014 | 0.001741406 | 0.011490239 |
| AT1G26780 | Encodes LOF1 (LATERAL OR | -5.985309633 | 0.001751655 | 0.011550911 |
| AT3G10185 | Encodes a Gibberellin-regul | -3.815452775 | 0.001766373 | 0.01163333 |
| AT5G44190 | Encodes GLK2, Golden2-like | -2.335930946 | 0.001767005 | 0.01163333 |
| AT1G62360 | Class I knotted-like homeod | -7.389358766 | 0.001800335 | 0.011808003 |
| AT1G23380 | homeodomain transcription | -2.944121623 | 0.001803992 | 0.011828121 |
| AT4G32280 | Auxin inducible protein. | -3.68509124 | 0.001825851 | 0.011939639 |
| AT3G27920 | Encodes GL1, a Myb-like pro | -6.637173506 | 0.001830846 | 0.011961598 |
| AT1G68875 | hypothetical protein;(source | -6.656602944 | 0.001857296 | 0.012094766 |
| AT1G20870 | Encodes an anti-silencing fa | -3.089099999 | 0.001863627 | 0.012117919 |
| AT1G15050 | Belongs to auxin inducible g | -3.263024419 | 0.00194469 | 0.012540963 |
| AT3G12890 | Encodes a protein belonging | -5.271811083 | 0.001950809 | 0.012569334 |
| AT5G02190 | encodes an aspartic proteas | -3.020682898 | 0.001965251 | 0.012636392 |
| AT2G43445 | F-box and associated intera | -3.019260412 | 0.001990514 | 0.012778725 |
| AT1G67035 | homeobox Hox-B3-like prot | -4.254521715 | 0.002006295 | 0.012851308 |
| AT5G45720 | AAA-type ATPase family pro | -2.080380424 | 0.002005133 | 0.012851308 |

| | | | | |
|---|---|---|---|---|
| AT1G36675 | glycine-rich protein;(source | -3.251371297 | 0.002009928 | 0.012870818 |
| AT2G42260 | Encodes a novel plant-speci | -3.507052286 | 0.002011407 | 0.012874595 |
| AT5G01075 | Encodes a small ER-localize | -2.241442384 | 0.002068971 | 0.013179625 |
| AT1G26540 | Agenet domain-containing | -2.049942451 | 0.002069741 | 0.013180697 |
| AT1G57820 | Encodes a 645-amino acid r | -2.43823677 | 0.002072787 | 0.013196264 |
| AT1G44446 | Encodes chlorophyllide <i>a | -2.270717047 | 0.002107977 | 0.013350522 |
| AT2G26520 | transmembrane protein;(so | -2.980388909 | 0.002111903 | 0.01336766 |
| AT3G14140 | 2-oxoglutarate-dependent c | -2.372569806 | 0.002111576 | 0.01336766 |
| AT2G01905 | cyclin J18 (cycJ18) | -4.307013868 | 0.002134731 | 0.013473264 |
| AT5G15310 | Member of the R2R3 factor | -2.137375688 | 0.002164187 | 0.013600451 |
| AT2G37390 | Chloroplast-targeted coppe | -4.310024828 | 0.002167877 | 0.013619737 |
| AT2G34925 | Belongs to a large gene fan | -2.345533837 | 0.002177905 | 0.013670983 |
| AT1G75520 | A member of SHI gene fam | -4.979105987 | 0.002203495 | 0.01378033 |
| AT4G39010 | glycosyl hydrolase 9B18;(so | -3.070642454 | 0.002203256 | 0.01378033 |
| AT4G22620 | SAUR-like auxin-responsive | -3.209997166 | 0.002205648 | 0.013785931 |
| AT2G37420 | ATP binding microtubule m | -2.087807042 | 0.002230723 | 0.013903021 |
| AT5G01420 | Glutaredoxin family protein | -3.364869619 | 0.00225763 | 0.014030836 |
| AT2G47750 | Encodes GH3.9, a member | -2.486589389 | 0.00228327 | 0.014150077 |
| AT5G28145 | transposable_element_gen | -5.012465043 | 0.002299391 | 0.014225858 |
| AT5G48170 | encodes an F-box protein w | -2.24076548 | 0.002354717 | 0.014514822 |
| AT2G28560 | Encodes a protein of the RA | -2.766520673 | 0.002378423 | 0.014624032 |
| AT1G79060 | TPRXL;(source:Araport11) | -3.043498717 | 0.002388167 | 0.014671603 |
| AT5G03995 | hypothetical protein;(source | -2.069205984 | 0.002396917 | 0.014712989 |
| AT2G42110 | hypothetical protein;(source | -4.370356054 | 0.002426955 | 0.014863082 |
| AT2G44190 | Encodes a novel microtubul | -3.385562918 | 0.00242747 | 0.014863082 |
| AT2G47360 | transmembrane protein;(so | -4.930864701 | 0.002438164 | 0.014919072 |
| AT2G29170 | NAD(P)-binding Rossmann- | -2.320046985 | 0.002456377 | 0.014996622 |
| AT1G69490 | Encodes a member of the N | -3.008303309 | 0.002460387 | 0.015010143 |
| AT5G25490 | Ran BP2/NZF zinc finger-lik | -2.874183254 | 0.002474279 | 0.01505301 |
| AT2G45180 | Bifunctional inhibitor/lipid- | -2.012065792 | 0.002487329 | 0.015105652 |
| AT1G65385 | pseudogene of serpin 3;(sou | -2.715366714 | 0.002502828 | 0.015167776 |
| AT4G14130 | xyloglucan endotransglycos | -3.663349582 | 0.002516406 | 0.015241637 |
| AT1G09170 | P-loop nucleoside triphosph | -2.278215807 | 0.002545156 | 0.015381776 |
| AT5G56570 | Leucine-rich repeat (LRR) fa | -4.402376291 | 0.002573713 | 0.01552014 |
| AT4G27310 | B-box type zinc finger famil | -2.85633109 | 0.002604025 | 0.015672731 |
| AT2G22920 | serine carboxypeptidase-lik | -2.766813155 | 0.002689804 | 0.016074309 |
| AT3G26760 | NAD(P)-binding Rossmann- | -5.726927565 | 0.002708795 | 0.01615258 |
| AT5G01790 | hypothetical protein;(source | -2.123052562 | 0.002726792 | 0.016208465 |
| AT4G13575 | hypothetical protein;(source | -2.052064161 | 0.002758356 | 0.016359131 |
| AT1G02630 | Nucleoside transporter fam | -6.440842897 | 0.002762011 | 0.016376382 |
| AT2G33205 | Serinc-domain containing s | -3.645543825 | 0.002768595 | 0.016410982 |
| AT5G57240 | OSBP(oxysterol binding pro | -2.431473319 | 0.002812649 | 0.016627161 |
| AT4G10310 | encodes a sodium transport | -2.185603513 | 0.00282334 | 0.01664996 |
| AT5G10890 | myosin heavy chain-like pro | -5.782286522 | 0.002837407 | 0.016710444 |
| AT2G37925 | encodes a member of copp | -3.422295982 | 0.002843655 | 0.016738252 |
| AT1G32780 | GroES-like zinc-binding deh | -2.686287681 | 0.002850937 | 0.01677211 |

| | | | | |
|---|---|---|---|---|
| AT3G30460 | RING/U-box superfamily pr | -2.371449474 | 0.002884181 | 0.016922276 |
| AT5G08020 | Encodes a homolog of Repli | -2.148610926 | 0.002911406 | 0.017036425 |
| AT5G38940 | RmlC-like cupins superfami | -5.473979345 | 0.0029123 | 0.017037108 |
| AT2G35210 | A member of ARF GAP dom | -2.194619863 | 0.002940286 | 0.017162047 |
| AT1G04650 | holliday junction resolvase;( | -2.010235278 | 0.002944496 | 0.017175052 |
| AT2G28690 | TOX high mobility group bo: | -4.823572005 | 0.002980963 | 0.017348613 |
| AT4G18550 | DSEL is cytosolic acylhydrol: | -2.301852054 | 0.002983171 | 0.017349869 |
| AT2G24692 | BTB/POZ domain protein;(s | -4.827749904 | 0.002989451 | 0.017375525 |
| AT3G61950 | basic helix-loop-helix (bHLH | -2.895157264 | 0.002991691 | 0.017380979 |
| AT2G44450 | beta glucosidase 15;(source | -2.65858612 | 0.003000139 | 0.017411605 |
| AT4G31910 | Encodes an acyltransferase | -3.620666457 | 0.003010897 | 0.017460174 |
| AT5G02810 | PRR7 and PRR9 are partiall | -2.280231499 | 0.003035953 | 0.017582224 |
| AT4G34510 | Encodes KCS17, a member | -4.138790078 | 0.003072865 | 0.017744438 |
| AT5G42053 | This gene encodes a small | -2.624632917 | 0.003187075 | 0.01826923 |
| AT4G31610 | Expressed specifically in rep | -5.198453239 | 0.003187949 | 0.018269467 |
| AT2G26750 | alpha/beta-Hydrolases supe | -5.697223146 | 0.003196408 | 0.018284485 |
| AT4G13235 | Encodes a defensin-like (DE | -4.906594551 | 0.003195644 | 0.018284485 |
| AT2G28680 | RmlC-like cupins superfami | -6.315448192 | 0.00321177 | 0.018353212 |
| AT2G28780 | P-hydroxybenzoic acid efflu: | -5.465426866 | 0.003234093 | 0.018447116 |
| AT1G55010 | Predicted to encode a PR (p | -4.93829208 | 0.003254457 | 0.018529532 |
| AT2G28050 | Encodes a relatively short re | -2.09017716 | 0.003308411 | 0.018758784 |
| AT1G07450 | NAD(P)-binding Rossmann- | -2.448761216 | 0.003333704 | 0.018858304 |
| AT1G53860 | Remorin family protein;(sou | -4.408154294 | 0.00333708 | 0.01886767 |
| AT3G43190 | Encodes a protein with sucr | -2.52511625 | 0.003336472 | 0.01886767 |
| AT4G37890 | Zinc finger (C3HC4-type RIN | -3.436708531 | 0.00334767 | 0.01891779 |
| AT2G15750 | transposable_element_gen | -4.704635166 | 0.003389702 | 0.019110982 |
| AT1G58430 | Encodes an anther-specific | -6.285653775 | 0.003407801 | 0.019183424 |
| AT3G44900 | member of Putative Na+/H | -2.418425819 | 0.003416621 | 0.019218273 |
| AT1G02070 | zinc ion-binding protein;(so | -5.37634974 | 0.003419363 | 0.019228763 |
| AT1G52750 | alpha/beta-Hydrolases supe | -2.928695831 | 0.003432581 | 0.019282273 |
| AT1G62770 | PMEI9 pectin methylesetera | -2.569268429 | 0.003440966 | 0.019310628 |
| AT3G12970 | serine/arginine repetitive m | -3.229871586 | 0.003473205 | 0.019476599 |
| AT5G59360 | hypothetical protein;(source | -2.902858222 | 0.003485418 | 0.019525111 |
| AT2G37300 | transmembrane protein;(so | -3.063290359 | 0.003528759 | 0.019717527 |
| AT5G38000 | Zinc-binding dehydrogenase | -2.666908613 | 0.003544267 | 0.01978905 |
| AT5G57770 | auxin canalization protein ( | -5.586221446 | 0.003587317 | 0.019983615 |
| AT2G30370 | Encodes a small, potentially | -2.939535074 | 0.003629392 | 0.020197472 |
| AT2G45050 | Encodes a member of the G | -2.824079915 | 0.003657575 | 0.020328442 |
| AT3G50310 | Encodes a member of MEKI | -2.79658329 | 0.00366616 | 0.020352124 |
| AT5G49380 | N/AN/AN/AN/AN/AN/A | -2.459183144 | 0.003675691 | 0.020398178 |
| AT2G28040 | Eukaryotic aspartyl protease | -3.170400581 | 0.0036977 | 0.020499561 |
| AT4G08113 | transposable_element_gen | -3.785935208 | 0.003744237 | 0.020679128 |
| AT1G19830 | SAUR-like auxin-responsive | -6.302079281 | 0.00374948 | 0.020697659 |
| AT5G02750 | Encodes an E3 ligase, SHOC | -3.073066569 | 0.003762663 | 0.020765205 |
| AT2G42530 | Encodes COR15B, a protein | -2.151940743 | 0.003767361 | 0.020770219 |
| AT1G71890 | Encodes a sucrose transpor | -2.086573445 | 0.003772837 | 0.020789129 |

| | | | | |
|---|---|---|---|---|
| AT4G01670 | hypothetical protein;(source | -2.665907407 | 0.003773636 | 0.020789129 |
| AT1G03120 | responsive to abscisic acid 2 | -2.303926594 | 0.003777709 | 0.020795896 |
| AT3G14540 | Encodes a sesterterpene sy | -6.250074018 | 0.003783525 | 0.020820456 |
| AT4G28560 | encodes a member of a nov | -2.384803263 | 0.003792493 | 0.020845876 |
| AT2G42290 | Leucine-rich repeat protein | -2.094628825 | 0.003847245 | 0.021088672 |
| AT5G46915 | transcriptional factor B3 fai | -3.24329791 | 0.003888494 | 0.021256327 |
| AT5G07810 | SNF2 domain-containing pr | -2.087877356 | 0.003890257 | 0.021260663 |
| AT2G45900 | Phosphatidylinositol N-acet | -2.349524241 | 0.003912771 | 0.021367724 |
| AT2G37210 | Encodes a protein of unknov | -2.685290051 | 0.003942206 | 0.021498249 |
| AT3G49900 | Phototropic-responsive NPH | -2.076824559 | 0.003964627 | 0.021586396 |
| AT5G56220 | P-loop containing nucleosid | -2.262854285 | 0.004115072 | 0.022224474 |
| AT2G43870 | Pectin lyase-like superfamil | -6.192350059 | 0.004126782 | 0.022275682 |
| AT3G51230 | chalcone-flavanone isomera | -2.897083942 | 0.004142834 | 0.022323856 |
| AT4G01410 | Late embryogenesis abunda | -2.130213427 | 0.004208932 | 0.022569099 |
| AT1G74055 | transmembrane protein;(so | -2.946621791 | 0.004245254 | 0.02271387 |
| AT1G78990 | HXXXD-type acyl-transferas | -2.253955514 | 0.004278112 | 0.022839514 |
| AT3G54710 | Encodes a cyclin-dependent | -2.223904196 | 0.00429569 | 0.022921975 |
| AT1G54840 | Encodes an atypical membe | -3.449826845 | 0.004323758 | 0.02302159 |
| AT1G11850 | transmembrane protein;(so | -3.868539828 | 0.004347971 | 0.02308879 |
| AT5G21150 | AGO9-dependent sRNA sile | -3.688318496 | 0.00436253 | 0.023154885 |
| AT1G36940 | myotubularin-like protein;(s | -2.043558786 | 0.004365488 | 0.02316498 |
| AT2G21200 | SAUR-like auxin-responsive | -2.35578924 | 0.004394723 | 0.02328582 |
| AT1G78090 | homologous to the C-termi | -3.592083209 | 0.004413644 | 0.023341427 |
| AT3G18070 | beta glucosidase 43;(source | -2.738422274 | 0.004413352 | 0.023341427 |
| AT4G11460 | Encodes a cysteine-rich rece | -6.241111968 | 0.004420937 | 0.02336309 |
| AT1G58460 | hypothetical protein;(source | -6.213365718 | 0.004443049 | 0.023447881 |
| AT3G11260 | WOX5 is a member of the V | -4.969145095 | 0.0044801 | 0.023571987 |
| AT5G16350 | O-acyltransferase (WSD1-li | -2.373774123 | 0.004483042 | 0.023571987 |
| AT2G46455 | OxaA/YidC-like membrane i | -2.266259989 | 0.004505512 | 0.023678775 |
| AT1G02800 | Encodes a protein with simi | -3.652911535 | 0.004513156 | 0.023698084 |
| AT5G62170 | LOW protein: M-phase indu | -2.088988062 | 0.004527473 | 0.023754337 |
| AT1G02340 | Encodes a light-inducible, n | -2.431511881 | 0.004625519 | 0.024167499 |
| AT3G43850 | hypothetical protein;(source | -2.912656837 | 0.004638023 | 0.024206926 |
| AT2G01610 | Plant invertase/pectin meth | -2.249324343 | 0.004654282 | 0.024262907 |
| AT4G35810 | 2-oxoglutarate (2OG) and F | -5.410126003 | 0.004702017 | 0.024442903 |
| AT1G74650 | Member of the R2R3 factor | -2.345947426 | 0.004740504 | 0.024577984 |
| AT3G26050 | TPX2 (targeting protein for | -3.189223589 | 0.004740114 | 0.024577984 |
| AT1G01030 | AP2/B3-like transcriptional | -2.837761999 | 0.004751558 | 0.024598618 |
| AT4G27570 | UDP-Glycosyltransferase su | -2.372876828 | 0.004768152 | 0.024647936 |
| AT3G57860 | Encodes a protein that cont | -2.030586605 | 0.004780787 | 0.024693412 |
| AT3G27500 | Cysteine/Histidine-rich C1 d | -3.173089615 | 0.004786632 | 0.024702914 |
| AT3G24450 | Heavy metal transport/detc | -3.682454414 | 0.004790654 | 0.024703639 |
| AT2G12440 | transposable_element_gen | -2.039140037 | 0.004899806 | 0.025124682 |
| AT3G60900 | FASCICLIN-like arabinogala | -2.669084172 | 0.004927862 | 0.025233141 |
| AT4G04408 | pseudogene of histone H2A | -3.164392746 | 0.004939054 | 0.025266845 |
| AT1G23965 | transcription factor;(source | -4.219388989 | 0.004969845 | 0.025388822 |

| | | | | |
|---|---|---|---|---|
| AT3G52550 | transcription repressor OFP | -4.487930516 | 0.004974521 | 0.025406496 |
| AT5G42700 | AP2/B3-like transcriptional | -3.904335922 | 0.005012196 | 0.025563484 |
| AT1G75190 | hypothetical protein;(source | -2.097995357 | 0.005068719 | 0.025779806 |
| AT4G20420 | Tapetum specific protein TA | -2.251711636 | 0.005126239 | 0.026018035 |
| AT2G38720 | microtubule-associated pro | -2.038378907 | 0.005160233 | 0.026184511 |
| AT1G02681 | Pseudogene of AT4G02160 | -3.272447567 | 0.00516749 | 0.02620314 |
| AT3G60890 | binding protein;(source:Ara | -3.905051853 | 0.005196885 | 0.026321762 |
| AT5G01190 | putative laccase, a membe | -2.486808949 | 0.00526135 | 0.026580727 |
| AT1G14240 | GDA1/CD39 nucleoside pho | -2.18615245 | 0.00528853 | 0.026705738 |
| AT3G10190 | Encodes a protein with seq | -2.065255412 | 0.005380689 | 0.02705894 |
| AT5G59990 | CCT motif family protein;(sc | -2.755132527 | 0.005536776 | 0.027691456 |
| AT1G65620 | required for formation of a | -2.340564081 | 0.005604145 | 0.028008479 |
| AT1G66060 | hypothetical protein (DUF57 | -2.730857725 | 0.005696593 | 0.028384975 |
| AT5G25380 | core cell cycle genes | -3.371171819 | 0.005718013 | 0.028468018 |
| AT2G32550 | Cell differentiation, Rcd1-lil | -3.538507712 | 0.005783256 | 0.028734117 |
| AT5G37940 | Zinc-binding dehydrogenase | -10.77604683 | 0.005814305 | 0.028875302 |
| AT1G11112 | hypothetical protein;(source | -3.047884448 | 0.005826877 | 0.028924635 |
| AT2G36400 | Growth regulating factor er | -2.636348563 | 0.005950356 | 0.029405637 |
| AT3G43840 | 3-oxo-5-alpha-steroid 4-deh | -4.538250116 | 0.005999208 | 0.029617504 |
| AT1G68360 | Encodes a nuclear localized | -4.491901418 | 0.006012831 | 0.029639686 |
| AT3G57130 | Encodes BOP1. Contains Pf | -6.05144541 | 0.006117401 | 0.030026568 |
| AT5G63087 | Encodes a Plant thionin fam | -4.149339801 | 0.006167431 | 0.030184619 |
| AT2G23030 | encodes a member of SNF1 | -3.726116769 | 0.00621501 | 0.030386149 |
| AT5G35640 | Putative endonuclease or gl | -2.843459606 | 0.006215538 | 0.030386149 |
| AT4G33560 | Wound-responsive family p | -2.19742011 | 0.006260194 | 0.030542132 |
| AT5G22930 | enabled-like protein (DUF16 | -2.467154539 | 0.006326896 | 0.030759095 |
| AT1G70985 | hydroxyproline-rich glycopro | -2.680222817 | 0.006339858 | 0.030808448 |
| AT3G63210 | encodes a novel zinc-finger | -2.385260872 | 0.006367697 | 0.030909481 |
| AT1G31990 | transmembrane protein;(so | -4.686884753 | 0.006495966 | 0.03139062 |
| AT5G03010 | Galactose oxidase/kelch rep | -4.695669724 | 0.006535822 | 0.031537068 |
| AT4G06746 | encodes a member of the D | -2.054220936 | 0.006564679 | 0.031641492 |
| AT4G34770 | SAUR-like auxin-responsive | -2.58672364 | 0.006579505 | 0.031699019 |
| AT1G11600 | member of CYP77B | -10.59638168 | 0.00668666 | 0.032077707 |
| AT5G35960 | Protein kinase family protei | -2.919988551 | 0.006698722 | 0.032107673 |
| AT3G20898 | hypothetical protein;(source | -2.403364895 | 0.00670829 | 0.032141654 |
| AT5G45850 | hypothetical protein (DUF68 | -3.545572088 | 0.006878447 | 0.032738047 |
| AT5G41710 | transposable_element_gen | -2.287234255 | 0.00693923 | 0.032961345 |
| AT1G11740 | ankyrin repeat family protei | -3.681543379 | 0.007103225 | 0.033638297 |
| AT2G42150 | DNA-binding bromodomain | -2.012578059 | 0.007185171 | 0.033964124 |
| AT4G30380 | Encodes a Plant Natriuretic | -2.528036055 | 0.007194634 | 0.03399042 |
| AT3G30180 | Encodes a cytochrome p450 | -2.469001554 | 0.007210876 | 0.034037821 |
| AT5G41590 | LURP-one-like protein (DUF | -2.83948067 | 0.007366584 | 0.034586663 |
| AT5G37980 | Zinc-binding dehydrogenase | -3.628874734 | 0.007370755 | 0.034598839 |
| AT1G62190 | Kua-ubiquitin conjugating e | -4.946180177 | 0.007413295 | 0.034743997 |
| AT2G35310 | Transcriptional factor B3 fa | -4.302801354 | 0.007454759 | 0.034851395 |
| AT4G18422 | transmembrane protein;(so | -2.217847329 | 0.007506099 | 0.035016751 |

| | | | | |
|---|---|---|---|---|
| AT5G38930 | RmlC-like cupins superfami | -5.307610767 | 0.007520498 | 0.035061543 |
| AT2G16210 | Transcriptional factor B3 fa | -5.878236703 | 0.007606686 | 0.035335637 |
| AT4G16748 | other_RNA;(source:Araport | -4.681441429 | 0.007600921 | 0.035335637 |
| AT5G63090 | Involved in lateral organ de | -5.878236703 | 0.007606686 | 0.035335637 |
| AT2G02850 | Encodes plantacyanin one o | -2.922696689 | 0.007697239 | 0.035643012 |
| AT5G02140 | Pathogenesis-related thaun | -2.695342238 | 0.00777006 | 0.03590439 |
| AT3G19040 | Encodes a protein similar to | -2.51957106 | 0.008033717 | 0.036812391 |
| AT1G27080 | Encodes a protein with low- | -2.483882038 | 0.008052139 | 0.036881385 |
| AT3G43715 | transposable_element_gen | -5.940574376 | 0.008252237 | 0.037601526 |
| AT1G26911 | Pseudogene of AT1G66590; | -3.127520908 | 0.008257585 | 0.037607239 |
| AT4G27530 | hypothetical protein;(source | -2.123954793 | 0.008270479 | 0.037653348 |
| AT3G61840 | auxin response factor, puta | -2.720172079 | 0.0083343 | 0.037880981 |
| AT5G63590 | flavonol synthase 3;(source | -3.848358814 | 0.008401994 | 0.038133332 |
| AT3G20395 | RING/U-box superfamily pr | -4.95177223 | 0.008418601 | 0.038169201 |
| AT3G59440 | Encodes an endomembrane | -4.792433345 | 0.008481722 | 0.03837603 |
| AT3G60700 | hypothetical protein (DUF1: | -4.792433345 | 0.008481722 | 0.03837603 |
| AT5G25970 | Core-2/I-branching beta-1,6 | -5.805255852 | 0.008486442 | 0.038389465 |
| AT4G14103 | F-box/RNI-like superfamily | -2.183661722 | 0.008535079 | 0.038529986 |
| AT1G67780 | Zinc-finger domain of mono | -2.952679757 | 0.008610433 | 0.038782852 |
| AT3G54180 | Arabidopsis homolog of yea | -3.111336359 | 0.008667166 | 0.038965815 |
| AT4G11950 | transmembrane protein, pu | -4.881636314 | 0.008770715 | 0.039318484 |
| AT4G37740 | Growth regulating factor en | -2.02233122 | 0.008850901 | 0.039588915 |
| AT2G35200 | DUF740 family protein;(sou | -5.195310886 | 0.008860909 | 0.039617515 |
| AT1G70880 | Polyketide cyclase/dehydras | -3.397083499 | 0.008903123 | 0.039790025 |
| AT3G54060 | myosin-M heavy protein;(sc | -2.098547104 | 0.008915761 | 0.039830273 |
| AT4G02810 | A member of the FAF famil | -3.599262758 | 0.009035314 | 0.040200545 |
| AT5G45040 | Encodes a Class I cytochrom | -2.093780399 | 0.009118538 | 0.040496874 |
| AT5G19100 | Eukaryotic aspartyl protease | -2.871250383 | 0.00914849 | 0.040597005 |
| AT1G64450 | Glycine-rich protein family; | -2.287133942 | 0.009259469 | 0.040981655 |
| AT1G47760 | AGAMOUS-like 102;(source | -3.935909842 | 0.009350754 | 0.041318953 |
| AT3G23190 | HR-like lesion-inducing prot | -2.678504416 | 0.009360345 | 0.041344668 |
| AT4G25410 | basic helix-loop-helix (bHLH | -2.447934089 | 0.009363864 | 0.041351881 |
| AT1G74490 | Protein kinase superfamily | -2.449644495 | 0.009426096 | 0.041568099 |
| AT3G53010 | carbohydrate esterase, puta | -2.299552291 | 0.009684123 | 0.042441353 |
| AT4G14403 | N/AN/AN/AN/AN/AN/A | -2.506549298 | 0.009728921 | 0.042595111 |
| AT4G30662 | hypothetical protein;(source | -2.351468096 | 0.009750669 | 0.042664773 |
| AT4G32950 | Protein phosphatase 2C fam | -2.649174736 | 0.009778585 | 0.042727233 |
| AT3G06640 | PAS domain-containing pro | -4.232492393 | 0.009783127 | 0.042738561 |
| AT1G70220 | RNA-processing, Lsm doma | -4.201155835 | 0.009824429 | 0.042867746 |
| AT1G75550 | glycine-rich protein;(source | -4.120201302 | 0.00985418 | 0.042971909 |
| AT5G03670 | histone-lysine N-methyltrar | -2.714654968 | 0.009897902 | 0.043119691 |
| AT5G12270 | 2-oxoglutarate (2OG) and F | -5.71609765 | 0.010146732 | 0.044002619 |
| AT1G15460 | Encodes a efflux-type boron | -4.917241525 | 0.01017586 | 0.044080104 |
| AT1G61760 | Late embryogenesis abunda | -2.119888611 | 0.010243594 | 0.044265501 |
| AT1G09460 | Carbohydrate-binding X8 do | -2.473431351 | 0.010336641 | 0.044561775 |
| AT2G24700 | Transcriptional factor B3 fa | -3.217705148 | 0.01043282 | 0.04486174 |

| | | | | |
|---|---|---|---|---|
| AT5G16486 | RNA-directed DNA polymer | -3.990630594 | 0.010544879 | 0.04527257 |
| AT4G34970 | A member of actin polymer | -3.288369512 | 0.010581703 | 0.045386231 |
| AT5G14650 | Pectin lyase-like superfamil | -2.296237884 | 0.010672366 | 0.045694648 |
| AT1G02620 | Ras-related small GTP-bind | -3.044335926 | 0.010690596 | 0.045754833 |
| AT2G46570 | putative laccase, a membe | -2.561929338 | 0.010875354 | 0.046364576 |
| AT5G46105 | pre-tRNA tRNA-Pro (antico | -3.691183106 | 0.010885629 | 0.04639936 |
| AT5G37072 | pseudogene of Ribosomal p | -2.670426229 | 0.010944959 | 0.046615999 |
| AT5G56620 | NAC domain containing pro | -3.167223136 | 0.011053588 | 0.047005618 |
| AT3G51325 | RING/U-box superfamily pr | -2.749728274 | 0.01107403 | 0.047083416 |
| AT5G15150 | homeobox-containing gene | -4.166257518 | 0.011077154 | 0.04708757 |
| AT3G57670 | Encodes a a C2H2/C2HC zin | -2.343450181 | 0.01110202 | 0.047165838 |
| AT3G10290 | Nucleotide-sugar transport | -3.982895307 | 0.011108255 | 0.047183184 |
| AT3G53232 | ROTUNDIFOLIA like 1;(sour | -2.876080418 | 0.011152665 | 0.047353474 |
| AT1G54660 | Possibly not a pseudogene l | -2.272242267 | 0.011206432 | 0.047498983 |
| AT5G26630 | MADS-box transcription fac | -2.686402135 | 0.011277765 | 0.047745948 |
| AT3G13840 | GRAS family transcription f | -5.667951767 | 0.011431149 | 0.048311368 |
| AT1G29110 | Cysteine proteinases superf | -3.5763918 | 0.011474836 | 0.048461678 |
| AT3G05727 | Encodes a defensin-like (DE | -7.350862667 | 0.011561567 | 0.048738896 |
| AT2G23530 | Zinc-finger domain of mono | -3.442668206 | 0.01162094 | 0.048934269 |
| AT2G25150 | HXXXD-type acyl-transferas | -3.698387017 | 0.011790703 | 0.049478078 |
| AT3G06220 | AP2/B3-like transcriptional | -3.464503621 | 0.011825782 | 0.049608405 |
| AT3G61250 | LATE MERISTEM IDENTITY2 | -2.469558349 | 0.011924745 | 0.049919329 |
| AT1G10710 | Computational predictions s | -2.039869555 | 0.011951419 | 0.04999939 |

wild type and *trm5-1* .

| **Type** |
| --- |
| UP |
| UP |
| UP |
| UP |
| UP |
| UP |
| UP |
| UP |
| UP |
| UP |
| UP |
| UP |
| UP |
| UP |
| UP |
| UP |
| UP |
| UP |
| UP |
| UP |
| UP |
| UP |
| UP |
| UP |
| UP |
| UP |
| UP |
| UP |
| UP |
| UP |
| UP |
| UP |
| UP |
| UP |
| UP |
| UP |
| UP |
| UP |
| UP |
| UP |
| UP |
| UP |
| UP |

UP
UP
UP
UP
UP
UP
UP
UP
UP
UP
UP
UP
UP
UP
UP
UP
UP
UP
UP
UP
UP
UP
UP
UP
UP
UP
UP
UP
UP
UP
UP
UP
UP
UP
UP
UP
UP
UP
UP
UP
UP
UP
UP
UP
UP
UP

UP
UP
UP
UP
UP
UP
UP
UP
UP
UP
UP
UP
UP
UP
UP
UP
UP
UP
UP
UP
UP
UP
UP
UP
UP
UP
UP
UP
UP
UP
UP
UP
UP
UP
UP
UP
UP
UP
UP
UP
UP
UP
UP
UP
UP
UP

UP
UP
UP
UP
UP
UP
UP
UP
UP
UP
UP
UP
UP
UP
UP
UP
UP
UP
UP
UP
UP
UP
UP
UP
UP
UP
UP
UP
UP
UP
UP
UP
UP
UP
UP
UP
UP
UP
UP
UP
UP
UP
UP
UP
UP
UP

UP
UP
UP
UP
UP
UP
UP
UP
UP
UP
UP
UP
UP
UP
UP
UP
UP
UP
UP
UP
UP
UP
UP
UP
UP
UP
UP
UP
UP
UP
UP
UP
UP
UP
UP
UP
UP
UP
UP
UP
UP
UP
UP
UP
UP
UP

UP
UP
UP
UP
UP
UP
UP
UP
UP
UP
UP
UP
UP
UP
UP
UP
UP
UP
UP
UP
UP
UP
UP
UP
UP
UP
UP
UP
UP
UP
UP
UP
UP
UP
UP
UP
UP
UP
UP
UP
UP
UP
UP
UP
UP
UP

UP
UP
UP
UP
UP
UP
UP
UP
UP
UP
UP
UP
UP
UP
UP
UP
UP
UP
UP
UP
UP
UP
UP
UP
UP
UP
UP
UP
UP
UP
UP
UP
UP
UP
UP
UP
UP
UP
UP
UP
UP
UP
UP
UP
UP
UP

UP
UP
UP
UP
UP
UP
UP
UP
UP
UP
UP
UP
UP
UP
UP
UP
UP
UP
UP
UP
UP
UP
UP
UP
UP
UP
UP
UP
UP
UP
UP
UP
UP
UP
UP
UP
UP
UP
UP
UP
UP
UP
UP
UP
UP
UP

UP
UP
UP
UP
UP
UP
UP
UP
UP
UP
UP
UP
UP
UP
UP
UP
UP
UP
UP
UP
UP
UP
UP
UP
UP
UP
UP
UP
UP
UP
UP
UP
UP
UP
UP
UP
UP
UP
UP
UP
UP
UP
UP
UP
UP
UP

UP
UP
UP
UP
UP
UP
UP
UP
UP
UP
UP
UP
UP
UP
UP
UP
UP
UP
UP
UP
UP
UP
UP
UP
UP
UP
UP
UP
UP
UP
UP
UP
UP
UP
UP
UP
UP
UP
UP
UP
UP
UP
UP
UP
UP
UP

UP
UP
UP
UP
UP
UP
UP
UP
UP
UP
UP
UP
UP
UP
UP
UP
UP
UP
UP
UP
UP
UP
UP
UP
UP
UP
UP
UP
UP
UP
UP
UP
UP
UP
UP
UP
UP
UP
UP
UP
UP
UP
UP
UP
UP
UP

UP
UP
UP
UP
UP
UP
UP
UP
UP
UP
UP
UP
UP
UP
UP
UP
UP
UP
UP
UP
UP
UP
UP
UP
UP
UP
UP
UP
UP
UP
UP
UP
UP
UP
UP
UP
UP
UP
UP
UP
UP
UP
UP
UP
UP
UP

UP
UP
UP
UP
UP
UP
UP
UP
UP
UP
UP
UP
UP
UP
UP
UP
UP
UP
UP
UP
UP
UP
UP
UP
UP
UP
UP
UP
UP
UP
UP

**Type**
DOWN
DOWN
DOWN
DOWN
DOWN
DOWN
DOWN
DOWN
DOWN
DOWN
DOWN
DOWN
DOWN

DOWN
DOWN
DOWN
DOWN
DOWN
DOWN
DOWN
DOWN
DOWN
DOWN
DOWN
DOWN
DOWN
DOWN
DOWN
DOWN
DOWN
DOWN
DOWN
DOWN
DOWN
DOWN
DOWN
DOWN
DOWN
DOWN
DOWN
DOWN
DOWN
DOWN
DOWN
DOWN
DOWN
DOWN
DOWN
DOWN
DOWN
DOWN
DOWN
DOWN
DOWN
DOWN
DOWN
DOWN
DOWN
DOWN

DOWN
DOWN
DOWN
DOWN
DOWN
DOWN
DOWN
DOWN
DOWN
DOWN
DOWN
DOWN
DOWN
DOWN
DOWN
DOWN
DOWN
DOWN
DOWN
DOWN
DOWN
DOWN
DOWN
DOWN
DOWN
DOWN
DOWN
DOWN
DOWN
DOWN
DOWN
DOWN
DOWN
DOWN
DOWN
DOWN
DOWN
DOWN
DOWN
DOWN
DOWN
DOWN
DOWN
DOWN
DOWN
DOWN

DOWN
DOWN
DOWN
DOWN
DOWN
DOWN
DOWN
DOWN
DOWN
DOWN
DOWN
DOWN
DOWN
DOWN
DOWN
DOWN
DOWN
DOWN
DOWN
DOWN
DOWN
DOWN
DOWN
DOWN
DOWN
DOWN
DOWN
DOWN
DOWN
DOWN
DOWN
DOWN
DOWN
DOWN
DOWN
DOWN
DOWN
DOWN
DOWN
DOWN
DOWN
DOWN
DOWN
DOWN
DOWN
DOWN

DOWN
DOWN
DOWN
DOWN
DOWN
DOWN
DOWN
DOWN
DOWN
DOWN
DOWN
DOWN
DOWN
DOWN
DOWN
DOWN
DOWN
DOWN
DOWN
DOWN
DOWN
DOWN
DOWN
DOWN
DOWN
DOWN
DOWN
DOWN
DOWN
DOWN
DOWN
DOWN
DOWN
DOWN
DOWN
DOWN
DOWN
DOWN
DOWN
DOWN
DOWN
DOWN
DOWN
DOWN
DOWN
DOWN

DOWN
DOWN
DOWN
DOWN
DOWN
DOWN
DOWN
DOWN
DOWN
DOWN
DOWN
DOWN
DOWN
DOWN
DOWN
DOWN
DOWN
DOWN
DOWN
DOWN
DOWN
DOWN
DOWN
DOWN
DOWN
DOWN
DOWN
DOWN
DOWN
DOWN
DOWN
DOWN
DOWN
DOWN
DOWN
DOWN
DOWN
DOWN
DOWN
DOWN
DOWN
DOWN
DOWN
DOWN
DOWN
DOWN

DOWN
DOWN
DOWN
DOWN
DOWN
DOWN
DOWN
DOWN
DOWN
DOWN
DOWN
DOWN
DOWN
DOWN
DOWN
DOWN
DOWN
DOWN
DOWN
DOWN
DOWN
DOWN
DOWN
DOWN
DOWN
DOWN
DOWN
DOWN
DOWN
DOWN
DOWN
DOWN
DOWN
DOWN
DOWN
DOWN
DOWN
DOWN
DOWN
DOWN
DOWN
DOWN
DOWN
DOWN
DOWN
DOWN

DOWN
DOWN
DOWN
DOWN
DOWN
DOWN
DOWN
DOWN
DOWN
DOWN
DOWN
DOWN
DOWN
DOWN
DOWN
DOWN
DOWN
DOWN
DOWN
DOWN
DOWN
DOWN
DOWN
DOWN
DOWN
DOWN
DOWN
DOWN
DOWN
DOWN
DOWN
DOWN
DOWN
DOWN
DOWN
DOWN
DOWN
DOWN
DOWN
DOWN
DOWN
DOWN
DOWN
DOWN
DOWN
DOWN

DOWN
DOWN
DOWN
DOWN
DOWN
DOWN
DOWN
DOWN
DOWN
DOWN
DOWN
DOWN
DOWN
DOWN
DOWN
DOWN
DOWN
DOWN
DOWN
DOWN
DOWN
DOWN
DOWN
DOWN
DOWN
DOWN
DOWN
DOWN
DOWN
DOWN
DOWN
DOWN
DOWN
DOWN
DOWN
DOWN
DOWN
DOWN
DOWN
DOWN
DOWN
DOWN
DOWN
DOWN
DOWN
DOWN

DOWN
DOWN
DOWN
DOWN
DOWN
DOWN
DOWN
DOWN
DOWN
DOWN
DOWN
DOWN
DOWN
DOWN
DOWN
DOWN
DOWN
DOWN
DOWN
DOWN
DOWN
DOWN
DOWN
DOWN
DOWN
DOWN
DOWN
DOWN
DOWN
DOWN
DOWN
DOWN
DOWN
DOWN
DOWN
DOWN
DOWN
DOWN
DOWN
DOWN
DOWN
DOWN
DOWN
DOWN
DOWN
DOWN

DOWN
DOWN
DOWN
DOWN
DOWN
DOWN
DOWN
DOWN
DOWN
DOWN
DOWN
DOWN
DOWN
DOWN
DOWN
DOWN
DOWN
DOWN
DOWN
DOWN
DOWN
DOWN
DOWN
DOWN
DOWN
DOWN
DOWN
DOWN
DOWN
DOWN
DOWN
DOWN
DOWN
DOWN
DOWN
DOWN
DOWN
DOWN
DOWN
DOWN
DOWN
DOWN
DOWN
DOWN
DOWN
DOWN

DOWN
DOWN
DOWN
DOWN
DOWN
DOWN
DOWN
DOWN
DOWN
DOWN
DOWN
DOWN
DOWN
DOWN
DOWN
DOWN
DOWN
DOWN
DOWN
DOWN
DOWN
DOWN
DOWN
DOWN
DOWN
DOWN
DOWN
DOWN
DOWN
DOWN
DOWN
DOWN
DOWN
DOWN
DOWN
DOWN
DOWN
DOWN
DOWN
DOWN
DOWN
DOWN
DOWN
DOWN
DOWN
DOWN

DOWN
DOWN
DOWN
DOWN
DOWN
DOWN
DOWN
DOWN
DOWN
DOWN
DOWN
DOWN
DOWN
DOWN
DOWN
DOWN
DOWN
DOWN
DOWN
DOWN
DOWN
DOWN
DOWN
DOWN
DOWN
DOWN
DOWN
DOWN
DOWN
DOWN
DOWN
DOWN
DOWN
DOWN
DOWN
DOWN
DOWN
DOWN
DOWN
DOWN
DOWN
DOWN
DOWN
DOWN
DOWN
DOWN

DOWN
DOWN
DOWN
DOWN
DOWN
DOWN
DOWN
DOWN
DOWN
DOWN
DOWN
DOWN
DOWN
DOWN
DOWN
DOWN
DOWN
DOWN
DOWN
DOWN
DOWN
DOWN
DOWN
DOWN
DOWN
DOWN
DOWN
DOWN
DOWN
DOWN
DOWN
DOWN
DOWN
DOWN
DOWN
DOWN
DOWN
DOWN
DOWN
DOWN
DOWN
DOWN
DOWN
DOWN
DOWN
DOWN

DOWN
DOWN
DOWN
DOWN
DOWN
DOWN
DOWN
DOWN
DOWN
DOWN
DOWN
DOWN
DOWN
DOWN
DOWN
DOWN
DOWN
DOWN
DOWN
DOWN
DOWN
DOWN
DOWN
DOWN
DOWN
DOWN
DOWN
DOWN
DOWN
DOWN
DOWN
DOWN
DOWN
DOWN
DOWN
DOWN
DOWN
DOWN
DOWN
DOWN
DOWN
DOWN
DOWN
DOWN
DOWN
DOWN

DOWN
DOWN
DOWN
DOWN
DOWN
DOWN
DOWN
DOWN
DOWN
DOWN
DOWN
DOWN
DOWN
DOWN
DOWN
DOWN
DOWN
DOWN
DOWN
DOWN
DOWN
DOWN
DOWN
DOWN
DOWN
DOWN
DOWN
DOWN
DOWN
DOWN
DOWN
DOWN
DOWN
DOWN
DOWN
DOWN
DOWN
DOWN
DOWN
DOWN
DOWN
DOWN
DOWN
DOWN
DOWN
DOWN

DOWN
DOWN
DOWN
DOWN
DOWN
DOWN
DOWN
DOWN
DOWN
DOWN
DOWN
DOWN
DOWN
DOWN
DOWN
DOWN
DOWN
DOWN
DOWN
DOWN
DOWN
DOWN
DOWN
DOWN
DOWN
DOWN
DOWN
DOWN
DOWN
DOWN
DOWN
DOWN
DOWN
DOWN
DOWN
DOWN
DOWN
DOWN
DOWN
DOWN
DOWN
DOWN
DOWN
DOWN
DOWN
DOWN

DOWN
DOWN
DOWN
DOWN
DOWN
DOWN
DOWN
DOWN
DOWN
DOWN
DOWN
DOWN
DOWN
DOWN
DOWN
DOWN
DOWN
DOWN
DOWN
DOWN
DOWN
DOWN
DOWN
DOWN
DOWN
DOWN
DOWN
DOWN
DOWN
DOWN
DOWN
DOWN
DOWN
DOWN
DOWN
DOWN
DOWN
DOWN
DOWN
DOWN
DOWN
DOWN
DOWN
DOWN
DOWN
DOWN

DOWN
DOWN
DOWN
DOWN
DOWN
DOWN
DOWN
DOWN
DOWN
DOWN
DOWN
DOWN
DOWN
DOWN
DOWN
DOWN
DOWN
DOWN
DOWN
DOWN
DOWN
DOWN
DOWN
DOWN
DOWN
DOWN
DOWN
DOWN
DOWN
DOWN
DOWN
DOWN
DOWN
DOWN
DOWN
DOWN
DOWN
DOWN
DOWN
DOWN
DOWN
DOWN
DOWN
DOWN
DOWN
DOWN

DOWN
DOWN
DOWN
DOWN
DOWN
DOWN
DOWN
DOWN
DOWN
DOWN
DOWN
DOWN
DOWN
DOWN
DOWN
DOWN
DOWN
DOWN
DOWN
DOWN
DOWN
DOWN
DOWN
DOWN
DOWN
DOWN
DOWN
DOWN
DOWN
DOWN
DOWN
DOWN
DOWN
DOWN
DOWN
DOWN
DOWN
DOWN
DOWN
DOWN
DOWN
DOWN
DOWN
DOWN
DOWN
DOWN

DOWN
DOWN
DOWN
DOWN
DOWN
DOWN
DOWN
DOWN
DOWN
DOWN
DOWN
DOWN
DOWN
DOWN
DOWN
DOWN
DOWN
DOWN
DOWN
DOWN
DOWN
DOWN
DOWN
DOWN
DOWN
DOWN
DOWN
DOWN
DOWN
DOWN
DOWN
DOWN
DOWN
DOWN
DOWN
DOWN
DOWN
DOWN
DOWN
DOWN
DOWN
DOWN
DOWN
DOWN
DOWN
DOWN

DOWN
DOWN
DOWN
DOWN
DOWN
DOWN
DOWN
DOWN
DOWN
DOWN
DOWN
DOWN
DOWN
DOWN
DOWN
DOWN
DOWN
DOWN
DOWN
DOWN
DOWN
DOWN
DOWN
DOWN
DOWN
DOWN
DOWN
DOWN
DOWN
DOWN
DOWN
DOWN
DOWN
DOWN
DOWN
DOWN
DOWN
DOWN
DOWN
DOWN
DOWN
DOWN
DOWN
DOWN
DOWN
DOWN

DOWN
DOWN
DOWN
DOWN
DOWN
DOWN
DOWN
DOWN
DOWN
DOWN
DOWN
DOWN
DOWN
DOWN
DOWN
DOWN
DOWN
DOWN
DOWN
DOWN
DOWN
DOWN
DOWN
DOWN
DOWN
DOWN
DOWN
DOWN
DOWN
DOWN
DOWN
DOWN
DOWN
DOWN
DOWN
DOWN
DOWN
DOWN
DOWN
DOWN
DOWN
DOWN
DOWN
DOWN
DOWN
DOWN

DOWN
DOWN
DOWN
DOWN
DOWN
DOWN
DOWN
DOWN
DOWN
DOWN
DOWN
DOWN
DOWN
DOWN
DOWN
DOWN
DOWN
DOWN
DOWN
DOWN
DOWN
DOWN
DOWN
DOWN
DOWN
DOWN
DOWN
DOWN
DOWN
DOWN
DOWN
DOWN
DOWN
DOWN
DOWN
DOWN
DOWN
DOWN
DOWN
DOWN
DOWN
DOWN
DOWN
DOWN
DOWN
DOWN

DOWN
DOWN
DOWN
DOWN
DOWN
DOWN
DOWN
DOWN
DOWN
DOWN
DOWN
DOWN
DOWN
DOWN
DOWN
DOWN
DOWN
DOWN
DOWN
DOWN
DOWN
DOWN
DOWN
DOWN
DOWN
DOWN
DOWN
DOWN
DOWN
DOWN
DOWN
DOWN
DOWN
DOWN
DOWN
DOWN
DOWN
DOWN
DOWN
DOWN
DOWN
DOWN
DOWN
DOWN
DOWN
DOWN

DOWN
DOWN
DOWN
DOWN
DOWN
DOWN
DOWN
DOWN
DOWN
DOWN
DOWN
DOWN
DOWN
DOWN
DOWN
DOWN
DOWN
DOWN
DOWN
DOWN
DOWN
DOWN
DOWN
DOWN
DOWN
DOWN
DOWN
DOWN
DOWN
DOWN
DOWN
DOWN
DOWN
DOWN
DOWN
DOWN
DOWN
DOWN
DOWN
DOWN
DOWN
DOWN
DOWN
DOWN
DOWN
DOWN

DOWN
DOWN
DOWN
DOWN
DOWN
DOWN
DOWN
DOWN
DOWN
DOWN
DOWN
DOWN
DOWN
DOWN
DOWN
DOWN
DOWN
DOWN
DOWN
DOWN
DOWN
DOWN
DOWN
